# A chemoinformatics-guided platform for efficient discovery of RNA-binding small molecules: Proof-of-concept for myotonic dystrophy type 1

**DOI:** 10.64898/2026.05.08.723748

**Authors:** Amirhossein Taghavi, Jingsong Shan, Xiyuan Yao, Patrick R. A. Zanon, Kisu Sung, Álvaro Simba-Lahuasi, Sylwia Gorlach, Henning Labuhn, David Salthouse, Zhen Wang, Adeline Feri, Matthew D. Disney

**Affiliations:** Department of Chemistry, The Herbert Wertheim UF Scripps Institute for Biomedical Innovation & Technology, 130 Scripps Way, Jupiter, FL 33458, USA; Department of Chemistry, The Scripps Research Institute, 130 Scripps Way, Jupiter, FL 33458, USA; Depixus SAS, 3-5 Impasse Reille, 75014, Paris, France

## Abstract

Structured RNAs cause human diseases but remain challenging to target selectively with small molecules. Here, we report a chemoinformatics-guided discovery framework that integrates fingerprint-based molecular design, experimental validation, and mechanistic profiling to identify small molecules that bind highly structured, disease-associated RNAs. Using an RNA-binder fingerprint derived from known ligands, a Tversky similarity screen of >8 million compounds yielded a 150-member library enriched in chemical space for RNA-active scaffolds. Target engagement and cell-based assays identified multiple selective ligands for the pathogenic expanded triplet repeat, r(CUG)^exp^, that causes myotonic dystrophy type 1 (DM1) by binding and sequestering the RNA-binding protein muscleblind-like 1 (MBNL1). Biophysical and single-molecule analyses revealed that the small molecules bind the 1×1 nucleotide U/U internal loops formed when r(CUG)^exp^ folds, partially block MBNL1 binding, and modulate RNA folding equilibria. Two optimized scaffolds rescued MBNL1-dependent splicing in patient-derived myotubes with micromolar potency and minimal cytotoxicity. This study establishes a generalizable, data-driven platform for discovering drug-like RNA-binding lead small molecules and demonstrates its application to the toxic repeat expansion RNA underlying DM1.

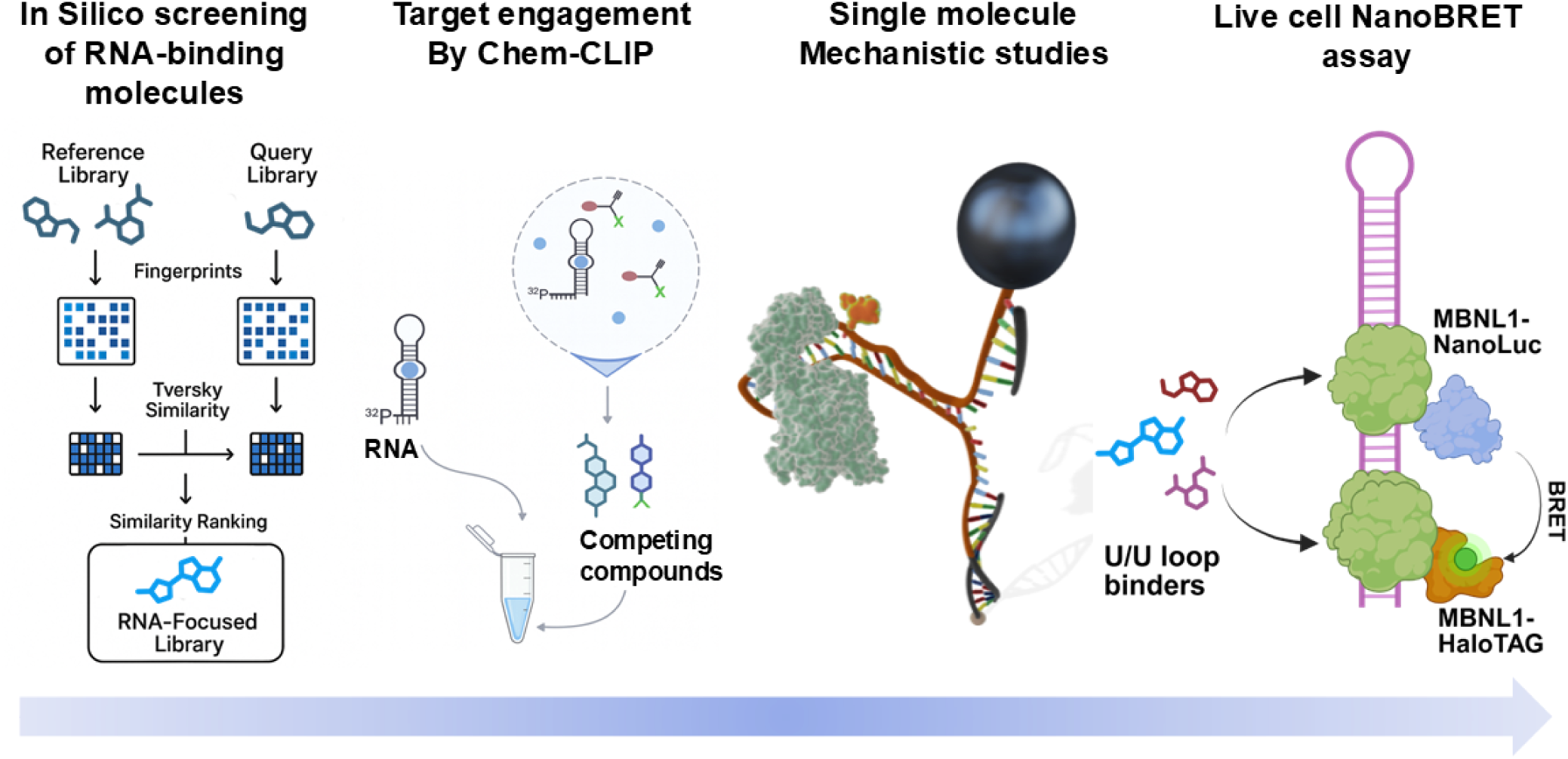
Graphical Abstract.

## INTRODUCTION

RNA structures play essential roles in gene expression, alternative pre-mRNA splicing, translation, and cellular stress responses.^1^ Disruption or aberrant formation of these structures contributes directly to human disease,^2^ yet discovering selective small-molecule ligands for RNA remains difficult. Screening hit rates are typically low, RNA-focused libraries are scarce, and most chemical space has been shaped around protein-targeting guidelines not suited for RNA. One practical approach has been the use of molecular features that recur in RNA-binding molecules. By extracting these features, large chemical libraries can be filtered toward RNA-enriched space. Early RNA-focused libraries revealed useful biases, yet they captured only a fraction of the structural diversity present across all known RNA-binding scaffolds. Furthermore, many libraries did not reflect the range of pocket geometries found in natural RNA structures.^3^

To address these gaps, we assembled a next-generation RNA-focused library by integrating molecular fingerprints from Inforna^4^ (a curated collection of 509 small molecules reported to bind RNA, compiled from both published literature and internally curated datasets) with similarity searching,^5^ shape-based clustering,^6^ and scaffold diversity analysis.^7^ The resulting library was then applied to the expanded r(CUG) repeat [r(CUG)^exp^] harbored in the 3’ untranslated region (UTR) of the dystrophia myotonic protein kinase (*DMPK*) mRNA. In healthy individuals, the number of r(CUG)_n_ repeats is 5-37 while individuals with DM1 have 50-2000 repeats.^8^ The expanded RNA folds into a hairpin structure with a periodic array of 1×1 nucleotide U/U loops that sequester proteins such as muscleblind-like protein 1 (MBNL1), a regulator of alternative pre-mRNA splicing. The protein’s sequestration results in alternative pre-mRNA splicing defects that cause disease.^9^ Moreover, MBNL1 and other RNA-binding proteins bound to r(CUG)^exp^ form nuclear foci, reducing nucleocytoplasmic transport and translation of *DMPK* mRNA.^10^ These biological data have informed various potential therapeutic approaches, including prevention or disruption of the complex formation between r(CUG)^exp^ and the MBNL1^11, 12^ and allele-selective elimination of the mutant RNA.^13^

In this work, chemoinformatics, a competitive target engagement assay that relies on covalent chemistry dubbed Chemical Cross-Linking and Isolation by Pull-down (Chem-CLIP),^14^ biophysical measurements, single-molecule magnetic force spectroscopy, and cell-based assays were combined into a unified workflow to identify starting ligands that bind to r(CUG)^exp^. Ligands that recognize the 1×1 nucleotide U/U internal loops and inhibit the r(CUG)^exp^–MBNL1 interaction were identified, optimized, and validated. This platform, in principle, is suitable for broad application to structured RNAs.

Despite selective recognition of r(CUG)^exp^ and bioactivity, the initial hit was limited by its solubility, a common issue for ligands that target RNA.^15^ To computationally address this limitation, a hit-expansion and optimization workflow integrating positional analogue scanning (PAS),^16^ solubility modeling, and functional validation was employed. Starting from the most potent bioactive lead compound, PAS was used to systematically reposition key polar functional groups to improve aqueous solubility, guided by computational logS predictions.^17^ The most soluble analogue then served as a template for molecular fingerprint-based similarity searching, yielding 25 structurally related compounds. Functional screening of the structurally related compounds using a live-cell NanoBioluminescence Resonance Energy Transfer (NanoBRET) assay^18^ identified five small molecules that significantly disrupted the r(CUG)^exp^-MBNL1 interaction in live cells with no detectable cytotoxicity. These results underscore the effectiveness of the PAS–logS–similarity strategy. Nuclear magnetic resonance (NMR) spectroscopy confirmed that the most potent compound in the NanoBRET assay binds selectively to the 1×1 nucleotide U/U internal loops within r(CUG)^exp^, inducing local structural perturbations while sparing canonically base-paired regions.

Together, these results highlight a generalizable, data-driven approach for optimizing RNA-binding small molecules, yielding more soluble, potent, and selective inhibitors of toxic RNA–protein complexes implicated in DM1. As with most lead optimization efforts—particularly for RNA-targeted initial hits—medicinal chemistry must be guided to identify follow-up compounds with improved properties. Augmenting early computational hit analysis to address solubility explicitly, as described here, enables more efficient optimization while minimizing suboptimal physicochemical properties in downstream compounds.

## RESULTS

### Design of a library of lead-like compounds targeting RNA

Various *in silico* screening methods have been validated to explore large, chemical databases to identify desired small molecules.^19, 20^ In this study, a structure-based similarity screening method was employed to identify drug-like chemical matter with a propensity to bind RNA (**Figure 1A**). The aim was to identify compounds that are most like an established query molecule(s), quantified as “similarity” or “degree of resemblance”.^21^ Various molecular descriptors, whether structural or physicochemical features, can be used to represent the small molecules. Molecular fingerprints, which store the information content of small molecules in a sequence of bits, are the most popular tool for similarity searching.^22^ These bit vectors are calculated for a set of query molecules as well as for a database of small molecules, which are compared in a pairwise manner to identify new molecules. Although two dimensional (2D) fingerprints do not contain conformational information as 3D fingerprints do, they have been very successful in many applications.^23^

**Figure 1.**
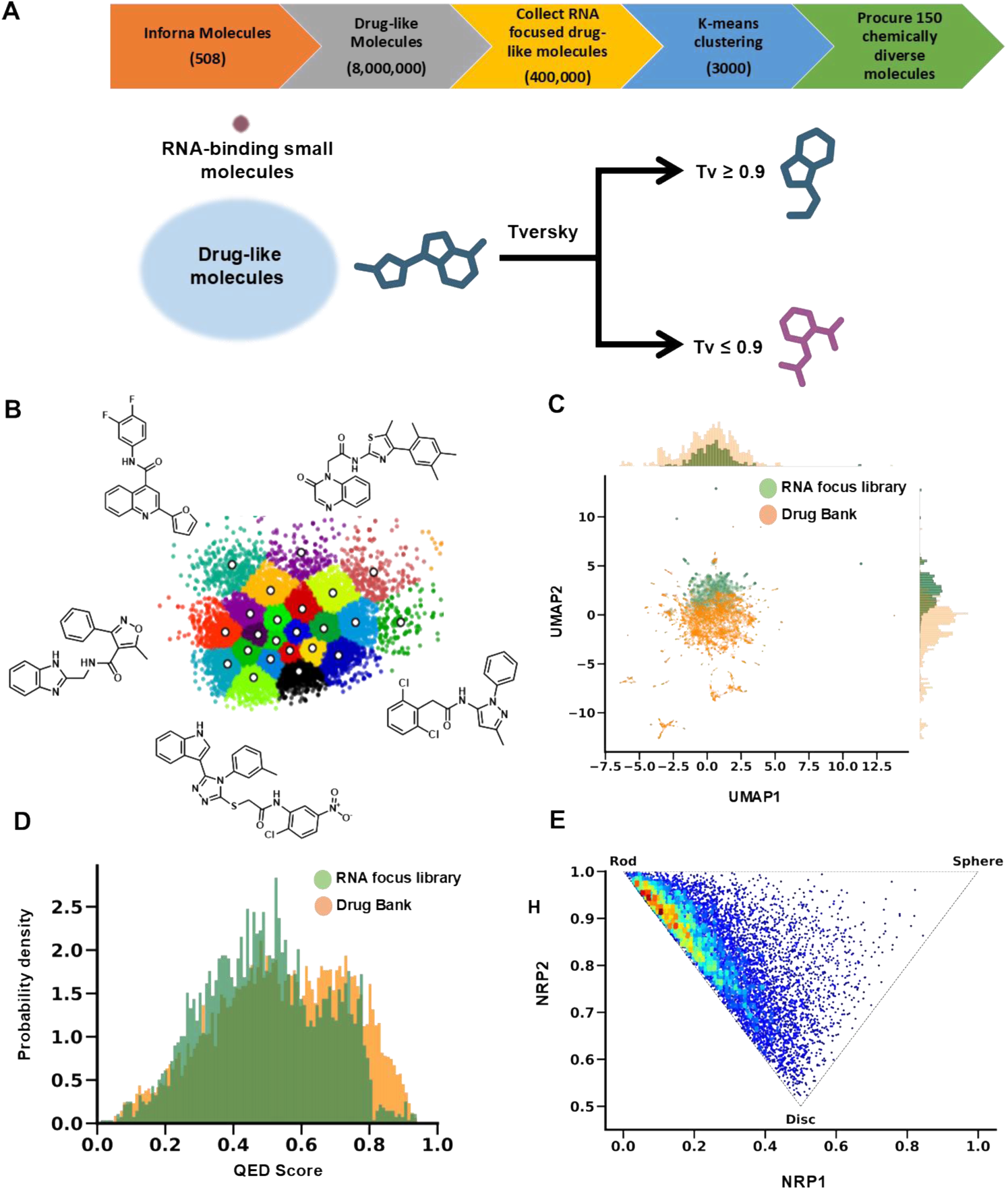
Exploring molecular features of known RNA binders to inform the design of an RNA-specific, drug-like small molecule library. (A) Pipeline to create an RNA-focused library by extracting the molecular features of all known small molecule RNA binders; screening of the molecular features in known RNA binders against a drug-like screening library with 8,000,000 members and identification of similar molecules using the Tversky (Tv) index. Molecules with a Tv index **≥** 0.9 are considered similar. (B) Two-step K-means clustering to group similar molecules (clustering). The center of each cluster was selected as a representative to create a diverse set of compounds (n = 150). (C) Overlaying the chemical space of the RNA-focused library with small molecules in DrugBank^93^ using the UMAP^32^ shows the degree of overlap between the two libraries. (D) Calculation of quantitative estimate of drug-likeness (QED) for the RNA-focused library and DrugBank. The degree of overlap indicates the drug-like properties of RNA-binding small molecules. (E) Principal moment of inertia (PMI) plot of the RNA-focused library explaining the shape distribution (rod, sphere or disc) of small molecules.

Here, “atom pairs” (AP) circular fingerprints were chosen, where atom pairs are encoded using atomic invariants in combination with their bond distances to assess the shortest topological path between two atoms.^24^ This method addresses some of the limitations associated with other fingerprints including ECFP4 and Morgan fingerprints,^25,^ which struggle to distinguish structural differences in large molecules and provide limited information on global features like size and shape.^26^ To perform virtual screening based on fingerprint similarity, an active molecule or set of active molecules representing the query is required to probe for similarities with a molecular database, preferably enriched in drug-like molecules. Inforna,^4^ a collection of RNA small molecule binders (n = 504), was selected as the query library and a collection of 8,000,000 in-stock compounds from Chemspace was selected as the molecular database (**Figure 1A**). The Tversky index (Tv),^27^ which incorporates an asymmetry element that gives more importance to the features of the reference molecule, allowing the similarity measure to evaluate how well a candidate compound preserves the key structural features of the reference ligand, was used to quantify similarity; thus Tv is more discriminatory than the traditionally used Tanimoto coefficient (Tc).^28^ Moreover, by implementing different weightings of the bit strings being compared, Tv is the most successful and least failure-prone among nine other similarity scores, as reported by ChEMBL.^29^ A cutoff of >0.9 was used to select for molecules with chemical properties similar to the Inforna dataset (**Figure 1A**). This threshold ensures that the resulting library retains the key molecular characteristics associated with RNA recognition. Importantly, the Tversky index constrains feature similarity rather than scaffold identity, meaning that compounds can share physicochemical and substructural properties while still possessing distinct core frameworks. Consistent with this, scaffold analysis of the resulting 400,000-compound library revealed extensive chemotype diversity, with a large number of unique Bemis–Murcko scaffolds present across the dataset (**Table S1**).

A two-step K-means clustering^30^ algorithm was used to preserve scaffold and chemical diversity throughout the selection process (**Figure 1B**). Scaffolds were defined using Bemis–Murcko^7^ rules, in which each compound is reduced to its core ring systems and linkers. By this definition, the parent RNA-focused library (n = 400,000) contains 133,797 unique scaffolds (**Table S1**). In the first clustering step, a randomly selected subset of 50,000 compounds was used to generate initial cluster centroids. In the second step, the Euclidean distance between the molecular fingerprints of each compound in the full 400,000-member library and the initial centroids was calculated, and each molecule was assigned to its nearest centroid to finalize cluster membership. A representative set of 3,000 structurally diverse compounds, located at the geometric centers of these clusters (**Figure 1B**), was then selected. This 3,000-compound working set retains 2,391 unique Bemis–Murcko scaffolds, preserving ∼80% of the scaffold diversity present in the parent library (**Table S2**).

Physicochemical properties of the selected compounds were calculated using RDKit^31^ and were consistent with features enriched among known RNA-binding small molecules, with average values of: molecular weight (MW) = 375 ± 7 Da; calculated octanol/water partition coefficient (cLogP) = 4.04 ± 0.09; topological polar surface area (TPSA) = 70.2 ± 1.9 Å²; hydrogen bond donors (HBD) = 1.0 ± 0.06; hydrogen bond acceptors (HBA) = 4.65 ± 0.15; rotatable bonds = 4.47 ± 0.16; aromatic rings = 3.54 ± 0.06; and fraction sp^3^ hybridized carbon (Csp^3^) = 0.14 ± 0.01. These property distributions align with those reported for known RNA-binding ligands and also provide indirect insight into aqueous solubility. In particular, the moderate molecular weight, balanced lipophilicity (cLogP ≈ 4), and presence of polar surface area and hydrogen-bonding functionality suggest that the compounds maintain sufficient polarity to support aqueous tractability while retaining aromatic character important for RNA recognition. Collectively, these features indicate that the library is expected to exhibit moderate but generally acceptable aqueous solubility, consistent with small molecules that balance RNA-binding interactions with practical biophysical properties.

Uniform Manifold Approximation and Projection (UMAP)^32^ analysis was used to visualize the chemical space and dimensionality of the 3,000 selected molecules (**Figure 1C**), allowing for direct comparison to small molecules in DrugBank.^33^ UMAP showed that the selected compounds form overlapping but separable distributions relative to DrugBank, indicating that our library retains drug-like character while being shifted toward physicochemical properties previously associated with RNA-binding small molecules. As many RNA binders do not follow Lipinski’s “Rule of Five”,^34^ it is important to expand the chemical space beyond traditional drug-like space.^35^ Quantitative estimate of drug-likeness (QED)^36^ was also evaluated; QED is a metric of drug-likeness scored on a scale of 0 (not drug-like) to 1 (drug-like) and is based on MW, cLogP, TPSA, numbers of HBDs and HBAs, the number of aromatic rings and rotatable bonds, and the presence of unwanted chemical functionalities (**Figure 1D**). A QED cutoff of ≥ 0.35 (corresponding to the lower bound typically associated with drug-like chemical space^37^) was applied during the final filtering step, ensuring retention of compounds with acceptable drug-likeness for downstream screening.

We further analyzed the three-dimensional (3D) shape distribution of the 400,000 molecules as well as the 3,000 molecules using the normalized principal moments of inertia ratios (NPRs)^38^ (**Figure 1E**). Molecule shape is an important descriptor as it identifies the pattern of molecular interactions with the target,^39^ and small molecules differentiate between targets via shape recognition; that is, the positioning of various functional groups in a binding pocket^40^ necessitates shape variations in the optimization process.^41^ To categorize 3D shapes, standard NPR-based geometric thresholds were used: rod-like (NPR1 < 0.3 and NPR2 > 0.7), disc-like (NPR1 > 0.7 and NPR2 < 0.3), and sphere-like (0.3 ≤ NPR1 and NPR2 ≤ 0.7). The parent library exhibited a broad distribution of 3D shapes, and the 3,000-compound subset retained a representative mix of rod-, disc-, and sphere-like geometries. Specifically, this subset consisted of approximately 55% rod-like, 35% disc-like, and 10% sphere-like structures, demonstrating deliberate preservation of 3D structural diversity rather than the rod-heavy bias typical of many screening libraries.^42^

Finally, K-means clustering was applied to this QED-filtered, shape-balanced subset to select 150 structurally diverse representatives, which were procured for experimental studies. This compound set preserved high scaffold diversity and maintained favorable QED values and balanced 3D shape composition, indicating that drug-likeness filtering and shape considerations did not erode the chemical or geometric diversity of the final library.

### Identification of r(CUG)^exp^ binders by using competitive cross-linking

This “lead-like” library of 150 small molecules was screened for binding to a validated *in vitro* model of r(CUG)^exp^, r(CUG)12,^43^ by *in vitro* competitive (C)-Chem-CLIP (**Figure 2A**). This target engagement approach relies on the formation of a covalent bond between a Chem-CLIP probe and an RNA target. The probe comprises an RNA-binding module, a cross-linking moiety such as diazirine or chlorambucil, and a chemical handle for subsequent pull-down and isolation of the RNA-small molecule adduct.^44^ In its competitive variant, an RNA is co-incubated with a Chem-CLIP probe known to engage the target and increasing concentrations of a small molecule of interest lacking a cross-linking module (**Figure 2A**). Thus, a competing small molecule reduces the amount of RNA pulled down by the Chem-CLIP probe, here radioactively labeled r(CUG)_12_. For these studies, Chem-CLIP probe **1** (**Figure 2B**), which was previously shown to bind r(CUG)^exp^ both *in vitro* and in cells,^45^ was used.

**Figure 2.**
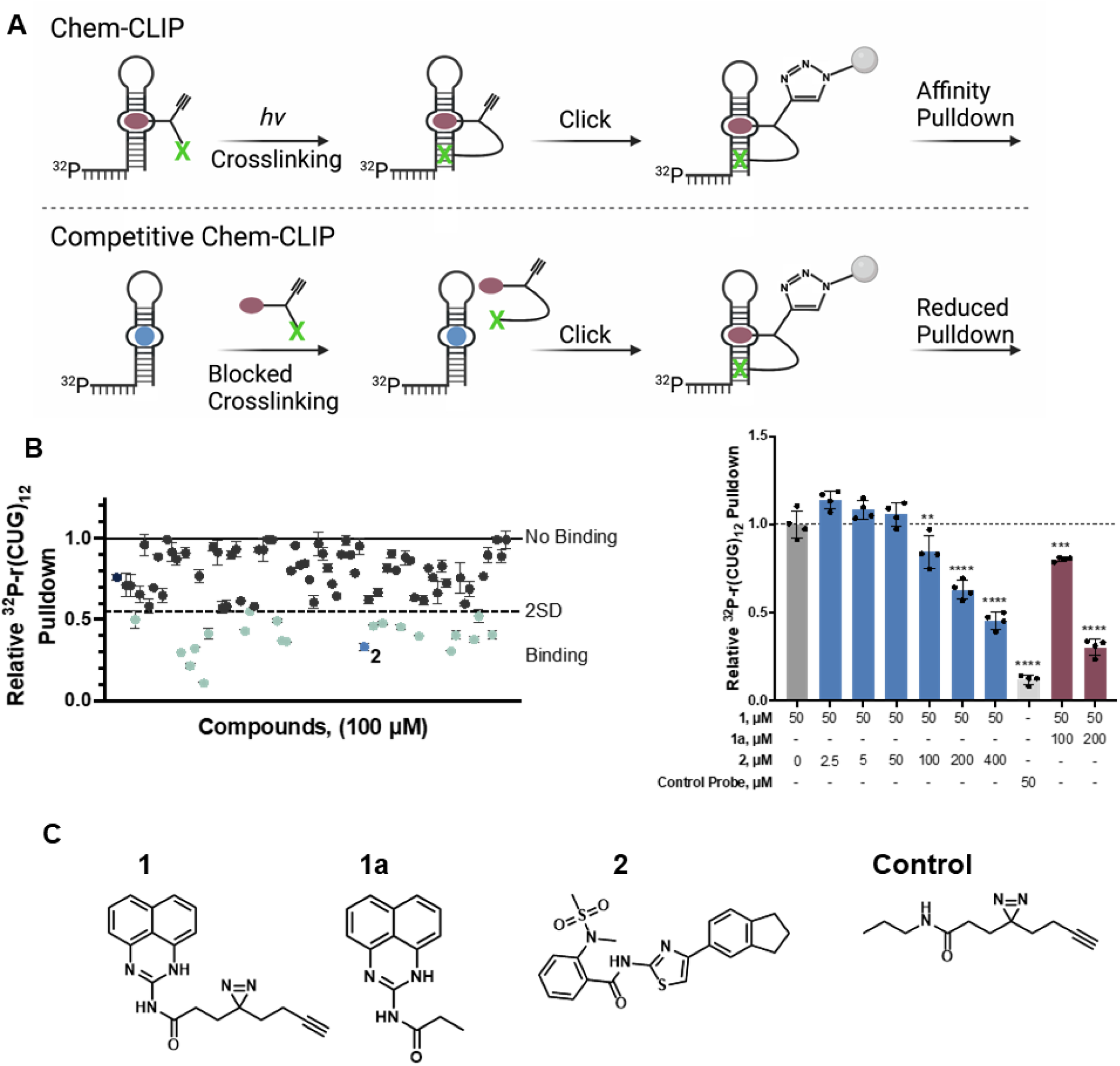
Screening of the RNA specific drug-like small molecule library by in vitro Chem-CLIP. (A) Schematic representation of the *in vitro* screening method Chemical Cross-Linking and Isolation by Pull-down (Chem-CLIP) and its competitive variant (C-Chem-CLIP) used to screen the 150-member small molecule library for binding to r(CUG) repeats. (B) (left)Relative radioactive signal upon pull-down of ^32^P-labeled r(CUG)_12_, incubated with each member of the small molecule. Compounds that reduced the amount of radioactivity pulled down by >2σ were considered preliminary hits to be further validated. The structures of these compounds can be found in **Figure S1**. (right) C-Chem-CLIP studies to demonstrate the competition between **1** and **2** for binding r(CUG)_12._ **, *p* < 0.01; ***, *p* < 0.001; ****, *p* < 0.0001; as determined by a one-way ANOVA with a Bonferroni test. Error bars represent SD. (C) Chemical structure of Chem-CLIP probe **1** used for competitive screening, **1a** the parent compounds for **1**, which was used as a positive control for competition, lead compound **2**, and the control probe that lacks an RNA-binding element

Of the 150 compounds screened in this assay, 21 compounds (100 μM) competed with **1** (50 μM) as indicated by a decrease in the relative radioactive signal (>2 standard deviations; 2σ), giving a hit rate of 14% (**Figures 2B and S1**). These hits were then verified by dose response in the same assay (25, 50, 100 μM), yielding 17 compounds with dose dependent competition profiles (**Figures 2C and S2**). Several molecules, including **7**, **9**, **12**, **13**, **18**, and **20**, produced the largest decreases in pull-down, reducing recovery to 27.8–56.3% of control at 100 µM (mean 42.5 ± 12.2% of control), corresponding to a 43.7–72.2% reduction. In contrast, compounds **3**, **6**, **8**, **10**, **11**, **15**, and **19** showed more modest effects, with pull-down values of 56.8–77.7% of control (mean 69.8 ± 7.5% of control), corresponding to only a 22.3–43.2% reduction. Thus, the first group decreased pull-down by nearly twofold more on average than the second group, consistent with stronger competition for binding to r(CUG)_₁₂_ under the assay conditions. The 17 validated molecules are chemically diverse, with only a few closely related examples. Compounds **8** and **9** share a common polycyclic lactam-like scaffold, while **13** and **19** (and to a lesser extent **11**) contain a common heterocyclic core (2-((4-oxo-3,4-dihydroquinazolin-2-yl)thio)-*N*-phenylacetamide); beyond these cases, no dominant chemotype is apparent. The identified compounds span scaffolds containing functionalities/substituents commonly found in diverse small molecule collections.

### Evaluating r(CUG)^exp^ binders for bioactivity in model and disease relevant cell lines

The 17 compounds that dose dependently competed **1** for binding *in vitro* were evaluated in a cellular luciferase reporter assay that measures alleviation of the nucleocytoplasmic transport defect of r(CUG)^exp^-containing mRNA.^46^ In brief, C2C12 mouse myoblasts stably express firefly luciferase fused to a 3′ UTR containing r(CUG)_800_. As the repeat is sequestered in nuclear foci, low amounts of luciferase are expressed. Compounds that bind to repeat expansion and displace RNA-binding proteins (RBPs) facilitate nucleocytoplasmic transport and hence increase luciferase expression (**Figure 3A**). C2C12 cells stably expressing a luciferase reporter fused to a 3′ UTR containing r(CUG)_0_ serve as a control. Of the 17 compounds, tested at concentrations that do not affect cell viability by >20% (typically up to 50 μM; **Figure S3**), seven dose dependently increased luciferase signal in r(CUG)_800_-expressing cells without affecting the signal in the control cell line (**Figures 3B, S3, and S4**).

**Figure 3.**
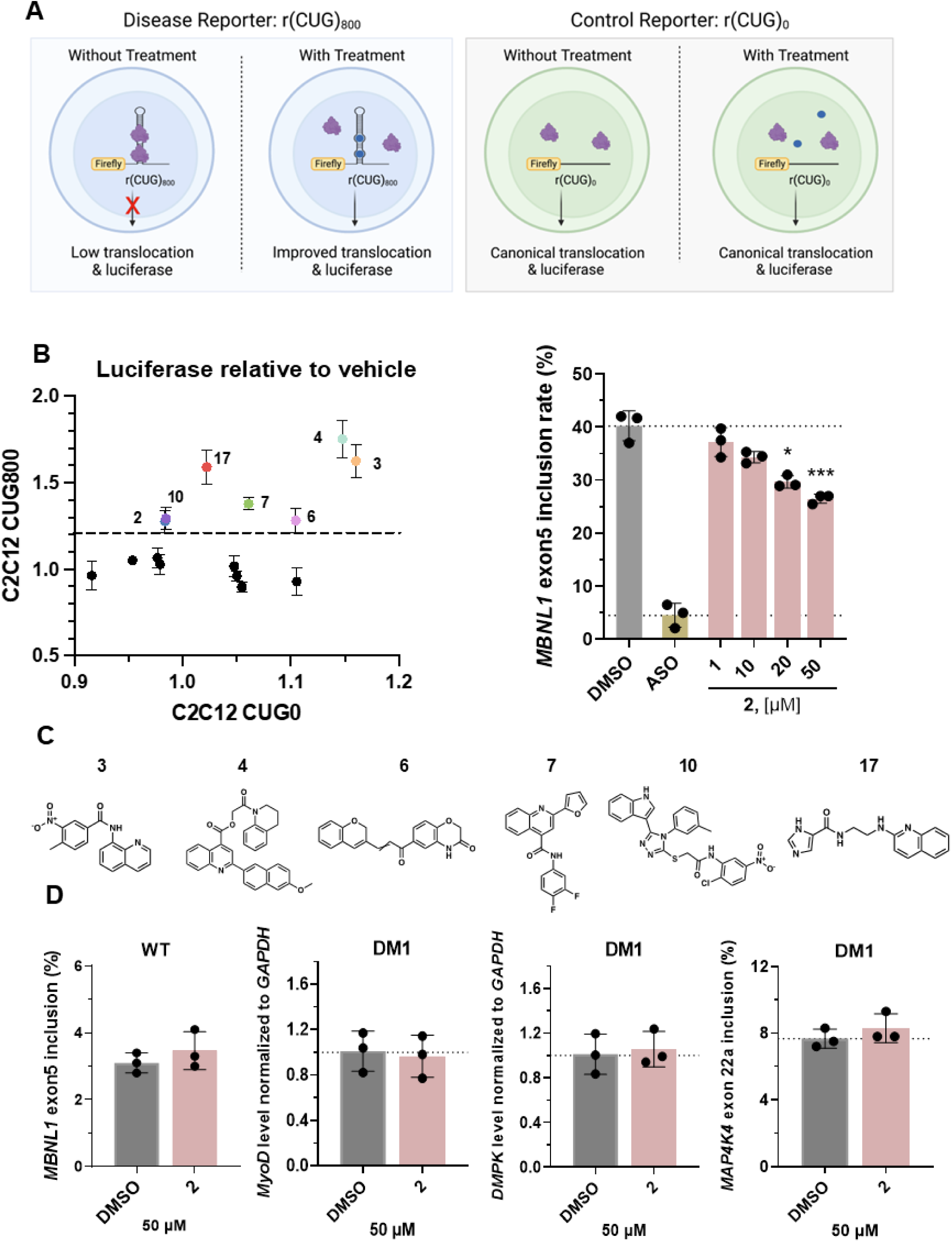
Identification of a bioactive small molecule in DM1 cellular models. (A) Schematic representation of C2C12 reporter cell lines used to identify small molecules that selectively bind r(CUG)^exp^ and alleviate defects observed in its nucleocytoplasmic transport. (Top left) MBNL1, a regulator of alternative pre-mRNA splicing, is sequestered in the nucleus by r(CUG)^exp^ and hence deactivated (loss of function). (B) (left)2D plot of the effects of small molecule treatment on luciferase signal in C2C12 stably expressing luciferase fused to r(CUG)_800_ as compared to C2C12 cells that stably express luciferase fused to r(CUG)_0_. (right) Effect of **2** on *MBNL1* exon 5 inclusion in DM1 myotubes as measured by RT-PCR (n = 3). The effect of 2.5 µM of CAG25 Vivo-Morpholino on *MBNL1* exon 5 splicing is included for comparison. (C) chemical structure of hit compounds. (D) Effect of **2** (50 μM) on *MBNL1* exon 5 inclusion in WT myotubes, on *MyoD* and *DMPK* abundance in DM1 myotubes, each assessed by RT-qPCR, and on NOVA-regulated *MAP4K4* exon 22a splicing assessed by end-point RT-PCR and fragment analyzer (n = 3 biological replicates for each measurement type). For all panels, Data are reported as mean ± standard deviation (n = 3), Statistical significance was determined by a One-way ANOVA with multiple comparisons with significance thresholds: *, p < 0.05; **, p < 0.01; ***, p < 0.001; and ****, p < 0.0001.

To verify the bioactivity observed in C2C12 assay resulting from MBNL1 displacement, we employed a previously established time-resolved fluorescence resonance energy transfer (TR-FRET) assay. In this system, RNA-Protein complex formation was monitored using bi-r(CUG)_12_ and MBNL1-His_6_. FRET signal will be detected between Streptavidin-XL665 and anti-His6 terbium-labeled antibody when the r(CUG)_12_-MBNL1 complex formed. Accordingly, small molecules that disrupt the r(CUG)_12_-MBNL1 interaction will produce a dose-dependent reduction in FRET signal. Among the 17 compounds from *in vitro* competitive Chem-CLIP, **8**, **9** and **22** shows solubility issue above 50 μM and therefore excluded for analysis. All seven bioactive hits from C2C12 r(CUG)_800_ cell show dose-dependent reduction in TR-FRET assay (**Figure S5**), with **2** (26.2 ± 3.1% inhibition at 100 μM) and **6** (26.6 ± 5.2% inhibition at 100 μM) show highest potency.The bioactivity of all seven compounds that dose dependently increase luciferase signal in r(CUG)_800_-expressing cells were next assessed in DM1 patient-derived and healthy myoblasts, specifically rescue of MBNL1-dependent pre-mRNA alternative splicing defects.^47^ DM1 patient-derived or wild type fibroblasts were forced into myogenic differentiation by doxycycline-induced expression of myoblast determination protein 1 (MYOD1)^48^ in the presence or absence of small molecule for 48 h. Following treatment, rescue of the *MBNL1* exon 5 alternative splicing defect^49^ was measured by end-point RT-PCR and quantification by fragment analyzer. Among the seven compounds that were active in both C2C12 cells and TR-FRET assay, **10** and **18** exhibited toxicity after 48h treatment (viability < 80%), and were therefore excluded from downstream analysis.

Following myogenic differentiation, the exon 5 inclusion rate in DM1 patient-derived cells was 30 ± 2%, compared with 3 ± 0.2% in wild type myoblasts (**Figure S6**). Only compound **2** rescued *MBNL1* exon 5 splicing at 50 µM with 31 ± 4% (p < 0.0001). Further experiment confirmed that **2** dose dependently rescued *MBNL1* exon 5 splicing with percent inclusion rates of = 26 ± 0.7% (p = 0.0131) and 37% ± 2% rescue (p = 0.0001) at the 20 µM and 50 µM doses, respectively (**Figures 3C, S7, and S8**). To confirm the end-point RT-PCR results, we verified these splicing changes by RT-qPCR. The *MBNL1* transcript containing exon 5 was reduced by 28 ± 6% (p = 0.0354) in the presence of 50 µM **2**. Notably, this decrease was not accompanied with a reduction in the MBNL1 exon 4-6 isoform or in the total MBNL1 transcript levels (**Figure S9**).

Additional RT-qPCR measurements verified that **2** did not alter the expression of *DMPK* or *MyoD*, ruling out off-target knockdown of these transcripts as the cause of the observed *MBNL1* exon 5 splicing rescue (**Figure 3D**). Notably, treatment with 50 μM of **2** had no effect on *MBNL1* exon 5 splicing in wild type myoblasts or on *MAP4K4* exon 22a alternative splicing, a NOVA-regulated splicing event,^50, 51^ in DM1 myoblasts (**Figures 3D and S10**).

Because **2** was the only molecule that demonstrated bioactivity in the cellular assays, it was prioritized for further investigation. **2** possesses physicochemical properties consistent with RNA recognition, including sufficient polarity and hydrogen-bonding capacity (TPSA ≈ 79 Å²; HBA/HBD = 5/1), while remaining within a drug-like property space (MW 428 Da; cLogP 3.95; no Lipinski violations; QED 0.65). The scaffold also displays moderate three-dimensional character (Fsp³ ≈ 0.24), which can improve optimization potential relative to more planar chemotypes. In addition, the sulfonamide functional group provides a polar anchor suitable for modular analog development. These features identify compound **2** as a promising starting point for further optimization (**Table S3 and Figure S11**).

### *In vitro* binding of 2 by chemical shift perturbation and Carr-Purcell-Meiboom-Gill (CPMG) relaxation dispersion nuclear magnetic resonance (NMR) spectrometry

The binding of **2** to r(CUG)_42_ was evaluated by ligand-based nuclear magnetic resonance (NMR) spectroscopy experiments, as they can detect weak to moderate affinity interactions.^53^ Binding can be assessed detecting changes in chemical shift (chemical shift perturbation; CSP) caused by differences in the magnetic environment between the bound and free states,^54^ or in the height of the signals (line broadening) related to the shortening of the relaxation time of the compound when interact with macromolecules, such as the RNA.^55^ In addition, the difference in the relaxation properties between the nuclei in small molecules and macromolecules can be exploited using the relaxion edited Carr-Purcell-Meiboom-Gill (CPMG) pulse sequence. Differences in the transverse relaxation times (T_2_), are directly dependent on the molecular rotation correlation time (τ_c_) and thus related to molecular weight. Small molecules relax slowly (long T_2_, second scale), resulting in sharp peaks, while molecules with a higher molecular weight, such as RNAs, relax quickly (short T_2_, millisecond scale),and exhibit broad signals. Transient binding of a small molecule to an RNA changes the relaxation time of the small molecule showing a signal attenuation.^56^

After the addition of r(CUG)_42_, compound **2** resonances exhibited CSP and LB, which are characteristic for changes in the chemical environment related to the binding. The most significant changes (combined LB and CSP) were observed in protons 6 (indane), 5 (thiazole), 1 and 2 (phenylmethanesulfonamide), suggesting a greater implication of these protons in the binding (**Figure S12**). In CPMG experiments, attenuation of the peaks of **2** when r(CUG)_42_ is added was consistently observed, which is related to the decrease of the T_2_ of the compound when forming a complex with the RNA. Remarkably, a greater effect on protons 6, 5, 1 and 2 was observed (**Figure S12**), which was consistent with the CSP and LB effect previously observed, and confirmed the dominant role of these protons in the binding of **2** to r(CUG)_42_.

### Saturation Transfer Difference (STD) NMR spectroscopy shows no binding of 2 to MBNL1

Saturation Transfer Difference (STD) NMR spectroscopy is a ligand-observed technique widely used to detect direct binding of small molecules to macromolecular targets.^57^ In this method, selective radiofrequency saturation is applied to the macromolecule, here MBNL1, and magnetization transfer through spin diffusion is relayed to bound ligands in close contact with the protein surface. The transferred saturation results in attenuation of ligand proton signals in the difference spectrum, providing direct evidence of binding and allowing identification of ligand protons involved in the interaction. No detectable STD signals were observed indicating that **2** does not bind to MBNL1 in solution (**Figure S13**).

### Single-molecule analysis of r(CUG)_21_–MBNL1 interactions

The dynamics of r(CUG)_21_ RNA were analyzed using MAGNA™ magnetic force spectroscopy, which enables parallel analysis of hundreds of individual RNA molecules and direct measurement of folding and unfolding behavior under applied force (**Figure 4A**). The r(CUG)_21_ was tethered between a flow-cell surface and a paramagnetic bead using DNA–RNA handles. Increasing magnetic force induced RNA unfolding, observed as discrete changes in bead position, while reducing force allowed refolding. This reversible process was repeated over multiple cycles to quantify unfolding and refolding forces and probabilities. Proteins or small molecules can be introduced to assess their effects on RNA dynamics.

**Figure 4.**
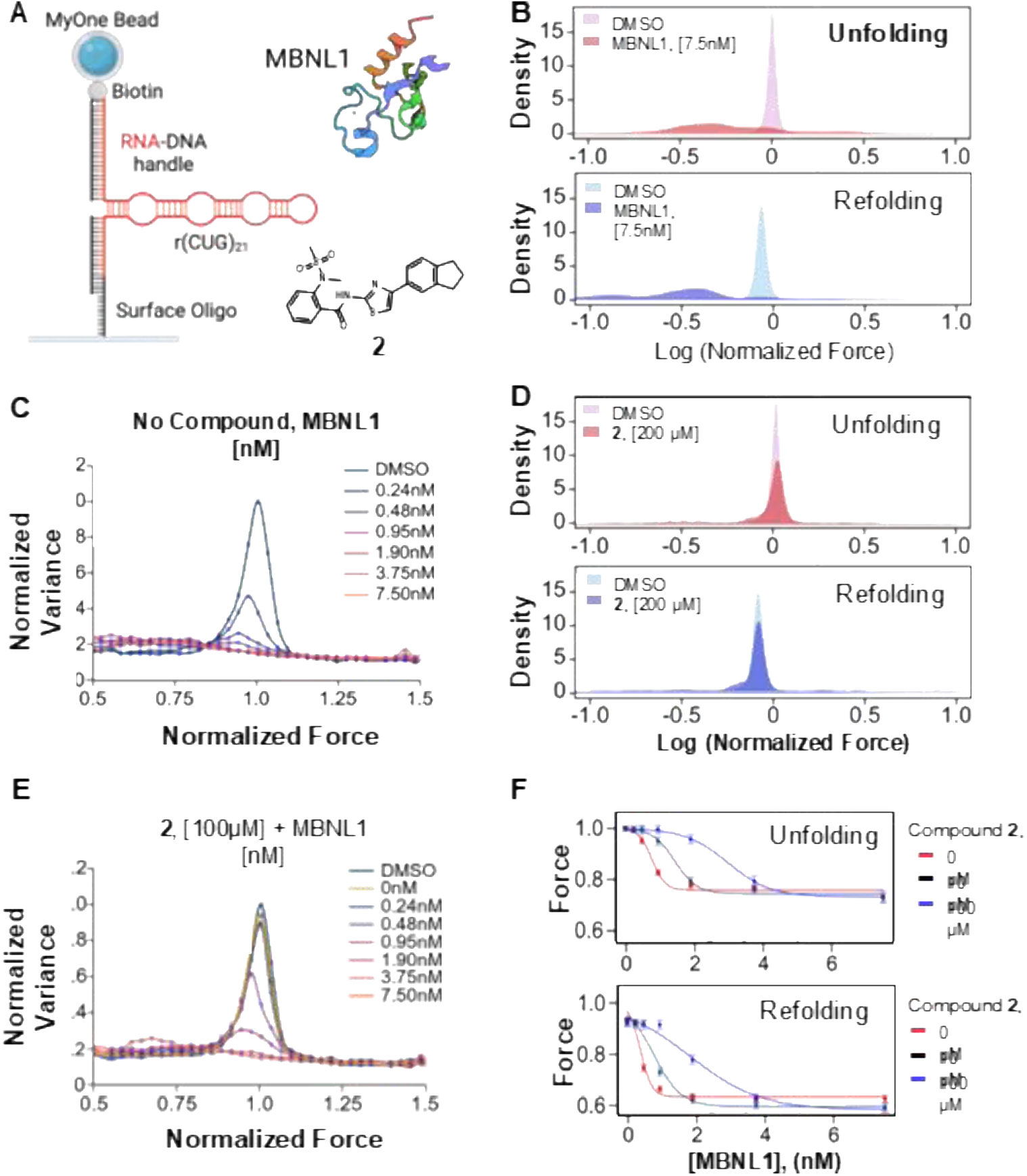
*In vitro* binding assessment of 2 to the r(CUG)^exp^ RNA by MAGNA™ magnetic force spectroscopy. (A) Schematic of the MAGNA^TM^ technology used to analyze the dynamics of the RNA’s structure. Left: The biotinylated RNA molecule is attached to a MyOne paramagnetic bead on one end and the flow cell surface on the other end. Right: The structure of MBNL1 protein (PDB:3D2N)^94^ and **2**. (B) Normalized unfolding force (upper) and refolding force (lower) distribution for all cycles, comparing the DMSO and the binding at 7.5 nM of MBNL1 (n = 255). (C) The displacement variance upon MBNL1 protein binding in constant-force experiments, in the absence of **2** (n = 153). (D) Normalized unfolding force (left) and refolding force (right) distribution for all cycles, comparing the DMSO and the binding at 100 µM of **2** (n = 249). (E) The displacement variance upon MBNL1 protein binding in step-wise constant-force experiments, in the presence of 100 µM of **2** (n =1 42). (F) The median unfolding force (left) and refolding force (right) with increasing concentrations of MBNL1 protein in the presence of 0, 10 and 100 µM of **2**, using ramp experiments (n = 255, 198, 148 for 0, 10, 100 µM of **2**, respectively). Data are median of different molecules with standard error of the mean (SEM)

Force-ramp experiments showed that binding of MBNL1 reduced the forces required for both unfolding and refolding of r(CUG)_21_ compared to RNA alone (**Figure 4B**), indicating destabilization of the repeat structure. Analysis of individual molecule trajectories revealed that MBNL1 prevented unfolding or refolding during a fraction of cycles, resulting in decreased folding and unfolding probabilities. This effect increased with MBNL1 concentration (**Figure S14B–C**). These observations indicate that MBNL1 preferentially binds r(CUG)_21_ when the RNA is unfolded (single stranded), consistent with prior studies.^12^ The reduced unfolding force suggests that MBNL1 remains associated with the RNA, leading to partial structural constraint and facilitating cooperative binding in subsequent cycles.^58^

To further characterize RNA dynamics, constant-force experiments were performed. At forces below the unfolding threshold, r(CUG)_21_ transitioned between folded and unfolded states, with equal occupancy at the equilibrium force (**Figure S14A**). RNA dynamics were quantified by calculating the variance in bead displacement at each force step. As force increased, displacement variance rose progressively until a maximum at the equilibrium force, then decreased as the RNA became fully unfolded (**Figures 5D and S14D**). In the presence of MBNL1, the maximum displacement variance was reduced and shifted slightly to lower forces, suggesting that protein binding prevents complete refolding and further destabilizes the r(CUG)_21_ structure. At the highest MBNL1 concentrations, displacement variance was nearly abolished, consistent with saturation of r(CUG)_21_ binding sites (**Figure 4C**). Taken together, these results demonstrate that MBNL1 preferentially binds single-stranded r(CUG) repeats and destabilizes the folded repeat architecture, thereby altering the folding landscape and dynamics of the pathogenic RNA structure.

**Figure 5.**
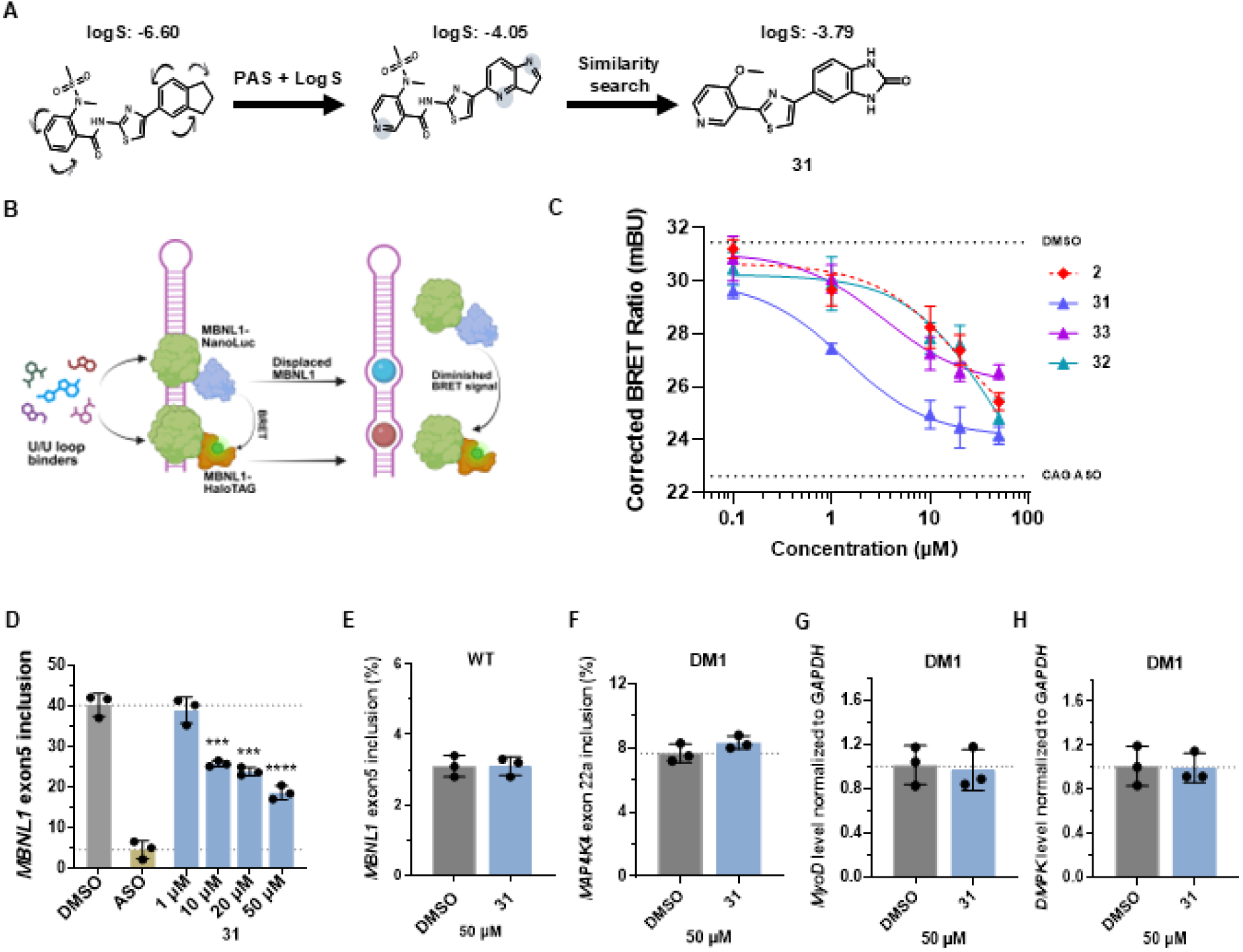
Screening of a lead optimized small molecule library for inhibition of r(CUG)^exp^−MBNL1 complex in live cells. (A) Schematic of the PAS-logS-similarity workflow. (B) Schematic of r(CUG)^exp^ forming complexes with MBNL1 and targeting the complex with small molecules. The binding of MBNL1 proteins tagged with NanoLuc (blue) and HaloTag (orange) to r(CUG)^exp^ generates a bioluminescence resonance energy transfer (BRET) signal. Upon treatment with small molecules that bind to r(CUG)^exp^ (light blue and brown), the toxic r(CUG)^exp^−MBNL1 complex is disrupted, reducing the BRET signal. (C) Dose dependent reduction of the corrected BRET ratios (mBU) for small molecules **2**, **31**, **32**, and **33**, identified from the primary screening assay. Corrected BRET ratios for vehicle-treated and cells treated with 10 μM of CAG25 Vivo-Morpholino ASO are included for comparison. Data are reported as mean ± standard deviation (n = 3 biological replicates). (D) Dose-response analysis **3** in DM1 myotubes. Cells were treated with increasing concentrations (1– 50 μM) of the compound of interest during differentiation, and exon 5 inclusion levels were measured by end-point RT-PCR. Results for compound **3** (previously shown in Figure 3C) and cells treated with 2.5 µM of CAG25 Vivo-Morpholino ASO are included for comparison (E) Effect of **3** (50 mM) on *MBNL1* exon 5 inclusion in WT myotubes, on *MyoD* and *DMPK* abundance in DM1 myotubes, each assessed by RT-qPCR, and on NOVA-regulated *MAP4K4* exon 22a splicing at, as assessed by end-point RT-PCR and fragment analyzer (n = 3 biological replicates for each measurement type). Statistical significance was determined relative to untreated controls using a two-tailed unpaired Student’s t test with significance thresholds: *, p < 0.05; **, p < 0.01; ***, p < 0.001; ****, p < 0.0001.

The molecular dynamics of r(CUG)_21_ were next examined in the presence of a constant concentration of **2** and increasing concentrations of MBNL1 protein (0.24–7.5 nM) using the same single-molecule force-ramp and constant-force assays. In force-ramp experiments, compound **2** alone (0.1–100 µM) did not significantly alter RNA unfolding or refolding forces compared to DMSO, indicating that **2** does not measurably affect r(CUG)_21_ structural stability on its own (**Figures 4D and S14E**). In contrast, in the presence of MBNL1, addition of **2** attenuated the decrease in unfolding and refolding forces caused by MBNL1 binding. This effect was most pronounced at 100 µM of **2** and intermediate at 10 µM (**Figures 4E–F**), indicating that **2** partially inhibits MBNL1–RNA interactions. Consistent results were obtained in constant-force experiments. Compound **2** alone (0.1–100 µM) did not significantly affect displacement variance or equilibrium force, confirming that it does not alter intrinsic RNA dynamics. However, in the presence of MBNL1, **2** (100 µM) delayed the MBNL1-induced reduction in maximum displacement variance and the shift to lower unfolding and refolding forces (**Figures 4E–F and S14F**), further supporting partial inhibition of MBNL1 binding.

Taken together, these data are consistent with structural and biophysical studies showing that MBNL1 binds single-stranded RNA and shifts the r(CUG)_21_ folding equilibrium toward the unfolded state.^59, 60^ The single-molecule measurements support a mechanism in which **2** engages folded r(CUG) repeats and partially prevents MBNL1 binding, thereby opposing MBNL1-induced RNA destabilization.

### Hit expansion and solubility optimization by positional analogue scanning and similarity searching

During secondary characterization of hit compound **2**, we observed that despite promising RNA-binding activity, its aqueous solubility was limited, a common challenge in RNA-targeted small-molecule discovery; chemotypes that recognize structured RNA are often enriched in aromatic and heterocyclic features leading to modest solubility under physiological conditions. Although solubility-related descriptors (e.g., TPSA, cLogP) were included among the physicochemical criteria used to select the initial screening set, experimental solubility can deviate from predicted values, and **2** was predicted to lie near the lower boundary of acceptable logS values.^17^

Given the need for adequate solubility for downstream biochemical and cellular studies, improving solubility was prioritized during hit expansion. To achieve this, positional analogue scanning (PAS)^16^ was applied in combination with computational logS^61^ (aqueous solubility) predictions. PAS systematically relocates polar or ionizable functional groups (e.g., hydroxyl, amine, or carboxyl groups) across the core scaffold, enabling identification of analogues with improved aqueous solubility while preserving RNA-binding potential. Starting from hit **2**, PAS was used to introduce and reposition polar substituents on the scaffold, and the resulting analogues were evaluated using predicted logS values to prioritize candidates with improved aqueous behavior. This process yielded **compound 2-1**, which retained the key aromatic scaffold of compound **2** but incorporated a strategically positioned polar substituent predicted to increase aqueous solubility while maintaining RNA-binding features. The structure of compound **2-1**, along with the parent scaffold and related analogues, is shown in **Figure 5A**.

This PAS-optimized, more soluble analog, compound **2-1**, was then used as the seed for a molecular fingerprint–based similarity search, which produced a focused set of 35 structurally related compounds designed to retain the core pharmacophore of **2** while biasing the series toward enhanced solubility (**Figure S15**). Compound **31** emerged from this 35-member similarity library, identified using a live-cell r(CUG)exp–MBNL1 NanoBRET assay as described in the following sections. Importantly, compound **31** displayed substantially improved predicted aqueous solubility (logS = −3.79), corresponding to an approximately 600-fold increase relative to the parent compound **2** (logS = −6.60) (**Figure 5A**).

### Identification of improved analogues using a live-cell r(CUG)^exp^–MBNL1 NanoBRET assay

The panel of 35 PAS-generated analogs was evaluated using a previously developed live-cell NanoBRET assay^18^ that quantitatively reports on formation of the r(CUG)^exp^–MBNL1 complex (**Figure 5B**). In this assay, r(CUG)^exp^ scaffolds the binding of NanoLuc-MBNL1 and HaloTag-MBNL1 fusions to generate a BRET signal; disruption of the complex by a small molecule decreases the BRET signal. Compound **2** disrupted the r(CUG)^exp^–MBNL1 complex in this NanoBRET assay in a dose-dependent manner with an IC_50_ of 40 ± 8 µM (**Figure 5C**). Each of the 35 analogs was screened at a single dose of 50 µM in biological triplicate (**Figure S16A**). Of these, sixteen are inactive (reduction of BRET signal by < 20%), thirteen compounds showed reduced activity relative to **2**, three showed similar activity, and three analogs (**31**, **32** and **33**) showed greater disruption of the complex (**Figure S16A** These three compounds exhibited minimal cytotoxicity (<10%) at the screening concentration (**Figure S16B**). Dose–response analysis of the three more potent analogs afforded IC_50_ values of 1 ± 0.3 µM for **31** (40-fold more potent than **2**), 8 ± 3 µM for **32** (5-fold more potent), and 39 ± 5 µM for **33** (equipotent) (**Figure 5C**). Compound **31**, which combined the greatest increase in potency with improved solubility, was selected for further mechanistic studies.

Analysis of the NanoBRET provide preliminary insight into SAR across the PAS-similarity derived series. Although the library is modest in size, the compounds can be binned into three categories based on NanoBRET activity (**Figure S17**). The three most potent small molecules identified in the single-dose screen, **31**, **32**, and **33**, share a conserved 2-thiazole scaffold, in which a sulfur-containing thiazole ring links an aromatic heterocycle to an amide-bearing side chain. This thiazole-centered framework is preserved across the active analogues and likely contributes to RNA recognition through a combination of heteroaromatic stacking and hydrogen-bonding functionality.

### Compound 31 rescues alternative pre-mRNA splicing defects in DM1 patient-derived myoblasts

The activity of **31** in DM1 patient-derived and WT (healthy) myoblasts was evaluated using the same MYOD1-induced myogenic differentiation system described above. Compared to **2**, which showed modest activity at 50 µM (37 ± 2% rescue; p = 0.0013), **31** exhibited improved potency with statistically significant rescue at the 10 µM dose (39% ± 2% rescue; p = 0.001), 20 µM dose (44% ± 2% rescue; p = 0.0007), and 50 µM dose (58% ± 4% rescue; p = 0.0002), affording an IC_₅₀_ of 20 ± 4 µM (**Figure 5D and S7**). As observed for **2** (**Figure S8**), rescue of *MBNL1* exon 5 inclusion rate was not driven by reduction in total MBNL1 transcript levels (**Figure S9**). Notably, an increase in the MBNL1 exon 4–6 isoform was observed following treatment with 50 µM of **31** (1.24 ± 0.07-fold; p = 0.0405), consistent with a modest rescue of this isoform upon compound exposure (**Figure S18)**. RT-qPCR analyses confirmed that treatment with **31** (50 µM) did not alter *DMPK* or *MyoD* transcript levels in DM1 myoblasts, and cell viability remained above 80% at the tested doses, indicating minimal cytotoxicity. (**Figures 5E and S18**). Additional RT-PCR analysis revealed dose dependent, partial restoration of normal splicing patterns of another MBNL1-regulated transcript, *MBNL2* exon 5 alternative splicing^62^ (**Figure S19**). *MBNL1* exon 5 splicing in WT myoblasts was unaffected by **31**-treatment (50 µM) as was *MAP4K4* exon 22a alternative splicing (Nova-regulated) in DM1 myoblasts (**Figures 5E and S8**).

### Compound 31 binds selectively to r(CUG) internal loops

In the NanoBRET assay, signal reduction could in principle arise from binding to either r(CUG)^exp^ or MBNL1 protein. Binding directly to MBNL1 is undesirable, as it may interfere with MBNL1’s native role in alternative pre-mRNA splicing. To confirm RNA engagement, binding of **31** was evaluated by NMR spectrometry using an RNA containing four r(CUG) repeats capped with a GAAA hairpin loop. This RNA forms two adjacent 1×1 U/U internal loops closed by GC base pairs (5’-C<u>U</u>GC<u>U</u>G-3’/3’-G<u>U</u>CG<u>U</u>C-5’ where the underlined nucleotides indicate the internal loop). Binding was assessed by monitoring changes in imino proton resonances, including peak broadening, intensity loss, and chemical shift perturbations, which reflect localized structural or dynamic changes upon ligand interaction.

Compound **31** produced dose-dependent perturbations in imino resonances of loop nucleotides (U5/U19 and U8/U22) and adjacent closing base pairs (G6/G20 and G9/G23) (**Figure S20A**). These effects, observed in a buffer containing 5 mM KH_2_PO_4_/K_2_HPO_4_, pH 6.0, and 50 mM NaCl, demonstrate direct engagement of the U/U internal loops. Minimal perturbations were observed for canonically base-paired regions, indicating selectivity for the pathogenic internal loop motif (**Figure S20B**). Analysis of concentration-dependent imino perturbations yielded a K_Dest_ of 34 ± 6 µM (**Figure S20C**). Consistent with the imino proton titration, monitoring the pyrimidine resonances of r(CUG)_₂_ during ligand titration using a 2D ¹H TOCSY experiment further confirmed binding of **31** to the internal loops. Specifically, significant line broadening and chemical shift perturbations (CSPs) were observed for U5, along with CSPs for U14 and the adjacent nucleotide C15, indicating interaction of the ligand with the U/U internal loop and its neighboring residues (**Figure S21**).

### Compound 31 engages longer r(CUG) repeats

The interaction of **31** with longer repeats, r(CUG)_42_, was evaluated by ligand-observed NMR methods, as described above for **2**, namely chemical shift perturbation^63^ and CPMG^64^ relaxation filtering studies. In 1D spectra, addition of r(CUG)_42_ caused pronounced line broadening and intensity loss of multiple ligand resonances, particularly protons 5 and 7 from the benzimidazolinone (**Figure S22A**), consistent with the attenuation of those same signals in the CPMG experiments (**Figure S22B**). These results are in line with the effects observed on proton 6 of the indane ring from **2** (**Figure S11**). Together, these complementary experiments confirm that **31** directly engages expanded r(CUG) repeats and form a stable complex under the conditions tested. Importantly, no binding of **31** was observed to MBNL1, as assessed by STD NMR spectrometry under the same buffering conditions and concentrations used in CSP and CPMG studies of r(CUG)_42_ (**Figure S23**).

### Molecular dynamics simulations support binding of 2 and 31 to U/U internal loops in r(CUG)^exp^

All-atom, unbiased molecular dynamics (MD) simulations provide a useful framework for characterizing small molecule binding modes without imposing constraints on ligand or target conformations while explicitly accounting for solvent and ion effects.^65^ Thus, to further support target engagement of the U/U internal loops in r(CUG)^exp^ by **2** and **31**, 10 µs unbiased (“brute-force”) MD simulations were performed. Separate simulation systems were constructed containing r(CUG)_12_, water, ions (Na^+^/Cl^-^), and either compound **2** or **31**. Both compounds interacted preferentially and persistently with 1×1 nucleotide U/U internal loops. Compound **2** localized within the widened minor-groove pocket formed by the 1×1 nucleotide U/U internal loops, where the uracil bases adopt conformations partially flipped out of the helical axis (**Figure S24A**). This binding mode is consistent with previously reported solution structures of r(CUG) repeats, which show that the U/U internal loops are conformationally dynamic and can transiently sample partially flipped or open states.^66^ These dynamic structural fluctuations are thought to generate transient binding pockets, providing opportunities for small molecules to recognize and stabilize specific loop conformations. The aromatic core of **2** stacked against the exposed uracil bases, while adjacent heterocyclic moieties formed directional hydrogen bonds with the O2 and N3 atoms of the uracils. The ligand remained associated with the 1×1 nucleotide U/U internal loops throughout the simulation. Clustering analysis revealed two closely related conformational states differing mainly in minor rotations of the aromatic system and substituent orientations. The low RMSD between cluster centroids (<1.5 Å) is consistent with a stable and well-defined binding mode. Within the limitations of the simulation time, three of the four loops were occupied by one ligand with similar orientation of the aromatic section (Figure S15A). Although stacking interactions are not directly observable in NMR studies, the significant changes in proton 6 of **2** in both CSP and CPMG confirm the engagement of the aromatic core in line with MD observation.

Compound **31**, the lead-optimized analog, formed a more extensive interaction network with the U/U loop than **2** did (**Figure S24B**). In addition to π-stacking with the uracil bases, **31** established multiple hydrogen bonds involving O2, O2′ and N3 atoms of the mismatched UU nucleotides, resulting in a persistent hydrogen-bonding network over the course of the trajectory. Additional stacking interactions with neighboring nucleotides increased burial of the ligand within the binding pocket. Clustering analysis again revealed two dominant, long-lived conformational states, consistent with a stable RNA–ligand complex. Notably, the benzimidazolinone protons that broaden most strongly in ligand-observed NMR are positioned at the stacked interface in the MD cluster centroids, consistent with direct participation of this ring in the bound complex. The MD simulations reveal partial flipping of the U/U loop nucleotides upon ligand binding consistent with the 2D ^1^H TOCSY.

Molecular mechanics generalized Born surface area^67^ (MM/GBSA) free energy analysis predicts more favorable binding of **31** to the U/U internal loop as compared to **2**, with estimated binding energies of –38.3±4.1 kcal/mol and –30.2±2.88 kcal/mol, respectively. Although absolute MM/GBSA values should be interpreted cautiously, the relative stabilization is consistent with the more extensive interaction network observed for **31** in the MD simulations and with its stronger experimental engagement. Overall, the unbiased MD simulations support selective binding of **2** and **31** to the U/U internal loops in r(CUG)^exp^ and suggest that the expanded interaction network observed for **31** contributes to its enhanced cellular activity.

## DISCUSSION

A major challenge in RNA-targeted small-molecule discovery is the low hit rate that typically emerges when screening general purpose compound libraries against structured RNA targets. This reflects both the inherent conformational complexity of RNA and the fact that most screening collections were not designed to recognize RNA structural motifs such as internal loops, bulges, and other noncanonical pairs. In this work, we sought to address this challenge by combining an augmented RNA-focused compound library with biological triaging of a small, prioritized subset of compounds, enabling efficient identification of molecules that not only bind RNA but also modulate disease-relevant biology.

Using this strategy, we identified initial lead compounds that selectively engage the U/U internal loops formed when r(CUG)^exp^ folds. While these early hits demonstrated target engagement and biological activity, they also exhibited limited aqueous solubility, a recurring issue in RNA-targeted chemical matter.^17^ Rather than abandoning the chemotype, a systematic framework combining positional analogue scanning^16^ with similarity-based expansion was implemented to improve solubility while retaining RNA-binding properties. This approach yielded **31**, which showed enhanced solubility (600-fold) and low– to mid-micromolar activity in DM1 patient-derived myoblasts, representing a clear improvement over the initial hit and demonstrating that solubility liabilities can be addressed without loss of target engagement.

There is increasing interest in using computational approaches, including machine learning– and artificial intelligence–augmented methods, to identify RNA-binding small molecules. However, such approaches require rigorous experimental validation, as computational predictions alone cannot establish binding mode, selectivity, or functional impact on disease-relevant pathways. In this study, we employed a tiered and orthogonal validation strategy spanning in vitro RNA binding and RNA–protein complex disruption, live-cell target engagement assays, and functional testing in patient-derived myoblasts. Single-molecule magnetic force spectroscopy was also incorporated, which provided unique mechanistic insight into RNA folding dynamics and ligand-dependent modulation of RNA–protein interactions. This single-molecule assay revealed features of compound behavior—such as partial inhibition of MBNL1 binding and altered folding equilibria—that are not accessible by ensemble biochemical or cellular assays alone, underscoring its value as a complementary and orthogonal method in RNA-targeted discovery. Molecular dynamics simulations further supported the experimental findings by providing atomistic insight into how **2** and **31** engage U/U internal loops, consistent with experimentally observed selectivity.^65^ These simulations were used as supportive evidence, rather than as a standalone validation, reinforcing the importance of integrating computational modeling with biochemical, biophysical, and cellular data.

Despite these advances, several limitations should be acknowledged. Because this study relied primarily on commercially available compounds, the chemical space explored was constrained, and the most potent molecules identified remain in the low micromolar range both for affinity and cellular activity. While sufficient to establish proof of concept and demonstrate activity in disease-relevant cells, additional medicinal chemistry optimization will be required to achieve nanomolar potency and favorable in vivo properties. Such efforts will be essential to translate RNA-targeted discovery strategies into therapeutically investable molecules.

In summary, this work demonstrates that combining RNA-focused library design, solubility-driven hit expansion, and rigorous multi-level biological validation—including single-molecule measurements—can help overcome key barriers in RNA-targeted small-molecule discovery. Continued advances in RNA target definition, compound optimization, and medicinal chemistry will be critical for fully realizing the therapeutic potential of RNA-targeted small molecules, and these studies provide a framework for those future efforts.

## METHODS

### General

RNAs were purchased from Dharmacon (GE Healthcare). Deprotection of 2’-ACE protected RNAs and desalting using PD-10 columns (Cytiva, 17085101) were performed according to each manufacturer’s recommended procedure. Concentrations of RNA stock solutions were quantified by UV/Vis spectrometry with a Beckman Coulter DU 800 spectrophotometer by measuring absorbance at 260 nm at 80 °C and using the extinction coefficient provided by the manufacturer. DNA primers were purchased from Integrated DNA Technologies (IDT, Inc.) (**Table S3**).

### Molecular similarity search

RDKit^68^ (Version 2.3) was used for the implementation of the atom-pair fingerprint to compute the fingerprint vectors. Tversky coefficients^29^ were used to measure the similarity between each pair of compounds, *a* and *b*, using their corresponding *l* –bit fingerprint vectors, *v*_*a*_and *v*_*b*_ as following (eq 2):

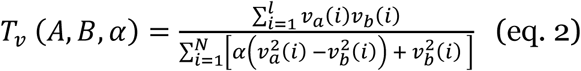

where *A* is the reference molecule, *B* is the database molecule, *l* is the length of the molecular bit vector and *v*(*i*) is the *i* –th element of vector *v*. The factor α weights the contribution of reference molecule *A*. As the α becomes larger the more weight is assigned on the bit settings of *A* and consequently, less on the database molecule *B*. Assuming the Tanimoto coefficient for binary vectors is identified as following:

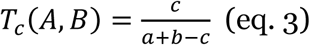

where *a* and *b* are the numbers of bits in molecular fingerprints *A* and *B* and *c* is the number of bits shared between *A* and *B*, Tversky coefficient can be simplified as following:

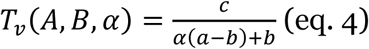

where *a*, *b*, and *c* correspond to the *T*_*c*_. These quantities are identical to those used to compute the Tanimoto coefficient (defined above). The Tversky coefficient generalizes Tanimoto by introducing a weighting parameter to bias similarity toward features of molecule A or B.

### K-means clustering

A python script (https://github.com/PatWalters/kmeans) that provides an implementation of K-means clustering using mini batch K-means from SciKit^69^ (version 1.2.2) combined with fingerprints calculation from RDKit was used for cluster analysis. K-means^30^ algorithm is a member of prototype-based clustering methods in which each cluster is represented by either a centroid (cluster center) or a medoid (same as centroid but restricted to be member of the data set). Regarding the large number of molecules, a two-step clustering approach was adopted. First, a subset (n = 20,000) of the whole library (n = 8,000,000) was selected randomly for clustering creating the initial cluster centers. Then, the Euclidean distance between the fingerprint of each molecule in the library and the fingerprint of the cluster centers is calculated and each molecule is assigned to the closest cluster center.

### Chemical space mapping and calculation of drug-likeness

Uniform Manifold Approximation and Projection (UMAP)^32^ was used for visualization of the chemical space and dimensionality reduction using the atom-pair molecular fingerprint. Quantitative estimation of drug-likeness (QED)^36^ which reflects the distribution of molecular properties like molecular weight, logP, topological polar surface area, number of hydrogen bond donors and acceptors, the number of aromatic rings and rotatable bonds, and the presence of unwanted chemical functionalities was measured with RDKit implementation (https://www.rdkit.org/docs/source/rdkit.Chem.QED.html).

### Scaffold analysis

The Bemis-Murcko framework^7^ was used for scaffold analysis. In this method, monovalent atoms are successively removed from a compound until only ring atoms and linker atoms remain. RDKit implementation of the Murcko scaffolds was used to extract and count the scaffolds present in the parent library.

### 5’-32P labeling of r(CUG)_10_

Approximately 1 nmole of RNA was radiolabeled with [γ-^32^P] ATP (PerkinElmer) using T4 polynucleotide kinase (New England Biolabs) at 37°C for 45 min and purified by using a denaturing 15 % (v/v) polyacrylamide gel. The RNA was visualized by UV shadowing wherein it was excised from the gel, eluted via tumbling in 300 mM NaCl at 4°C overnight. The resulting RNA was then purified by ethanol precipitation (20 μg of glycogen (RNA grade; Invitrogen), and the sample was incubated for 15 min at –80°C. The sample was centrifuged at max speed for 10 min at 4°C to pellet the RNA, which was dissolved in 40 μL of Nanopure water.

### Screening of the RNA-focused library by *in vitro* competitive Chem-CLIP

*In vitro* Chem-CLIP experiments were carried out following established procedures.^45^ In brief, 5’ end ^32^P labeled RNA (600K CPM) was folded in 20 mM HEPES, pH 7.5, by heating at 95°C for 1 min followed by snap-cooling on ice. A 65 µL aliquot of the folded RNA was placed into 0.6 mL microcentrifuge tubes and supplemented with competitor compound at the indicated concentrations. Following incubation at room temperature for 15 min, 50 µM of **1** was added to each sample which was incubated for an additional of 15 min at room temperature. Subsequently, each reaction tube was irradiated with UV light (365 nm) for 15 min by UVP crosslinker (UV Stratalinker 2400).

Following irradiation, a freshly prepared click-reaction mixture—consisting of CuSO₄ (1 µL, 10 mM), THPTA ligand (0.6 µL, 50 mM), PEG₃-biotin azide (1.0 µL, 10 mM), and sodium ascorbate (0.6 µL, 250 mM, pH 7.0)—was added to each sample, and the reaction proceeded for 3 h at 37 °C. Streptavidin magnetic beads (15 µL slurry; Dynabeads MyOne Streptavidin C1) were then introduced, and samples were incubated for an additional 15 min at room temperature to capture biotinylated, cross-linked RNA–ligand adducts. Unbound RNA was removed using magnetic separation, and the beads were washed three times with Wash Buffer (10 mM Tris-HCl pH 7.0, 1 mM EDTA, 4 M NaCl, 0.2% Tween-20). Radioactivity associated with both the bead-bound and supernatant fractions was quantified by liquid scintillation counting.

### Cell culture & compound treatment

C2C12 mouse myoblast cells stably expressing either r(CUG)_0_ or r(CUG)_800_ in the 3’ UTR of luciferase^70^ were cultured in Dulbecco’s Modified Eagle Medium (DMEM; Corning, #15-017-CV) supplemented with 10% (v/v) fetal bovine serum (FBS; Gibco, #10437-028), 1% (v/v) Antibiotic-Antimycotic Solution (Corning; #30-004-CI), and 1% (v/v) Glutagro (Corning, #25-015-CI) in 100 mm diameter dishes up to passage 30. For compound treatment, cells were seeded into 96-well plates and cultured in growth medium until ∼80% confluent. Cells were treated with compound supplemented medium (0.1% (v/v) DMSO) for 24 h at 37°C.

DM1 conditional MyoD-fibroblast cells with 1300 r(CUG) repeats and wild-type conditional MyoD-fibroblast cells^47^ (gifts from D. Furling; Centre de Recherche en Myologie (UPMC/Inserm/CNRS), Institut de Myologie, Paris France) were maintained in Dulbecco’s Modified Eagle Medium (DMEM; Corning, #15-017-CV) supplemented with 15% (v/v) FBS (Gibco, #10437-028), 1% (v/v) Antibiotic-Antimycotic Solution (Corning; #30-004-CI), and 1% (v/v) Glutagro (Corning, #25-015-CI) in 100 mm diameter dishes up to passage 28. For compound treatment, cells were seeded into 6-well plates and cultured in growth medium until ∼90% confluent. Conditional MyoD-fibroblast cells were then differentiated in the presence of compound into myotubes for 48 h using a differentiation medium composed of 1× DMEM, 1% (v/v) Antibiotic-Antimycotic Solution, 0.1 mg/mL transferrin human (Sigma, #T8158), and 0.01 mg/mL insulin (Sigma, I0516).

HeLa480 cells^71^ were maintained in Dulbecco’s Modified Eagle Medium (DMEM; Corning, #15-017-CV) supplemented with 10% (v/v) FBS (Gibco, #12676029), 1% (v/v) Antibiotic-Antimycotic Solution (Corning, #30-004-CI), and 1% (v/v) Glutagro (Corning, #25-015-CI).^71^ Cells were cultured at 37 °C in a humidified atmosphere containing 5% CO_₂_ up to passage 25. These cells were used for NanoBRET assays for which more details can be found below.

### Rescue of the *DMPK* nucleocytoplasmic transport defect in the C2C12 luciferase assay.^70^

Mouse C2C12 cells stably expressing r(CUG)_0_ or r(CUG)_800_ in the 3’ UTR of luciferase were plated in 96-well plates (4,000 cells/well) and treated as described above. Following compound treatment, cell viability was measured using 5 µL/well of CellTiter-Fluor (Promega) following the manufacturer’s protocol. After a 30 min incubation at 37°C, fluorescence was measured using a SpectraMax M5 plate reader (λ_ex_ = 380 nm and λ_em_ = 505 nm). To measure luciferase activity, 25 µL of One-Glo (Promega) was added to each well following the manufacturer’s protocol. Cells were incubated for 5 min at room temperature in the dark and luminescence was measured using a SpectraMax M5 plate reader with an integration time of 500 ms. Luminescence signal was normalized to cell viability, and the ratio of luminescence/cell viability for vehicle-treated cells was mathematically normalized to 1 to assess relative changes in luminescence upon compound treatment.

### RT-qPCR analysis of splicing defects and *DMPK* and *MyoD* abundances in DM1 and WT myotubes

DM1 patient-derived fibroblasts were seeded in 12-well plates and maintained in growth medium until reaching 100% confluency. The growth medium was replaced with differentiation medium with or without compound (final concentration of 0.2% (v/v) DMSO), and the cells were incubated for 48 h. Following compound treatment, total RNA was harvested using a Zymo Research Quick-RNA Mini Prep Kit following the manufacturer’s protocol with DNase I treatment. The concentration of RNA was determined by a Nanodrop UV/vis spetrophotometer, and 500 ng of total RNA was reverse transcribed with a qScript cDNA synthesis kit (20 µL total reaction volume, Quanta BioSciences) following the manufacturer’s protocol. Subsequently, 2 µL of the reverse transcription reaction was amplified using GoTaq DNA polymerase (Promega) per manufacturer’s protocol in a total volume of 25 µL. RT-PCR was carried out under the following conditions for 30 cycles: 95°C for 30 s, 58°C for 30 s, 72°C for 1 min, with final extension at 72°C for 5 min using primers listed in **Table S3** for *MBNL1* exon 5 or *MAP4K4* exon 22a. The amplified PCR products were analyzed and quantified using a 5300 Fragment Analyzer System (Agilent Technologies) with the dsDNA 910 Reagent Kit (35–1500 bp, #DNF910K0500).

Next, 3.5 µL of the same cDNA samples were subjected to a 1:10 dilution in a SYBR Green (Thermo Fisher Scientific) qPCR reaction using primers for *DMPK* and *MyoD* primers (570 nM each; **Table S3**). The qPCR reaction was then aliquoted into three technical replicates (10 µL) and analyzed by a QuantStudio 5, 384-well Block Real-Time PCR System (Applied Biosystems). Relative abundance of each transcript was determined by normalizing to the housekeeping gene (*GAPDH*) using the 2^-ΔΔCt^ method.^72^

### NanoBRET: live-cell r(CUG) repeat-MBNL1 displacement assays

Complex formation between r(CUG) repeats and MBNL1 *in live cells* was measured using a previously reported NanoBRET assay.^73^ Briefly, HeLa480 cells were co-transfected with 2500 ng MBNL1-HaloTag and 50ng MBNL1-NanoLuc plasmids in a 6-well plate using 8 µL of FuGENE® HD Transfection Reagent (Promega, #E2311) in 110 μL of Opti-MEM (Giboco, #31985070), following the manufacturer’s protocol. After incubating the cells overnight, they were trypsinized and resuspended in an appropriate volume of assay medium (Opti-MEM + 4% (v/v) FBS). The cell suspension was then re-seeded into white 96-well tissue culture plates (Corning, #3903) where 100 μL of growth medium containing 200 nM HaloTag NanoBRET 618 Ligand (Promega, #N1662) was added to each well, delivering 1.6 × 10⁴ transfected cells (affording ∼70% confluency). Additionally, for each experiment, cells were dispensed into three wells containing 100 µL of growth medium with no HaloTag ligand. Approximately 4 h after reseeding, the compound of interest was added to each well at the indicated concentrations by delivering 10 µL of compound solution prepared in assay medium (final concentration of 0.1% (v/v) DMSO). The following day, 1× NanoLuc substrate (Promega, #N1662) and 20 μM extracellular NanoLuc inhibitor (Promega, #N2160) in the 25 μL assay medium were added to each well. The interaction between MBNL1 and r(CUG)_480_ was measured by detecting the BRET signal using the GloMax Discover System (Promega, #GM3000). Dual-filtered luminescence was collected with a 460/80 nm bandpass filter for the donor (NanoLuc protein) and a 610 nm long-pass filter for the acceptor (HaloTag ligand) using an integration time of 500 ms. The corrected NanoBRET ratio, expressed in milliBRET units (mBU), was calculated using Equation 1:

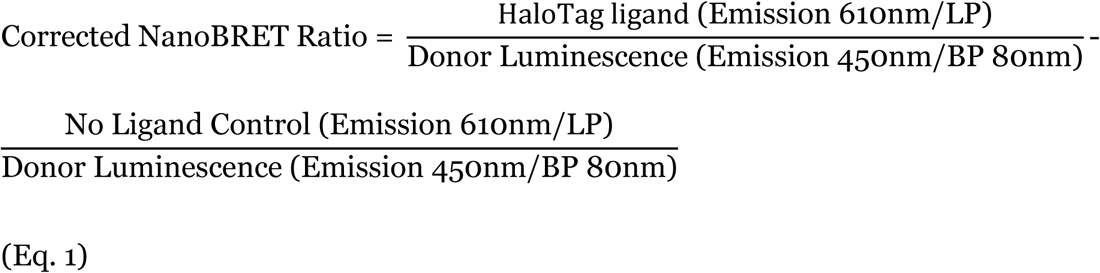

### NMR spectroscopy

Ligand-based NMR experiments were acquired on a Bruker Avance III 600 MHz spectrometer equipped with a cryoprobe. RNA-based experiments were acquired on a Bruker Advance III 700 MHz equipped with a cryoprobe. Duplex RNA, r(GACAGC<u>U</u>GCUGUC)_2_ where the U nucleotide that forms the loop is underlined) was purchased from Dharmacon (GE Healthcare) as an HPLC-purified oligonucleotide. The r(CUG)_42_ (5’-GGGCGGG(CUG)_42_GGGGAUC-3’; 140 nt; forms 42 1×1 nucleotide U/U internal loops) and r(CUG)_2_ (5’-GCGC<u>U</u>GCU<u>GAAAA</u>GC<u>U</u>GCGC-3’, where internal and hairpin loop nucleotides are underlined; 20 nt) hairpin construct were synthesized by in vitro transcription. Fully base paired control RNA where UU loop was replaced with AU base pair (5’-GCGCUGCU<u>GAAAA</u>GCAGCGC-3’) was purchased from Dharmacon (GE Healthcare) as an HPLC-purified oligonucleotide.

The DNA template for r(CUG)_42_ was obtained from kanamycin-resistant pCTG109 vector previously cloned from a pCNA plasmid containing the repeat expansion sequence.^74^ The plasmid was linearized using BamHI restriction enzyme (New England Biolabs, Ipswich, MA, USA). The linearized plasmid, at a concentration of 25 mg/mL, was in vitro transcribed using a 32 mM NTP mix, 48 mM MgCl_₂_, 0.07 mg/mL T7 RNA polymerase and 8% (v/v) DMSO in Transcription Buffer containing 40 mM Tris, pH 8, 1 mM spermidine, 0.01% (v/v) Triton X-100 and 5 mM DTT. The transcription reaction was incubated for 4 h at 37 °C. The RNA was then purified using a 12% (29:1) acrylamide:bisacrylamide denaturing gel containing 7 M urea. The gel was visualized by UV shadowing at 254 nm, the product band was excised, and the RNA was extracted using passive elution with 3 M sodium acetate, pH 5.3, overnight at 4 °C. The following day, the supernatant was ethanol precipitated twice and desalted using Sephadex G-25 cartridges (Cytiva, Marlborough, MA, USA). The DNA template for r(CUG)_2_ was obtained from IDT.

RNA stocks were diluted with NMR Buffer 5 mM KH_2_PO_4_/K_2_HPO_4_, pH 6.0, and 50 mM NaCl. The duplex RNA was folded by heating at 95°C for 3 min followed by slow cooling to room temperature to favor duplex formation. The r(CUG)_42_, and r(CUG)_2_ oligonucleotides were folded by heating at 95°C for 3 min followed by snap cooling on ice to favor hairpin formation. For all NMR experiments, the compound of interest was dissolved in 99.8% DMSO-d_6_ to minimize the signal from the DMSO.

All ligand-based NMR samples were acquired in 10 mM KH_2_PO_4_/K_2_HPO_4_, 50 mM NaCl, pH 6.0, 0.05 mM EDTA, 5% (v/v) DMSO-d_6_ (Cambridge Isotope Labs) and 10% (v/v) D_2_O (Cambridge Isotope Labs) at 298 K. In brief, two samples were prepared per compound: one sample containing the compound alone at concentration of 100 μM and a second sample containing the compound at the same concentration and r(CUG)42 at an internal-loop:compound ratio of 1:2 (**2**) or 1:50 (**3**). 1D ^1^H and CPMG ^1^H ligand-based experiments^56^ were used to evaluate the interaction of **2** and **31** with r(CUG)_42_. CPMG experiments were collected with relaxation times of 6, 30, 300 and 600 ms. The water signal in both experiments was suppressed by excitation sculping with gradients.^75^

For RNA-based experiments, the interaction of **31** with r(CUG)_2_ was monitored with 1D and 2D (TOCSY, 60 ms mixing time) experiments at increasing RNA:ligand molar ratios of 1:0, 1:1, and 1:2 at 298 K. The NMR sample, prepared in 10 mM KH_2_PO_4_/K_2_HPO_4_, 50mM NaCl, pH 6.0, 0.05 mM EDTA in 100% D_2_O contained 200 μM r(CUG)_2_. The assignment of the imino protons was based on the analysis of the exchangeable/non-exchangeable ^1^H resonances from NOESY experiments (100 and 400 ms).

1D ^1^H NMR spectra of exchangeable (imino) protons (10-15 ppm) for both constructs (r(CUG)_2_ and r(CUG-CAG)_2_) were acquired in 5% D_2_O and 95% H_2_O at 5 °C (278K) using 50 μM of RNA in the absence of compound. Compounds were then dissolved in D6-DMSO and added to the RNA sample to achieve a final concentration of 100 μM (1:2 RNA:compound). The chemical shifts were referenced to the residual solvent peaks of D6-DMSO, which served as the internal calibration standard. NMR spectra were processed in Topspin 4.0.6.To quantify the binding affinity of **31**, chemical shift perturbations (CSP) of protons in **31** was measured as a function of r(CUG)_2_ concentration. A 50 μM solution of **31** was prepared in a final volume of 400 µL (Shigemi NMR tube). RNA (0.5 μL aliquots) was then added to afford concentrations ranging from 10-250 µM. The resulting curves plotting changes in CSPs as a function of RNA concentration were fitted using the one site specific binding model within GraphPad Prism. Chemical shift perturbations (CSPs) were quantified by measuring changes in the peak positions, and line broadening (LB) was monitored by observing reductions in peak intensity and increases in linewidth. Chemical shifts were internally referenced using residual DMSO present in the sample.

Saturation Transfer Difference (STD) NMR experiments were performed to evaluate potential binding of **2** and **31** to the MBNL1 protein. Samples were prepared in 5 mM KH_2_PO_4_/K_2_HPO_4_, 50 mM NaCl, pH 6.0, 0.05 mM EDTA containing 10% (v/v) D₂O for field locking. Protein samples contained MBNL1 2.5 µM, while ligands were added at a 20-fold molar excess (50 µM) to ensure efficient saturation transfer. Ligands were dissolved in DMSO-d_6_. STD NMR spectra were acquired at 298 K on a 600 MHz NMR spectrometer using a standard STD pulse sequence. Selective protein saturation was applied at 0.7 ppm (on-resonance), while an off-resonance frequency at 40 ppm served as the reference. A train of Gaussian-shaped pulses was used to achieve a total saturation time of 2 s. Difference spectra were obtained by subtracting on-resonance and off-resonance spectra.

Binding constant of compound **31** to r(CUG)_2_ repeats was evaluated by NMR titration experiments. r(CUG)₂ RNA was titrated into a solution containing a constant concentration of **31** (50 μM). Upon addition of RNA, chemical shift perturbations were monitored in the ¹H NMR spectrum of the ligand. In particular, the downfield shift of the O-methyl proton resonance at ∼3.6 ppm was used as a reporter for ligand–RNA interaction. The magnitude of the chemical shift change was plotted as a function of RNA concentration to generate a binding isotherm. Dissociation constants (K_Dist_) were determined by fitting the titration data using a one-site binding model implemented in GraphPad Prism.

### Single molecule studies (magnetic tweezers)

The RNA template used for the single molecule analysis consists of 21 r(CUG) repeats (63 nt, **Table S4**) flanked by 6 nucleotides of random single-stranded RNA bases as well as two constant sequences allowing the annealing of DNA handles for surface (3′ DNA splint in **Table S4**) and bead (5′ DNA splint in **Table S4**) attachment. The RNA was chemically synthesized by Dharmacon and tagged with a 5′-biotin. The DNA handles were synthesized by Eurofins.

The RNA was annealed to two DNA splints in equimolar ratio in the annealing buffer (10 mM Tris-HCl, pH7.4, 50 mM NaCl, 1 mM EDTA). Once annealed, the structures were purified through MicroSpin S-200 HR spin columns (Cytiva) and were mixed with an equal volume of GenTegraRNA (GenTegra) before storing at –20°C until further use.

The annealed RNA structure was mixed at a concentration of 4 fmol with 3 µL of MyOne T1 streptavidin beads (Invitrogen) in Hybridization Buffer (10% (w/v) PEG-8000, 5× SSC Buffer) for 10 min at room temperature. The beads were washed twice in buffer (50 mM HEPES, pH 7.4; 100 mM KCl; 1 mM MgCl_2_; 1% DMSO) to remove unbound RNA structures and the resulting beads containing the RNA were resuspended in the buffer before loading in the instrument.

*Preparation of muscleblind-like 1 (MBNL1) protein.* Human recombinant MBNL1 protein was obtained from Abnova (H00004154-P02, Lot MC091). To perform the tests, both **2** and MBNL1 protein were diluted at the indicated concentrations in Test Buffer (20 mM HEPES, pH 7.4, 100 mM KCl, 2 mM MgCl_2_; 5 mM DTT, and 1% (v/v) DMSO), maintaining 1% (v/v) DMSO as a final concentration.

*Instrument (MAGNA^TM^ technology) and experimental measurements.* All experiments presented in this paper were acquired on a Stereo Darkfield Interferometry system (SDI) MAGNA™ instrument.^76^ To attach the RNA molecules at the surface of the flow cell, a functionalized azide-coated coverslip (Susos) was grafted with a surface oligo (100 nM) harboring a 3′-DBCO group through click chemistry in the Click Buffer (500 mM NaCl, 1 µM PEG-DBCO) for 2 h. The flow cell was then passivated with OB after assembly. The RNA bound to the MyOne beads was injected into the flow cell and left to hybridize for 30 min in OB. After the molecules were captured at the surface of the flow cell, the buffer was exchanged to Test Buffer. The unbound structures were washed away, and the non-specifically bound beads were removed by increasing the magnetic force above 25 picoNewton (pN). The experiments were performed at 22 °C, and the data were recorded at a frequency of 30 Hz.

A control experiment was always recorded first in Test Buffer, which was used to normalize as well as to define the number of analyzable molecules. Then, experiments were performed with increasing concentration of either compound **2** or MBNL1 protein in the Test Buffer, maintaining the concentration DMSO at 1%. In experiments where both compound **2** and MBNL1 protein were tested together, the compound and the protein were mixed in Test Buffer (with DMSO at a final concentration of 1%) just prior to injecting into the flow cell.

Two types of experiment were performed: 1) Ramp experiments, consisted of moving the magnet position slowly (0.125 mm/s) to increase the force applied to the beads from ∼0.2 pN to ∼28 pN, then maintaining the highest force for 1s before moving back down to low force, with repetition of these steps for 100 cycles; 2) For constant-force experiments, the magnet was maintained at a constant position corresponding to a constant force for 30 s, starting at ∼6 pN, then moved closer to the flow cell surface in a stepwise manner until the force reached ∼19 pN.

*Analysis of the ramp experiments.* The 30 Hz recorded bead positions from every cycle of each molecule were first cleaned by removing signals with high noise (>1000 nm). Then, in order to be considered as an analyzable RNA molecule to be included in downstream analyses, the molecules had to satisfy the following parameters: 1) unfolding force between 5-15 pN; 2) unfolding size between 5-30 nm in DMSO conditions; 3) the total number of clean cycles must be above 20 cycles; 4) the control condition has more than 80% of clean cycles; 5) data have been successfully recorded on the molecule in more than half the conditions tested over the course of a single experiment; and 6) the distribution of 25%-75% of the unfolding force in control conditions is less than 2 pN. Following this selection, each cycle of each molecule was analyzed to determine the force required to unfold and fold the RNA structures as well as the size of the detected jump using the Python module HDBScan.^77^ The RNA structure was considered closed if all cycles did not contain any unfolding or refolding event. For each analyzable structure, we plotted each cycle if we detected an unfolding (+1) or refolding event (−1) whereas if no event was detected, it was plotted as 0. If the cycle was too noisy, it was annotated as ‘NA’. The unfolding and refolding probabilities were calculated by dividing the number of cycles with a jump event by the total number of cycles for a single molecule. The median unfolding and refolding probability of all structures were plotted against ligand or protein concentrations.

The median unfolding force of each analyzable molecule in the DMSO condition (over all cycles) was used to normalize all cycles of all conditions for that molecule. The normalized unfolding and refolding forces were plotted to study the impact of ligand or protein binding. The average (±SEM) of the median normalized force of each bead analyzed for a given condition was calculated and plotted against the ligand or protein concentration. The data were fitted with a sigmoid curve.

*Analysis of constant-force experiments*. The 30 Hz data tracking the Z-position of each bead were first cleaned by removing any consecutive data points greater than 40 nm within a rolling window of 20-data points) and discarding all data from any force-step with less than 250 data points. To quantify the amplitude of the folding and unfolding of the RNA structure, the bead displacement variance (squared distance to mean value) per force step was measured using the following method: 1) the data were partitioned into sections of 40 data points; 2) for each of these sections, the average variances over rolling windows of 20 data points were computed; 3) the variance for the force step was determined as the median of the values computed in step 2. This methodology allows compensation for any short-term signal fluctuations and longer-term drift. RNA molecules showing a low signal-to-noise ratio were excluded from further analysis. Finally, to allow for force differences caused by bead size variation, the force applied to each molecule was normalized to the force with maximum variance in the control condition. For plots, the median variances of all the analyzed RNA molecules within intervals of 0.03 along the normalized forces were used and curve smoothing was performed using third order splines.

### Molecular dynamics simulations

AMBER 24^78^ was used for MD simulations using the ff99OL3 (parmbsc0 α/γ^79^ + χOL3 to ff99^80^). The previously elucidated NMR structure of the r(CUG) repeat model with one U/U internal loop (r(GACAGC<u>U</u>UGC<u>U</u>GUC))^66^ was used for MD simulations. The system was first neutralized with Na^+^ ions^81^ and then solvated with OPC^82^ water molecules in a truncated octahedral box with periodic boundary conditions extended to 10 Å using the LEAP^83^ module of AMBER 24. The structures were minimized with the Sander module in AMBER in two steps. Positional restraints on RNA heavy atoms with restraint weights of 10 kcal mol^-1^ Å^-2^ were applied in the first step of minimization with 5000 steps of steepest-descent algorithm followed by 5000 steps of conjugate-gradient algorithm using the CPU implementation of Sander force field to avoid the truncation of forces or overflow of the fixed precision representation.

A second round of minimization was performed without restraints with 10,000 steps of steepest-descent. This minimization was followed by an equilibration protocol first in constant volume dynamics (NVT), where positional restraints were imposed on the RNA heavy atoms with restraint weights of 10 kcal mol^-1^ Å^-2^ while temperature was gradually increased from 0 K to 300 K within several nanoseconds using the Langevin^84^ thermostat. A second round of equilibration was performed at constant pressure (NPT), where temperature and pressure coupling^85^ were set to 300 K and 1.0 ps^-1^, respectively, while constraints were gradually removed.

After minimization and equilibration, MD simulation with a 2 ps time step was performed using NPT dynamics with isotropic positional scaling. The reference pressure was set to 1 atm with a pressure relaxation time of 2 ps. SHAKE^86^ was turned on for constraining bonds involving hydrogen atoms. An atom-based long-range cutoff of 10.0 Å was used in the production runs. The reference temperature was set to 300 K. The Particle Mesh Ewald^87^ (PME) was used to handle the electrostatics and the Langevin^88^ thermostat was applied with a coupling constant *γ* = 1.0 ps^-1^. Simulations were performed using the pmemd.cuda implementation (GPU accelerated) of AMBER24.

*Parametrization of **2** and **31**.* GAFF was used to assign the bonds, angles, torsions (proper (involves four consecutively bonded atoms i-j-k-l) and improper (any other torsion calculated based on four atoms not successively bonded)), improper torsions, and Lennard-Jones parameters using the Antechamber and Parmchk programs.^83^ In order to extract the charges, the compounds were geometry optimized at the quantum-mechanical (QM) HF/6-31G* level using Gaussian 09^89^ consistent with the AMBER force fields. Then atomic charges were determined by restrained electrostatic potential (RESP) charge fitting^90^. RED (RESP ESP charge Derive) program was used to generate the final charges.^91^

### Calculation of interactions

Interactions between the r(CUG) RNA and small molecules were extracted with fingeRNAt.^92^

### Free energy calculations

Binding free energies of ligand–RNA complexes were estimated using the Molecular Mechanics Generalized Born Surface Area (MM/GBSA) method as implemented in the MMPBSA.py module of the AMBER software suite. Snapshots from the equilibrated MD trajectories were extracted at regular intervals (every 10^th^ frame) and used for MM/GBSA analysis.

### Statistical analysis

All statistical analyses were performed using GraphPad Prism 9 (GraphPad Software, San Diego, CA, USA). For experiments involving comparisons among three or more experimental groups, such as compound screening assays, competition experiments, and cellular reporter assays, statistical significance was determined using one-way analysis of variance (ANOVA) followed by Bonferroni post-hoc multiple comparison tests. Statistical significance is indicated as *p* < 0.05*, p* < 0.01, *p* < 0.001, and p < 0.0001.

## Funding

This work was supported by the National Institutes of Health (R35 NS116846 to M.D.D.), Muscular Dystrophy Association (MDA) Development Grant 963835 (to A.T.), and the German Research Foundation (DFG) through a Walter Benjamin fellowship (#515396515 to P.R.A.Z). Purchase of the 600 MHz NMR spectrometer was supported in part by the National Institutes of Health grant S10 OD021550.

## Notes

The authors declare the following competing financial interest(s): M.D.D. is a founder of Ribonaut Therapeutics.

## ACKNOWLEDGMENT

The authors acknowledge University of Florida Research Computing for providing computational resources and support that have contributed to the research results reported in this publication. URL: http://www.rc.ufl.edu. We acknowledge Quentin MR Gibaut and Jessica A Bush for preliminary data.

