## supplemental data for "A chemoinformatics-guided platform for efficient discovery of RNA-binding small molecules: Proof-of-concept for myotonic dystrophy type 1"

[3]Depixus SAS, 39-41 rue Louis Blanc, 92400 Courbevoie, France

### Supplemental Figures

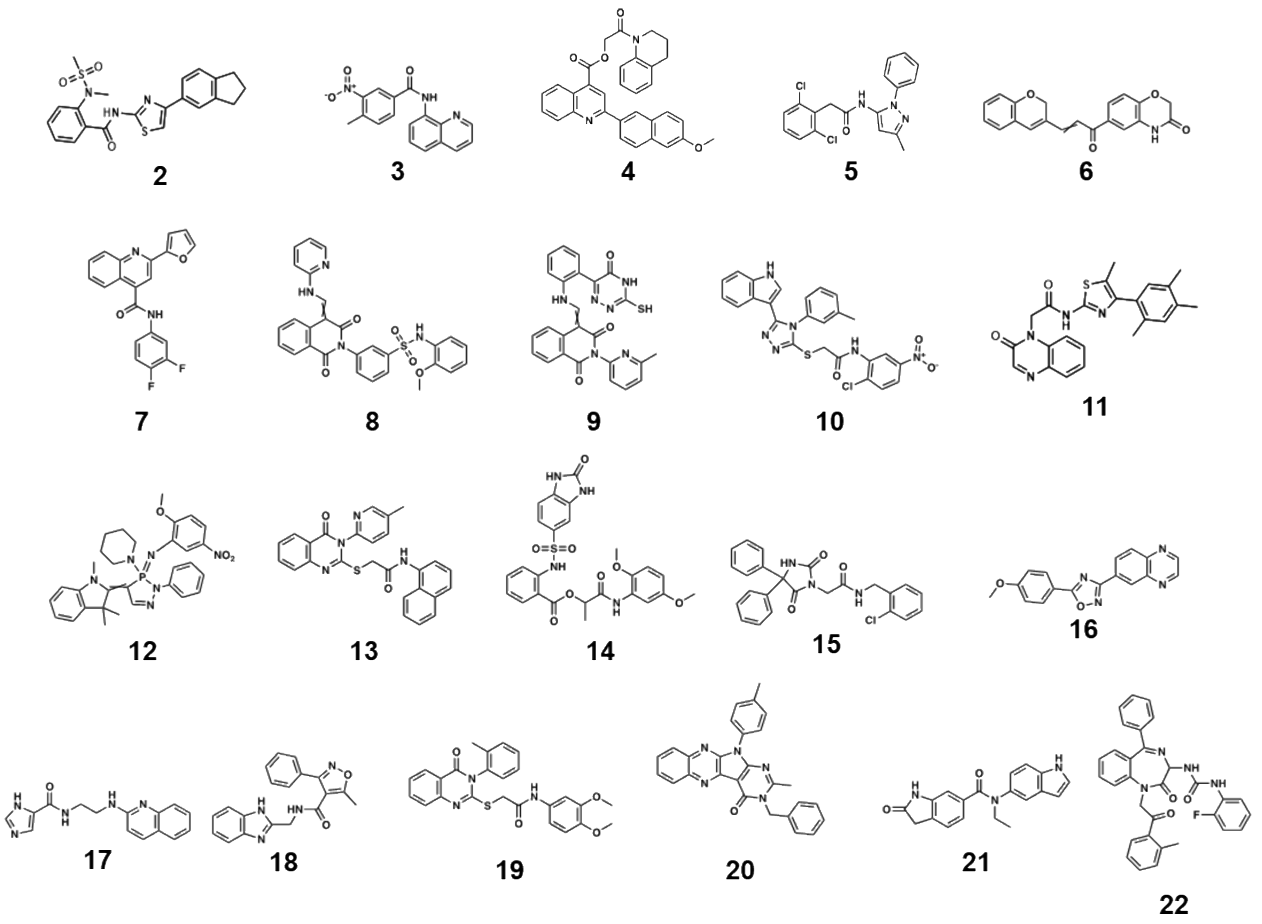


**Figure S1.** Structure of 22 hit compounds identified by an in vitro competitive Chem-CLIP assay to identify small molecules that bind to r(CUG) repeats.

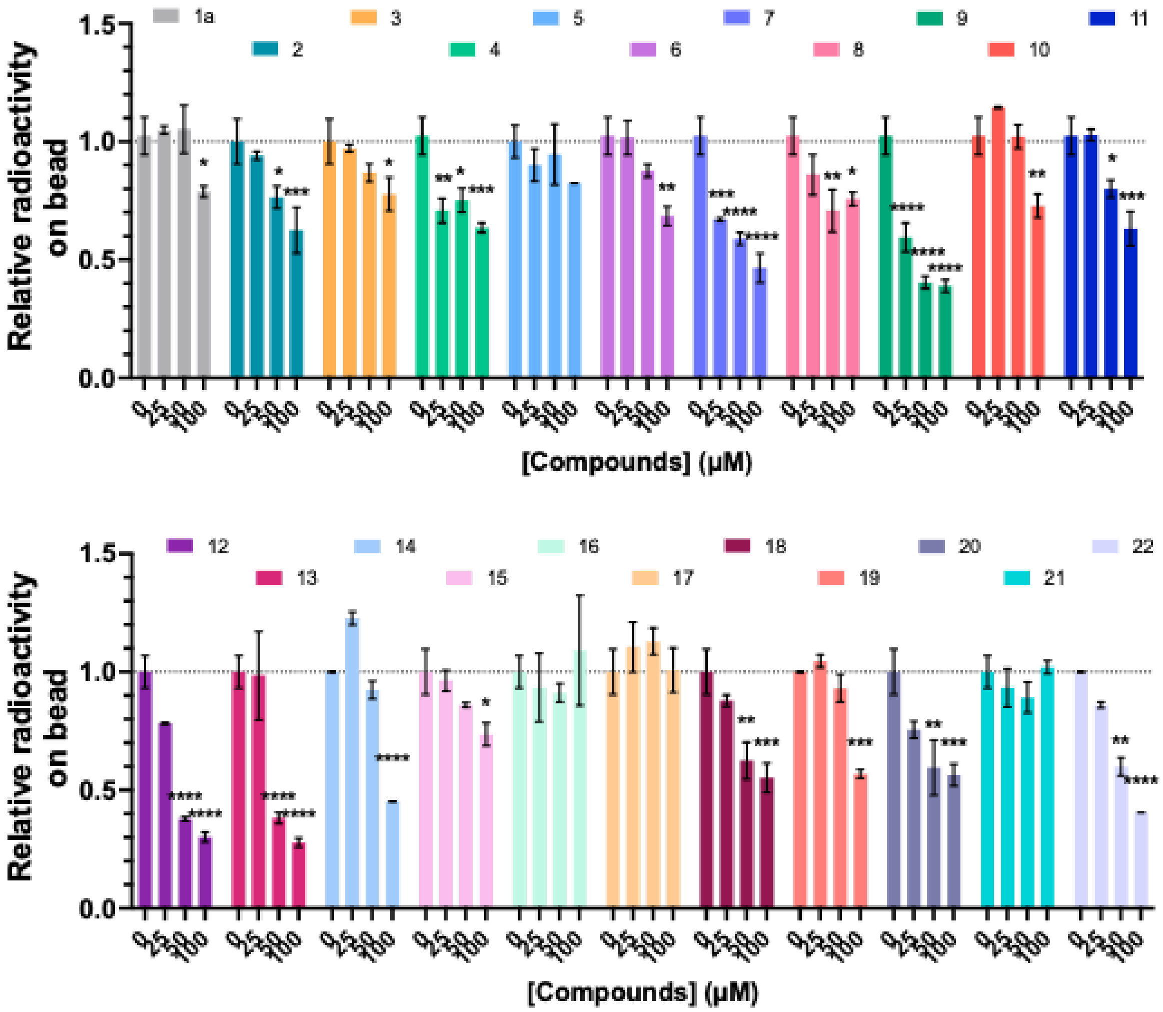


**Figure S2.** Dose response analysis for the competition between **1** and **1a** (positive control) and hit molecules from the primary screen (**2-22**) to bind $^{32}$P-r(CUG)$_{12}$ (n=2). *, $p$ < 0.05; **, $p$ < 0.01; ***, $p$ < 0.001; ****, $p$ < 0.0001; as determined by a One-way ANOVA with multiple comparisons.

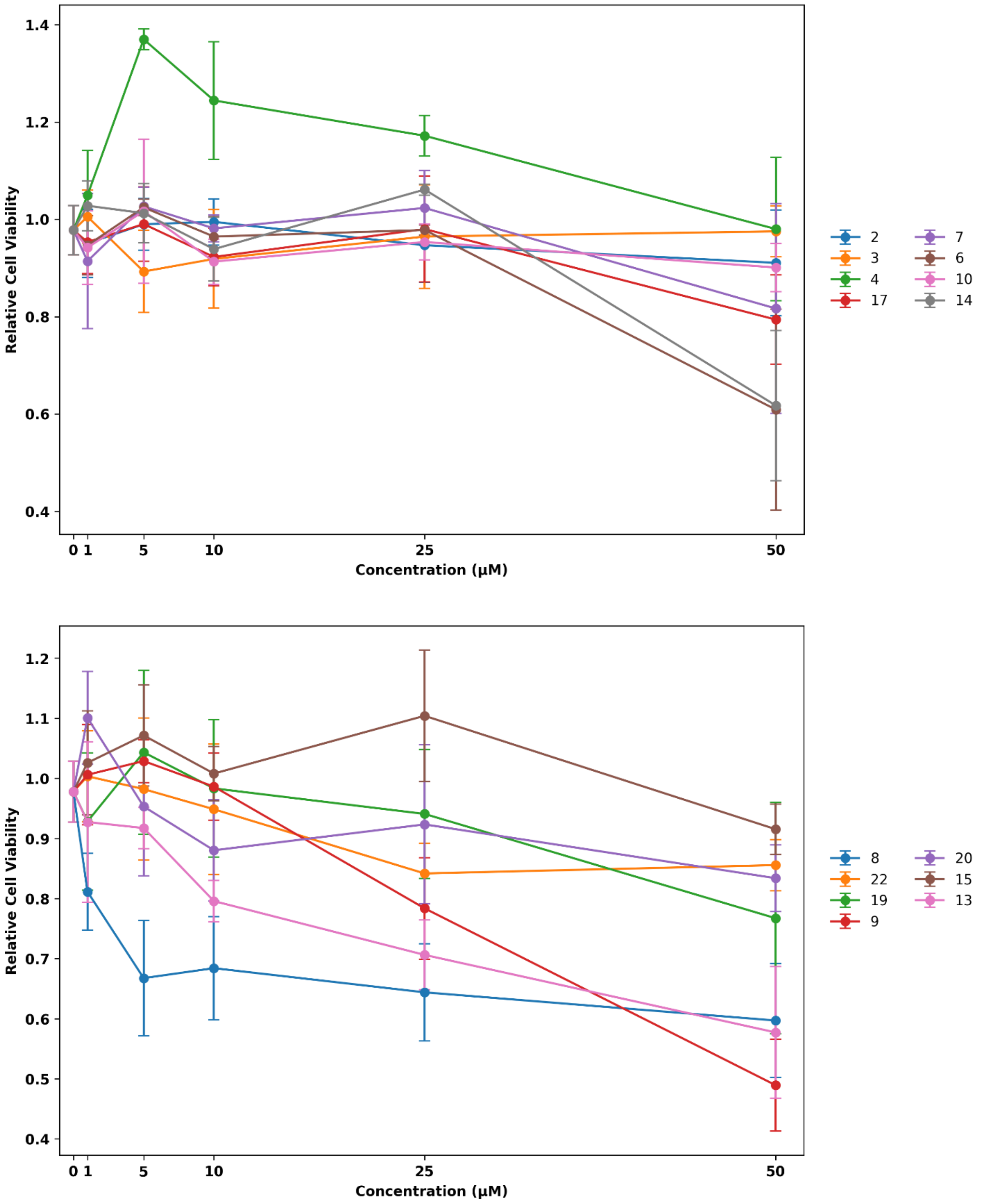


**Figure S3.** Relative viability of C2C12 cells that stably express luciferase fused to a 3' UTR harboring r(CUG)$_{800}$ upon treatment with compounds **2-22** in dose response, up to a maximum concentration of 50 µM, as determined by CellTiter Fluor. Viability was normalized to DMSO (0.1% (v/v) (vehicle)).

A

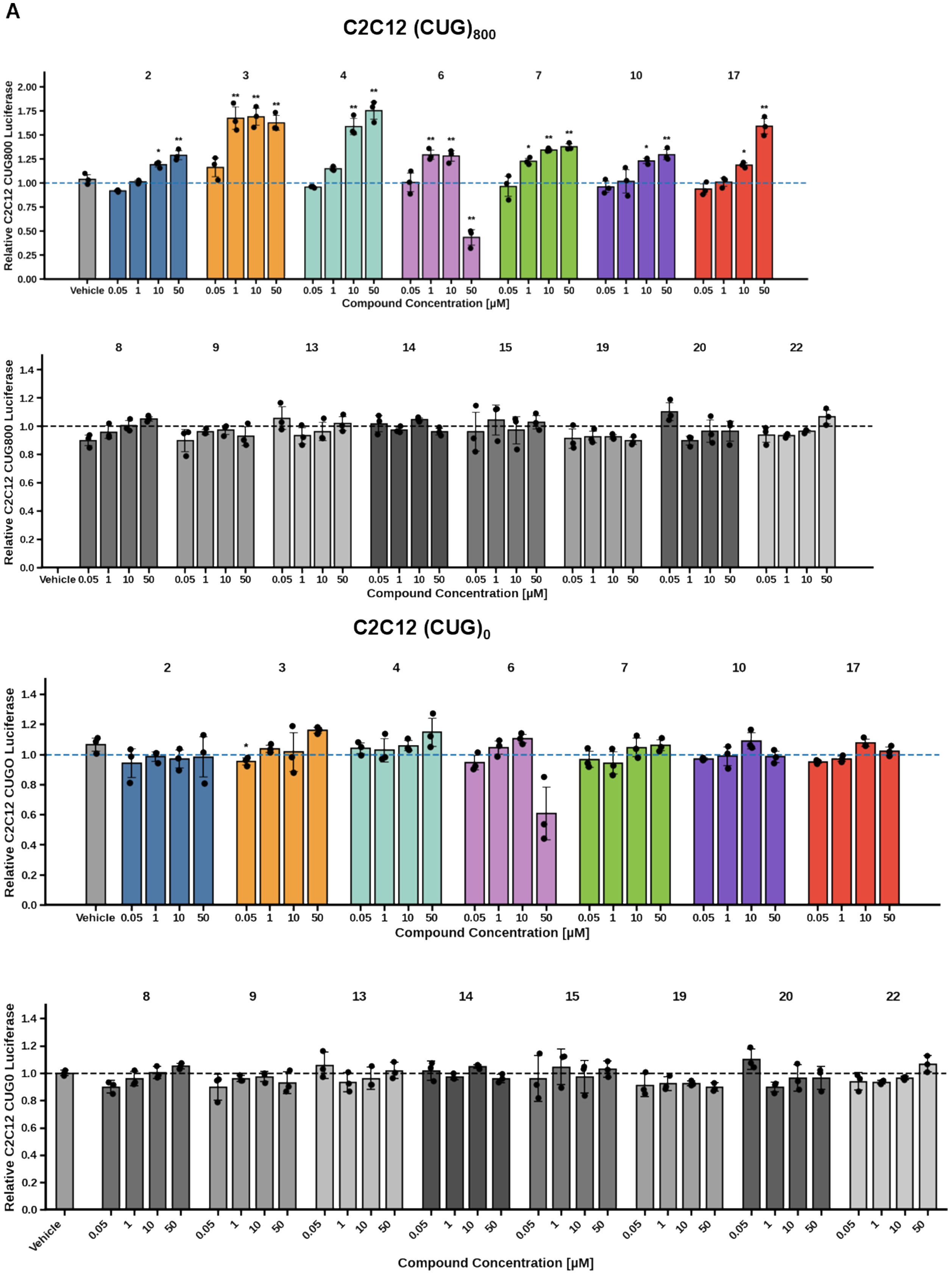
C2C12 (CUG)800
Relative C2C12 CUG800 Luciferase
2 3 4 6 7 10 17
Vehicle 0.05 1 10 50
Compound Concentration [µM]
8 9 13 14 15 19 20 22
C2C12 (CUG)0
Relative C2C12 CUG0 Luciferase

**Figure S4. Cellular evaluation of small molecules that compete with compound 1 for binding to r(CUG) repeats.** (A) Effects of small molecules **2–22** on relative luciferase signal in C2C12 mouse myoblasts stably expressing firefly luciferase fused to a 3′ UTR containing $r(CUG)_{800}$. In this reporter system, sequestration of r(CUG)exp-containing mRNA in nuclear foci suppresses luciferase expression, and compounds that bind the repeat and displace RNA-binding proteins increase luciferase signal by facilitating nucleocytoplasmic transport.(B) Effects of the same compounds on luciferase signal in C2C12 cells expressing luciferase fused to a 3′ UTR containing $r(CUG)_0$, which serves as a control lacking the repeat expansion. Cells were treated with the indicated concentrations and compared to vehicle (0.1% (v/v) DMSO). Data are reported as mean ± SD with individual measurements shown.

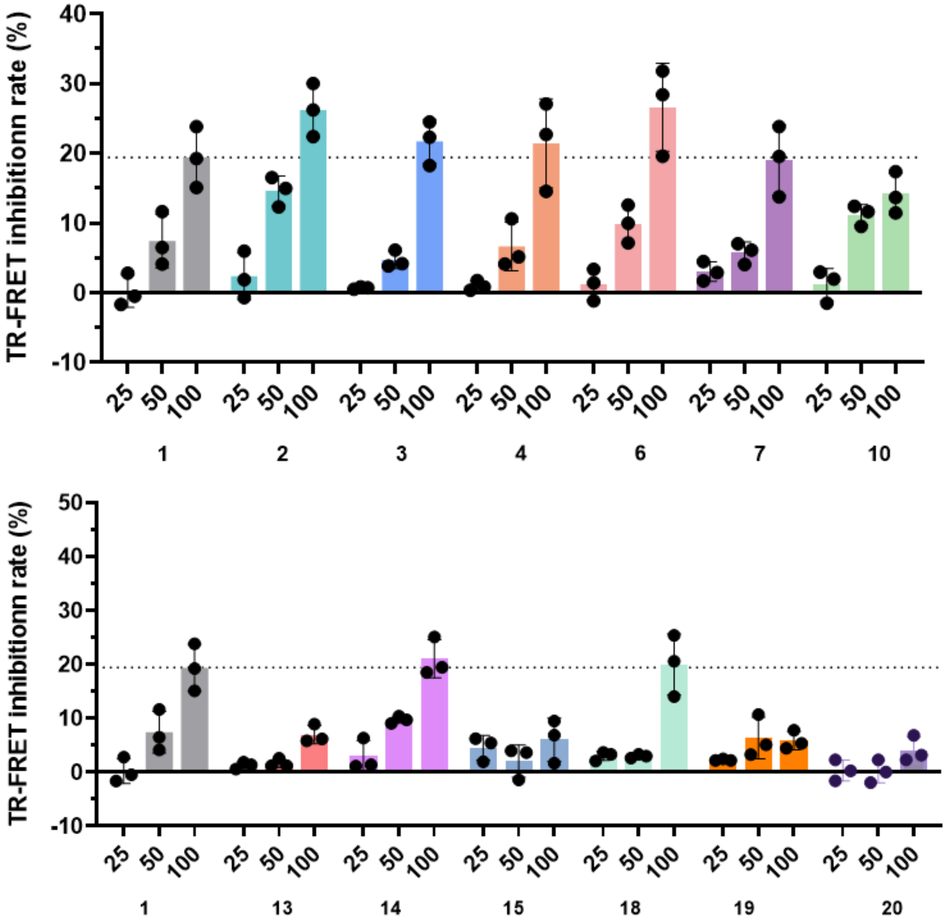


**Figure S5. Dose-dependent disruption of r(CUG)12-MBNL1 complex *in vitro* by small molecules in a TR-FRET assay.** Increasing concentrations of compounds (25 µM, 50 µM and 100 µM) were tested. The percentage disruption of the r(CUG)12-MBNL1 complex was calculated relative to untreated controls. Data are reported as mean ± SD from (n = 3 independent experiments).

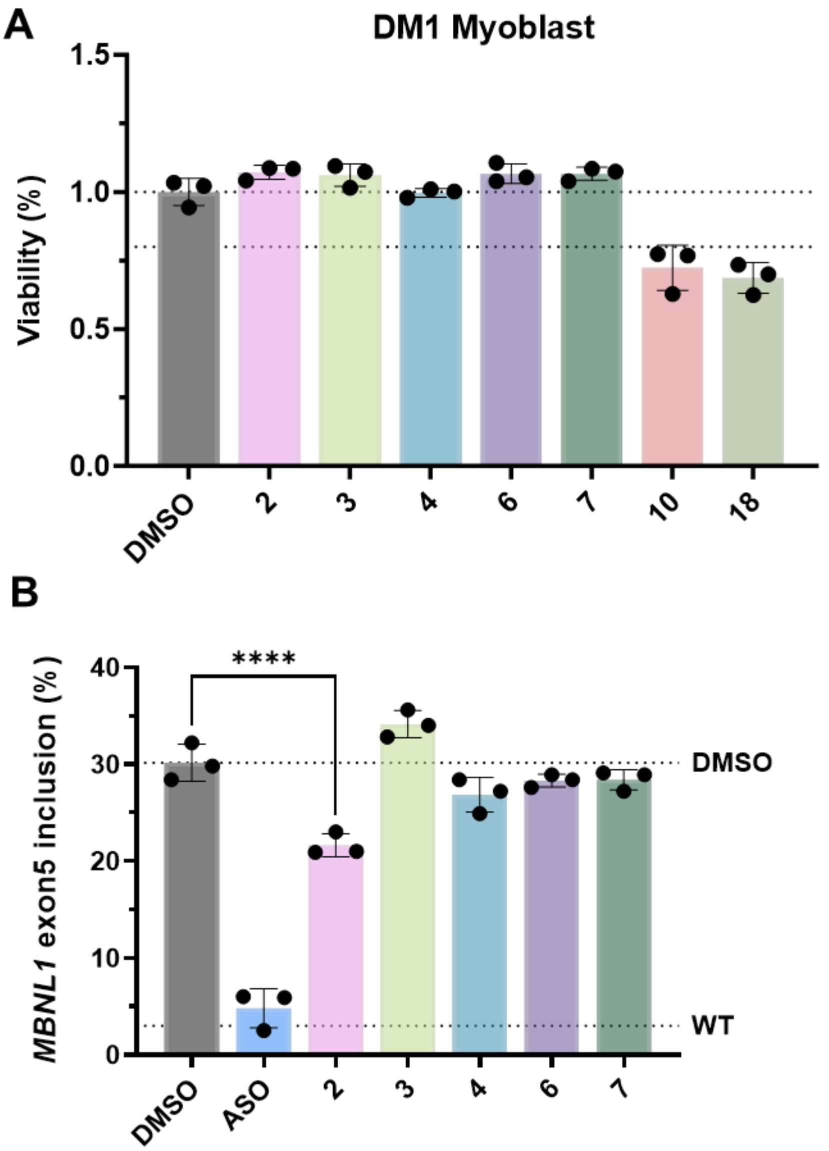


**Figure S6. Dose-dependent disruption of r(CUG)12-MBNL1 complex *in vitro* by small molecules in a TR-FRET assay.** (A) Viability of patient-derived DM1 myoblasts was measured following treatment with 50 µM of the indicated small molecules. Viability was measured using a luminescence-based ATP assay and normalized to the DMSO-treated control. Data are shown as mean ± standard deviation (n = 3 biological replicates). (B) Effect of five compounds (50 µM) without cell toxicity on *MBNL1* exon 5 alternative splicing in DM1 patient-derived myoblasts (n = 3 biological replicates).

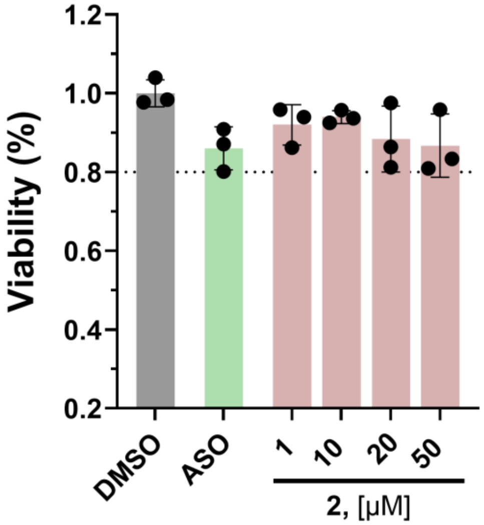


**Figure S7**. **Effect of 2 on the viability of DM1 myoblasts.** The viability of DM1 patient-derived myoblasts following 48 h treatment with **2** was measured using a luminescence-based ATP assay (CellTiter-Glo 2.0) and normalized to the DMSO-treated cells (0.02% (v/v)). Data are reported as mean ± standard deviation (n = 3 biological replicates).

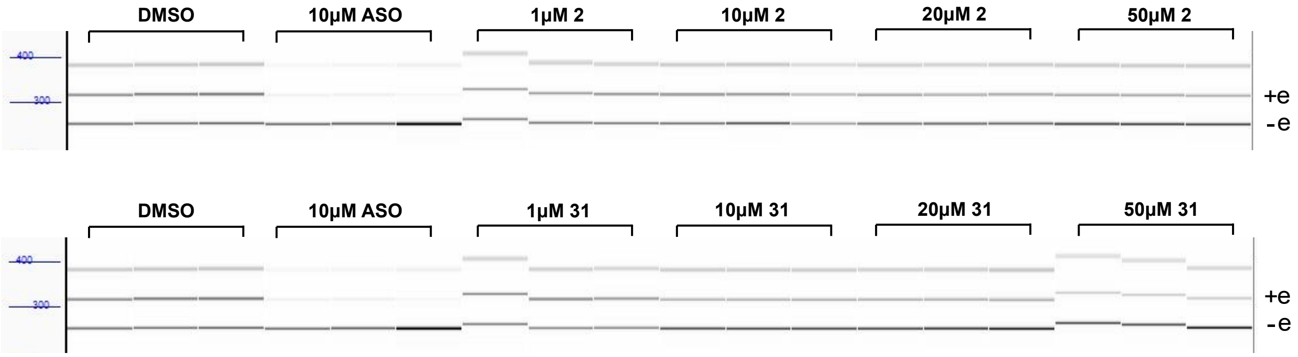


**Figure S8. Compounds 2 and 31 rescue the *MBNL1* exon 5 alternative splicing defect in DM1 patient-derived myoblasts, caused by sequestration of MBNL1 by r(CUG)$^{exp}$.** Effect of **2** and **31** on *MBNL1* exon 5 splicing, as assessed by end-point RT-PCR and fragment analyzer. DM1 patient-derived myoblasts were treated with 1, 10, 20, or 50 µM of each compound. Exon 5 inclusion is indicated by "+ e" while exclusion is indicated by "-e" (n = 3 biological replicates). Quantification of exon 5 inclusion for **2** and **31** is shown in **Figures 3C** and **5D**, respectively.

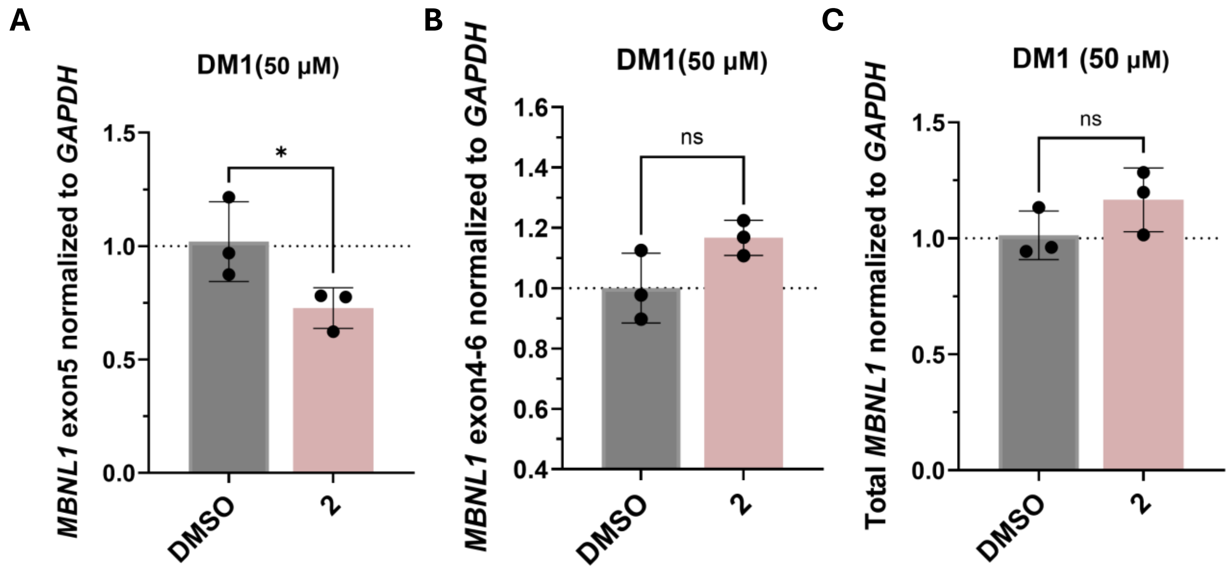


**Figure S9**. **Evaluation of total *MBNL1*, *MBNL1* exon 5, and *MBNL1* exon 4 – exon 6 transcript levels in DM1 patient-derived myoblasts upon treatment with 2**. Effect of compound **2** (50 µM) on the abundance of *MBNL1* exon 5 (A), the exon 4 – exon 6 splicing product (B), and total *MBNL1* transcripts (C) in DM1 myoblasts, each as determined by RT-qPCR (n = 3 biological replicates). Statistical significance was determined relative to untreated controls using a two-tailed unpaired Student's t test with significance thresholds: *, $p < 0.05$; **, $p < 0.01$; ***, $p < 0.001$; ****, $p < 0.0001$.

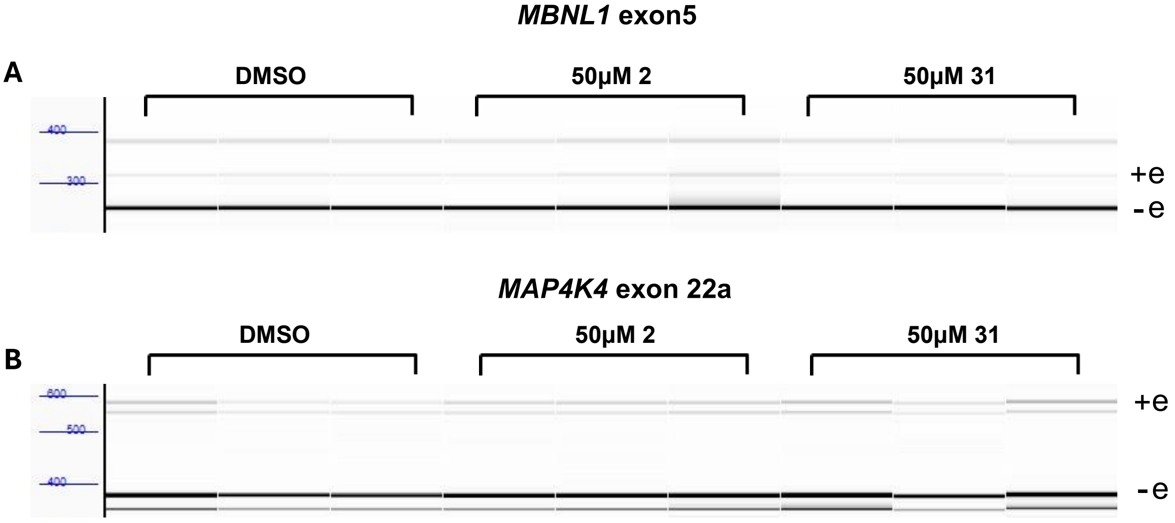


**Figure S10. Effect of 2 and 31 on *MBNL1* exon 5 alternative splicing event WT myoblasts and on a Nova-regulated splicing event in DM1 myoblasts.** (A) Effect of 50 μM of **2** or **31** on *MBNL1* exon 5 splicing in WT myoblasts, as assessed by end-point RT-PCR and separation by fragment analyzer (n = 3 biological replicates). Exon 5 inclusion is indicated by "+ e" while exclusion is indicated by "-e" (n = 3 biological replicates). (B) Effect of 50 μM of **2** or **31** on NOVA-regulated *MAP4K4* exon 22a splicing, [1, 2] as measured by end-point RT-PCR and separation by fragment analyzer (n = 3 biological replicates). Quantification is shown in **Figures 3D** and **6E-F**, respectively.

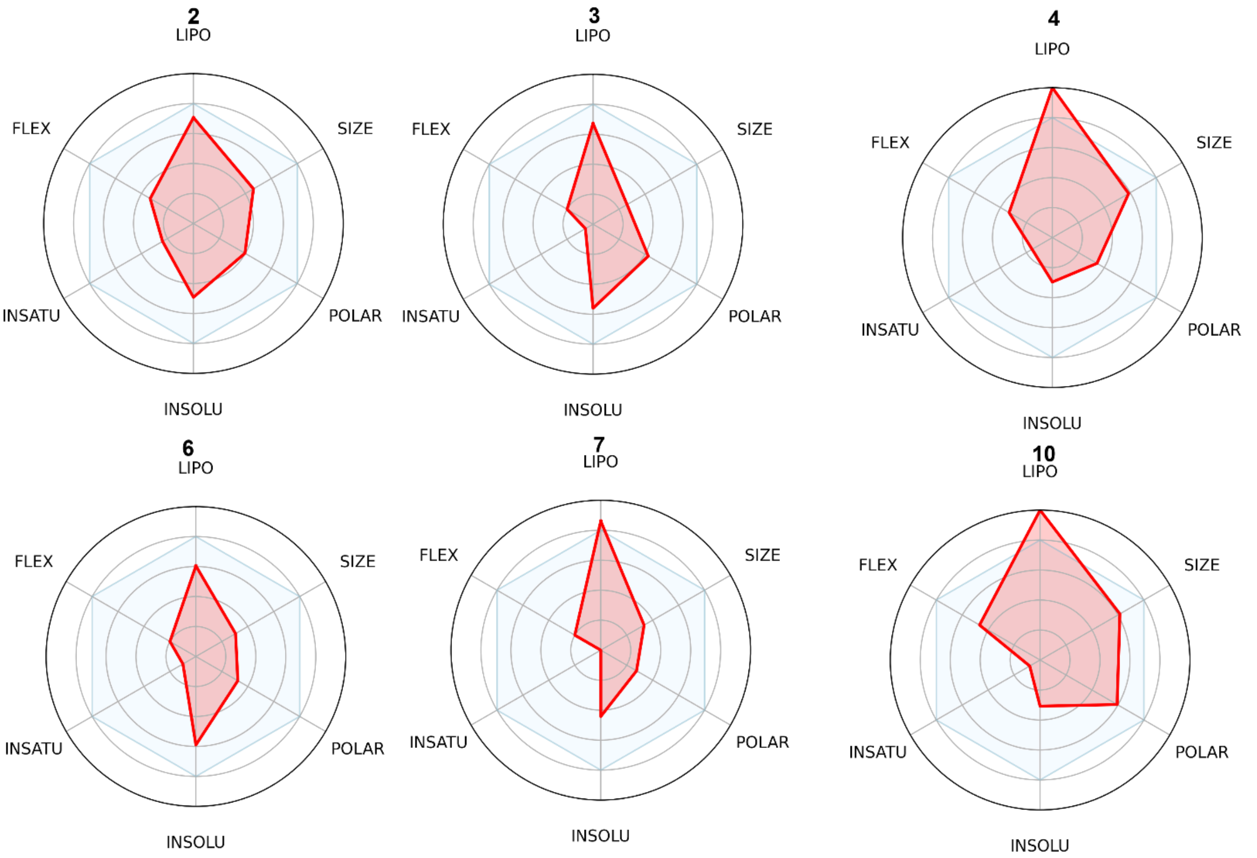


**Figure S11. Physicochemical property analysis of small molecules that alleviate the DM1-associated nucleocytoplasmic transport defect.** Radar plots illustrate the physicochemical profiles of representative compounds. Each plot corresponds to an individual compound (compound numbers shown above each panel). The six axes represent lipophilicity (LIPO), molecular size (SIZE), polarity (POLAR), solubility (INSOLU), saturation (INSATU), and molecular flexibility (FLEX). The pink region denotes the optimal range of physicochemical properties associated with drug-like molecules, as defined by the SwissADME[3] bioavailability radar model. The red polygon represents the calculated properties of each compound. Compounds whose profiles fall largely within the optimal region exhibit physicochemical characteristics consistent with favorable drug-like chemical space. Optimal ranges were defined according to established drug-likeness criteria including Lipinski and Veber rules.

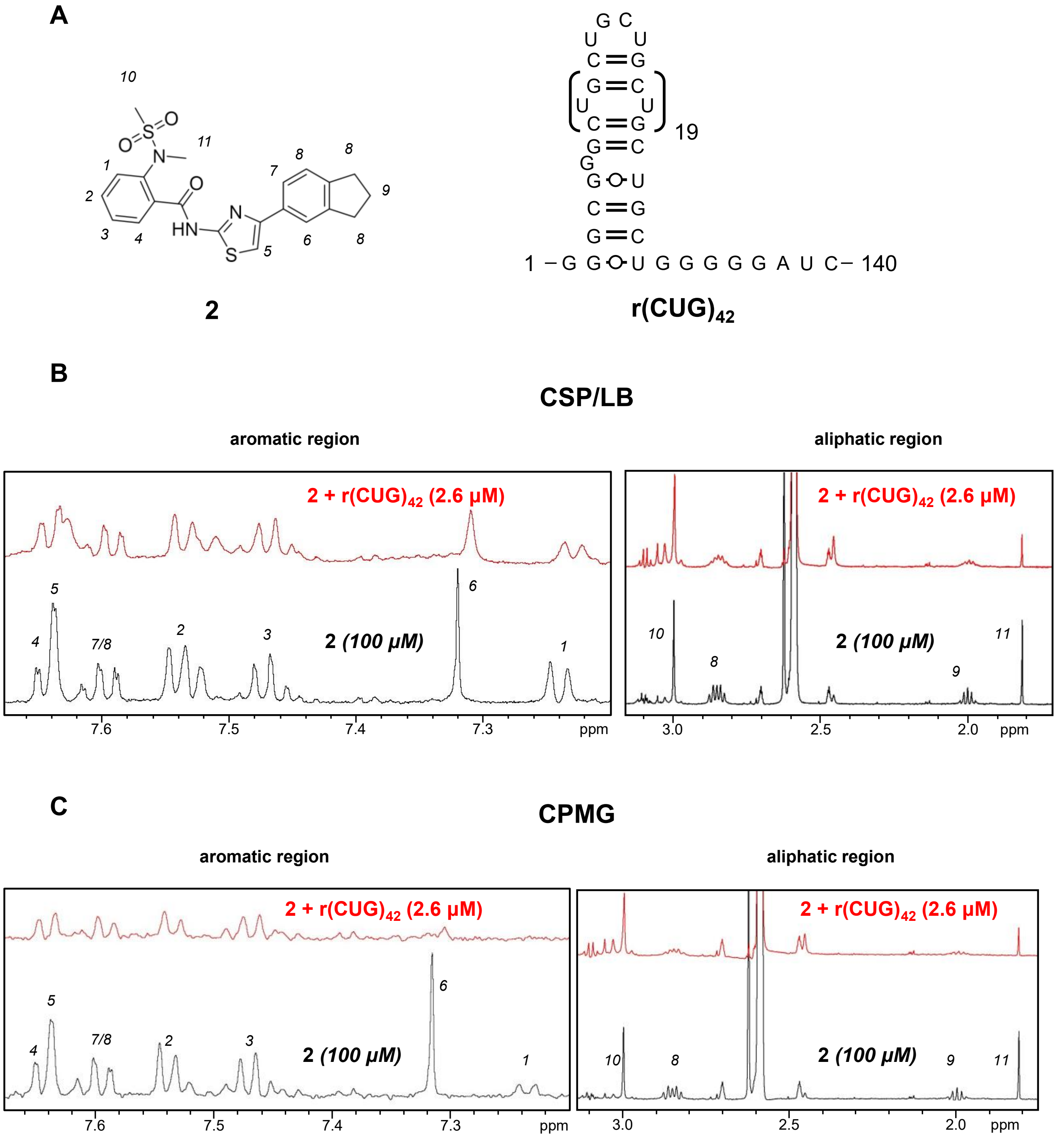


**Figure S12. Compound 2 binds to r(CUG)$_{42}$, as determined by ligand-based NMR experiments**. (A) Chemical structure of **2** and the secondary structure of r(CUG)$_{42}$. Analysis of the aromatic and aliphatic regions of **2** (100 µM) showed CSP/LB and peak reduction in $^1$H standard (B) and CPMG experiments (C) when r(CUG)$_{42}$ was added (red, 2.6 µM RNA or 50 µM internal loops) to the compound (black) at 1:2 internal loops:compound ratio. Particularly, both ligand-based experiments showed consistent greater effect for protons 6 and 1 form the indane and methanesulfonamide moieties, respectively. Buffer is composed of 10 mM $KH_2PO_4/K_2HPO_4$, pH 6.0, 0.05 mM EDTA.

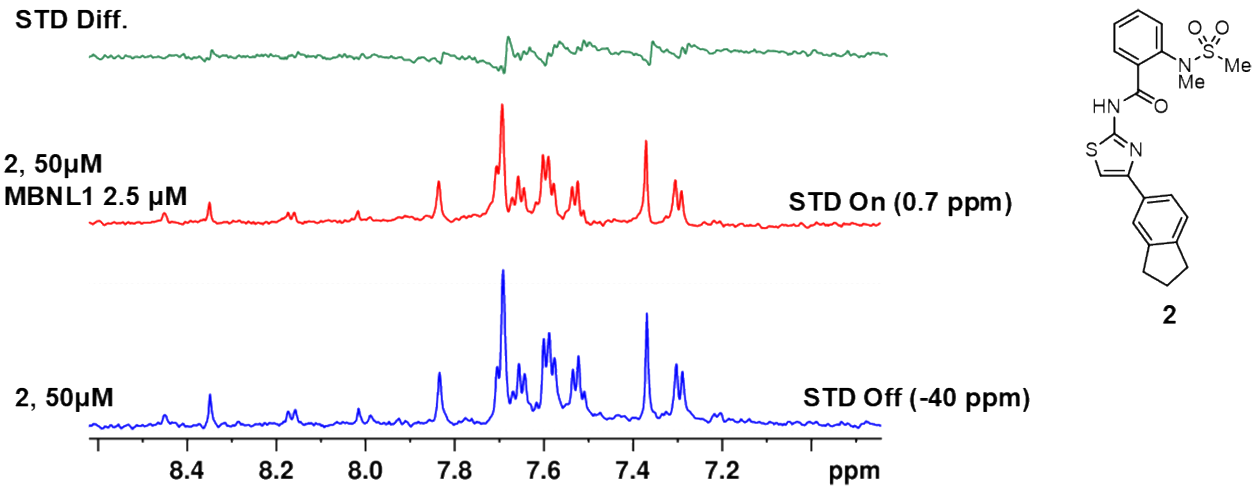


**Figure S13. Saturation transfer difference NMR (STD-NMR) shows no binding of 2 to MBNL1**. STD difference spectrum of **2** upon addition of MBNL1 (green), which is the result of difference (Δ) between STD NMR spectrum of **2** (50 μM) pulsed at an off- resonance (-40 ppm) (blue) and the STD NMR spectrum of **2** (50 μM) and 2.5 μM MBNL1 (red) pulsed at an on-resonance (0.7 ppm) (red). NMR buffer is composed of 5 mM $KH_2PO_4/K_2HPO_4$, 50 mM NaClpH 6.0, 0.05 mM EDTA.

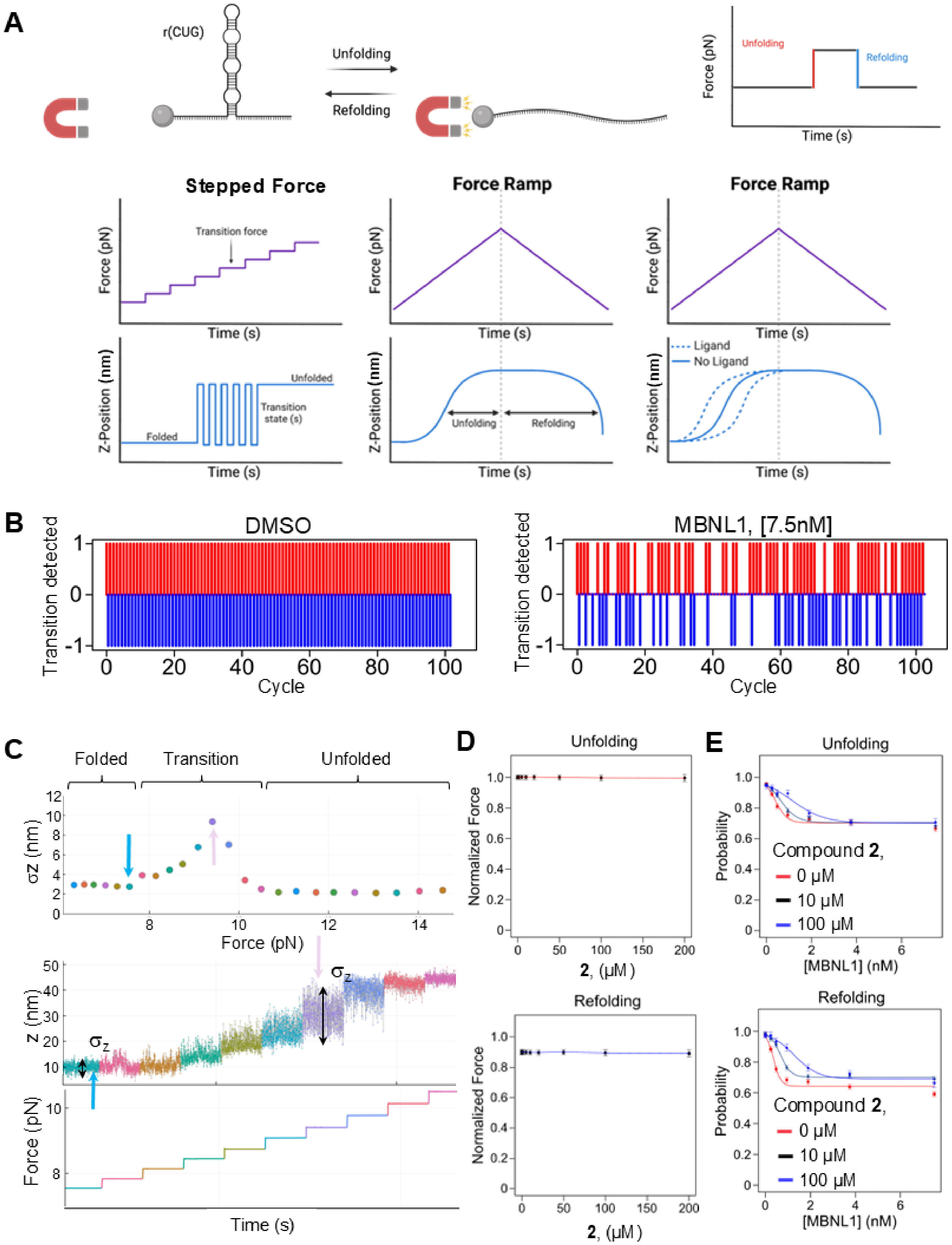
A
r(CUG)
Unfolding
Refolding
Force (pN)
Time (s)
Stepped Force
Force Ramp
Force Ramp
Transition force
Z-Position (nm)
Folded
Unfolded
Transition state (s)
Ligand
No Ligand
B
DMSO
MBNL1, [7.5nM]
Transition detected
Cycle
C
Folded
Transition
Unfolded
σz (nm)
Force (pN)
z (nm)
Time (s)
D
Unfolding
Refolding
Normalized Force
2, (μM)
E
Probability
Compound 2,
0 μM
10 μM
100 μM
[MBNL1] (nM)

**Figure S14. Schematic representation of the MFS technology and supplementary results**. (A) Schematic representation of the two types of MFS single molecule studies performed with expected data plots. (B) Changes in bead Z-position plotted as detected state transitions (folded/unfolded), during repeating cycles of unfolding and refolding of a single $r(CUG)_{21}$ molecule with and without MBNL1 (7.5 nM). Each red bar represents an unfolding event, and each blue bar represents a refolding event. If no event is detected during a cycle, the value is represented as zero. (C) Analysis of the stepped-force experiment for $r(CUG)_{21}$ folding/unfolding events. Top: plot of the change in Z-position (nm) with folding and transition states identified at varying force (pN). Middle: plot of the change in Z-position (nm) of each force step during the transition with the peak of the variance representing the transition state force for that RNA molecule. Bottom: plot of the force(pN) of each force step during the transition state as a function of time. The force was held constant at each step for 30 seconds, with each color representing a discrete force step. (D) Median normalized unfolding and refolding force of each RNA with increasing compound **2** concentration (n=249). The median force for all molecules and SEM is shown. (E) The unfolding and refolding probability upon MBNL1 protein binding in the presence of 0, 10 and 100 µM of compound **2**, using ramp experiments (n=255, 198, 148 for 0, 10, 100 µM of 2, respectively). The probability for each molecule is calculated and the medians with SEM of all molecules are shown.

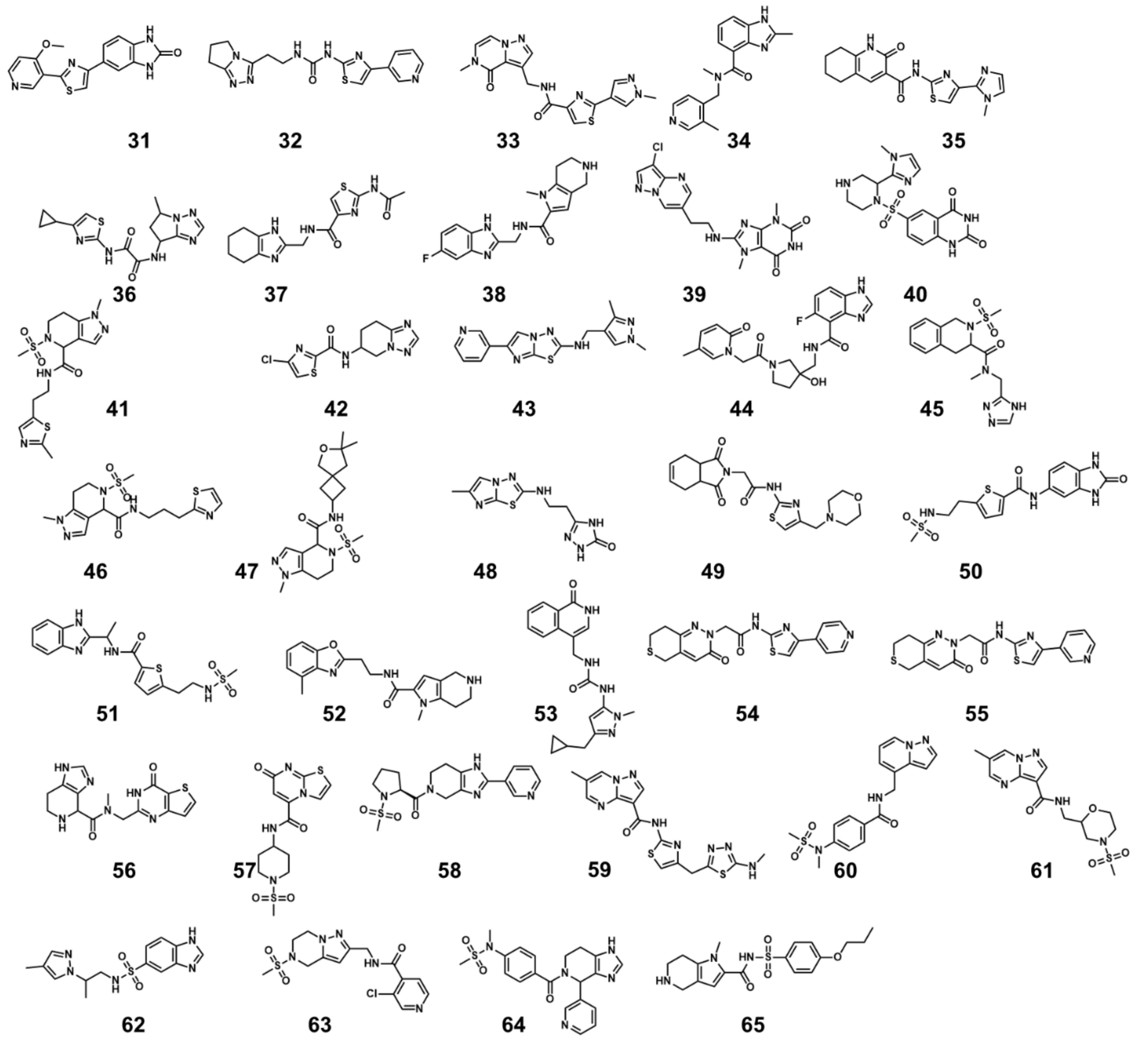


**Figure S15**. Chemical structures of 35 compounds derived from 2 through PAS-logS-similarity search process (refer to the main text for workflow explanation).

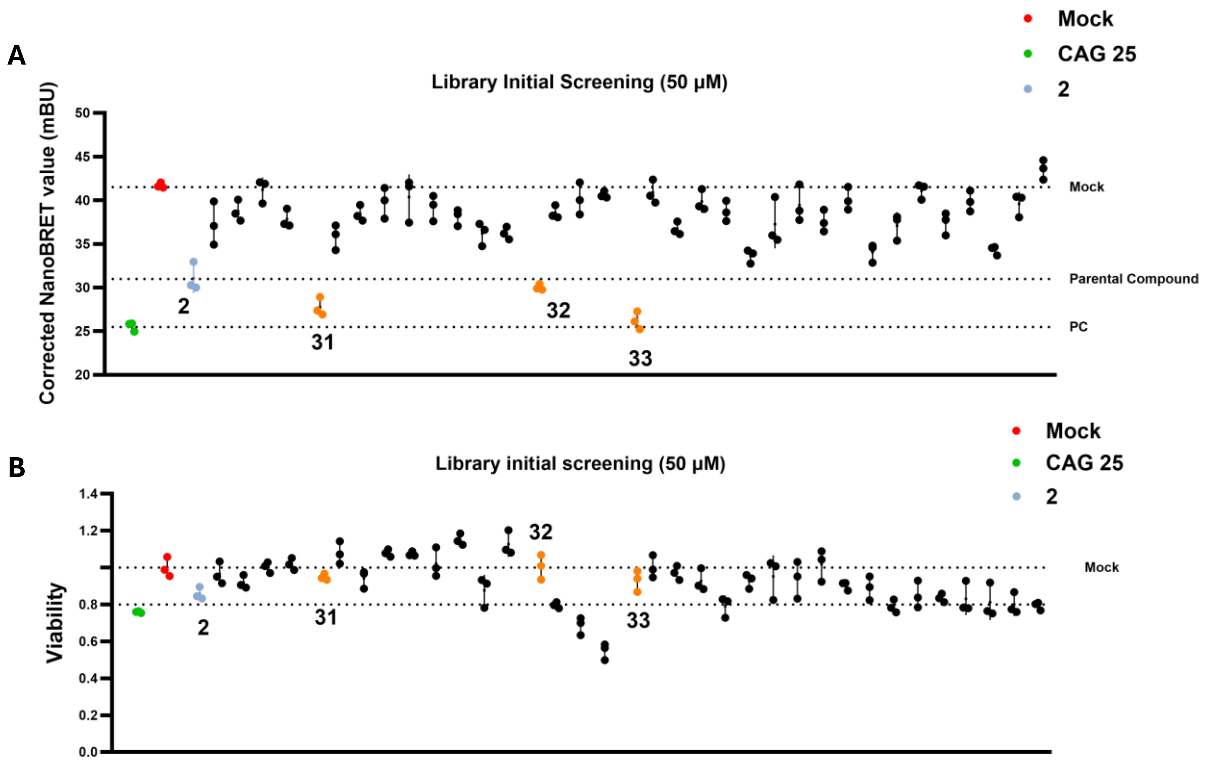


**Figure S16. High-throughput screening of small molecules for inhibition of r(CUG)$^{exp}$-MBNL1 complex formation in the NanoBRET assay and effect of the small molecules on viability.** (A) Small molecules were screened in triplicate in the HeLa480 NanoBRET assay at a concentration of 50 μM to identify inhibitors of the r(CUG)$^{exp}$-MBNL1 complex. The corrected NanoBRET values (NanoBRET signal of treated samples minus NanoBRET signal of control) are plotted for all screened compounds. Dashed lines represent the activity of **2**. Compounds that decreased the NanoBRET signal to a greater extent than parent compound **2** are highlighted in orange. Mock-treated controls (red) and CAG25 Vivo-Morpholino-treated cells (green) are included for comparison. (B) Cell viability assay performed in HeLa480 cells to assess cytotoxicity of small molecules identified from the NanoBRET screening. Small molecules were tested at 50 μM, and cell viability was measured using a luminescence-based ATP assay. Viability percentages are normalized to DMSO-treated cells (0.1% (v/v); red). The viability of cells treated with 10 μM of CAG25 Vivo-Morpholino is indicated in green. Dashed lines are present at 100% and 80% viability.

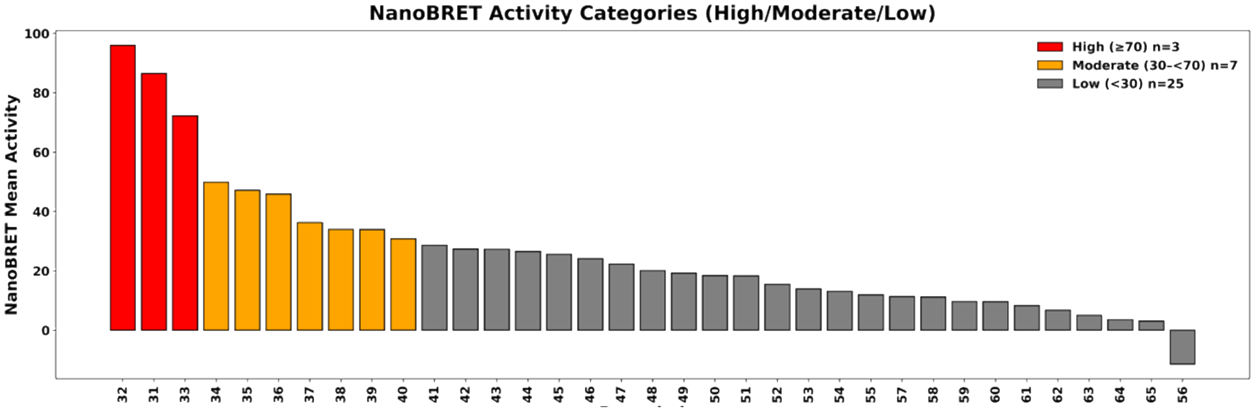


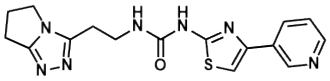

32

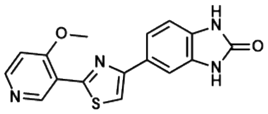

31

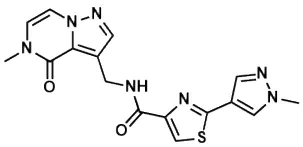

33

**Figure S17**. **Quantitative comparison of NanoBRET activity across 35 PAS-logS-similarity improved compounds.** Bar plot showing NanoBRET engagement values for a subset of similarity-derived compounds, ordered from highest to lowest activity. Compounds in the high-activity class are highlighted in red, while intermediate-activity compounds are shown in orange.

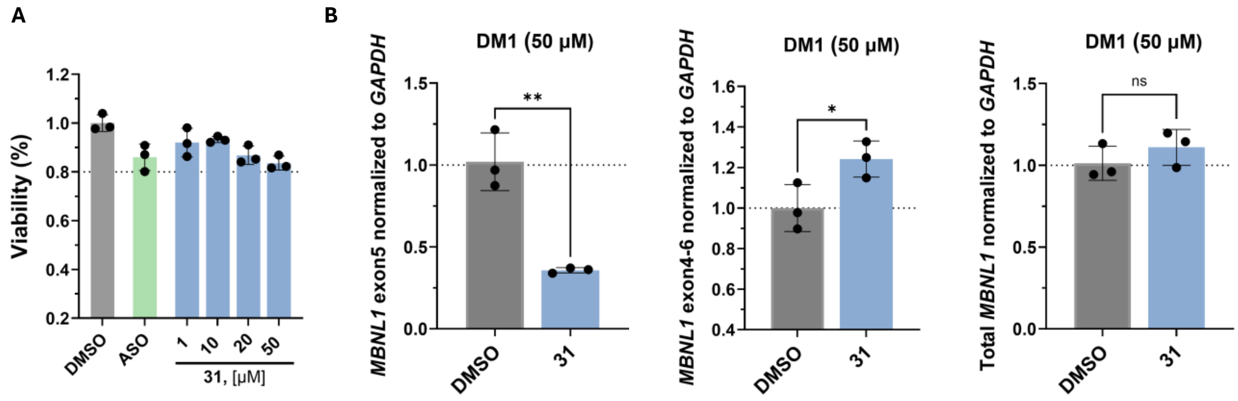


**Figure S18**. **Evaluation of viability, total *MBNL1*, *MBNL1* exon 5 and *MBNL1* exon 4 – exon 6 transcript levels in DM1 patient-derived myoblasts upon treatment with 31**. (A)Viability of DM1 patient-derived myoblasts following 48 h treatment with **31**. Viability was normalized to the DMSO-treated cells (0.1% (v/v)). Data are reported as mean ± standard deviation (n = 3 biological replicates). (B) Effect of **31** (50 μM) on the abundances of *MBNL1* exon 5, exon 4 – exon 6 splicing product, and total *MBNL1* transcripts in DM1 myoblasts, each as determined by RT-qPCR (n = 3 biological replicates). Statistical significance was determined relative to untreated controls using a two-tailed unpaired Student's t test with significance thresholds: *, $p < 0.05$; **, $p < 0.01$; ***, $p < 0.001$; ****, $p < 0.0001$.

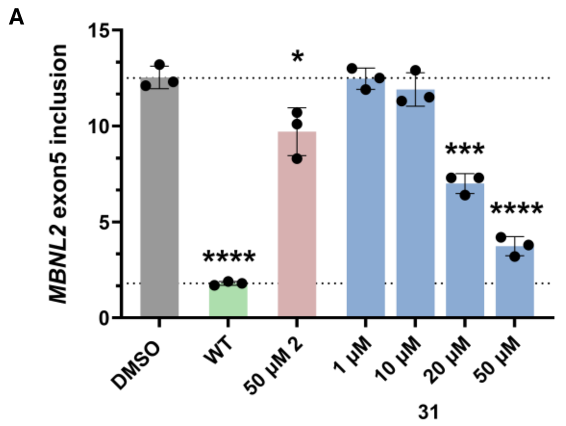


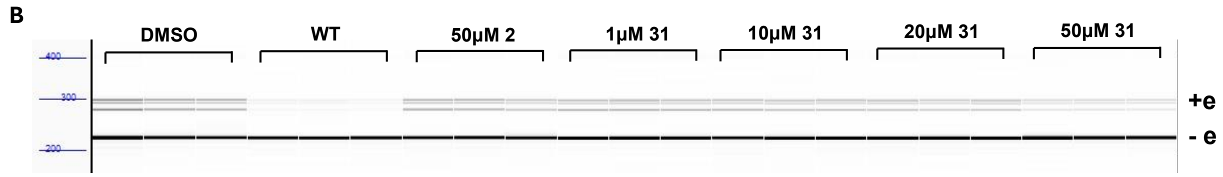


**Figure S19**. **Compound 31 rescues the *MBNL2* exon 5 splicing defects in DM1 patient-derived myoblasts.** (A) Quantification of the fragment analyzer traces to study rescue of the *MBNL2* exon 5 splicing defects. Splicing was assessed by end-point RT-PCR followed by separation of the splicing products by fragment analyzer (n = 3 biological replicates). Statistical significance was determined by a One-way ANOVA with multiple comparisons with significance thresholds: *, $p < 0.05$; **, $p < 0.01$; ***, $p < 0.001$; and ****, $p < 0.0001$. (B) Representative fragment analyzer trace for the rescue of the MBNL2 exon 5 splicing defects in DM1 myoblasts by **31** (1– 50 µM). Exon 5 inclusion is indicated by "+e" while exclusion is indicated by "-e".

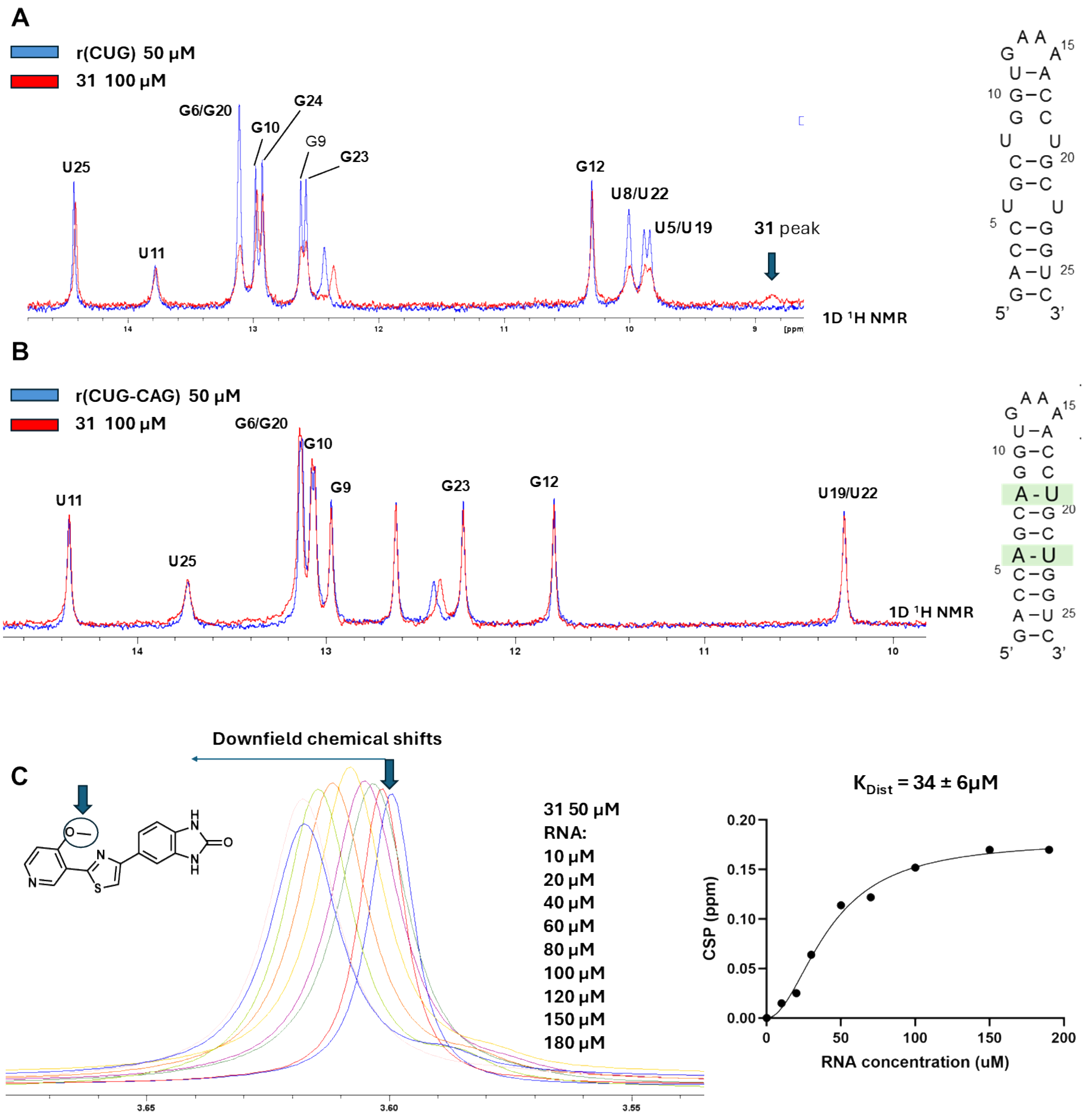


**Figure S20. Imino titration of r(CUG)$_2$ shows specificity of 31 for the U/U internal loops present in r(CUG)$^{exp}$.** (A) Imino $^1$H spectra of r(CUG)$_2$ upon addition of **31**. Exchange broadening of the U5/U19, U8/U22 confirms that **31** engages the U/U internal loops. (B) To further assess specificity, changes in the imino proton spectra of a fully paired RNA as a function of **31** concentration were also measured. No exchange broadening was observed for the base paired construct confirming the specificity of **31**. (C) Titration of r(CUG)$_2$ into a constant concentration of **31** (50 µM) to enable $K_{Dist}$ measurements. The downfield chemical shifts of the O-methyl proton (blue arrow) at 3.6 ppm was used to measure the $K_{Dist}$. Data fitting was performed with Prism one binding site model.

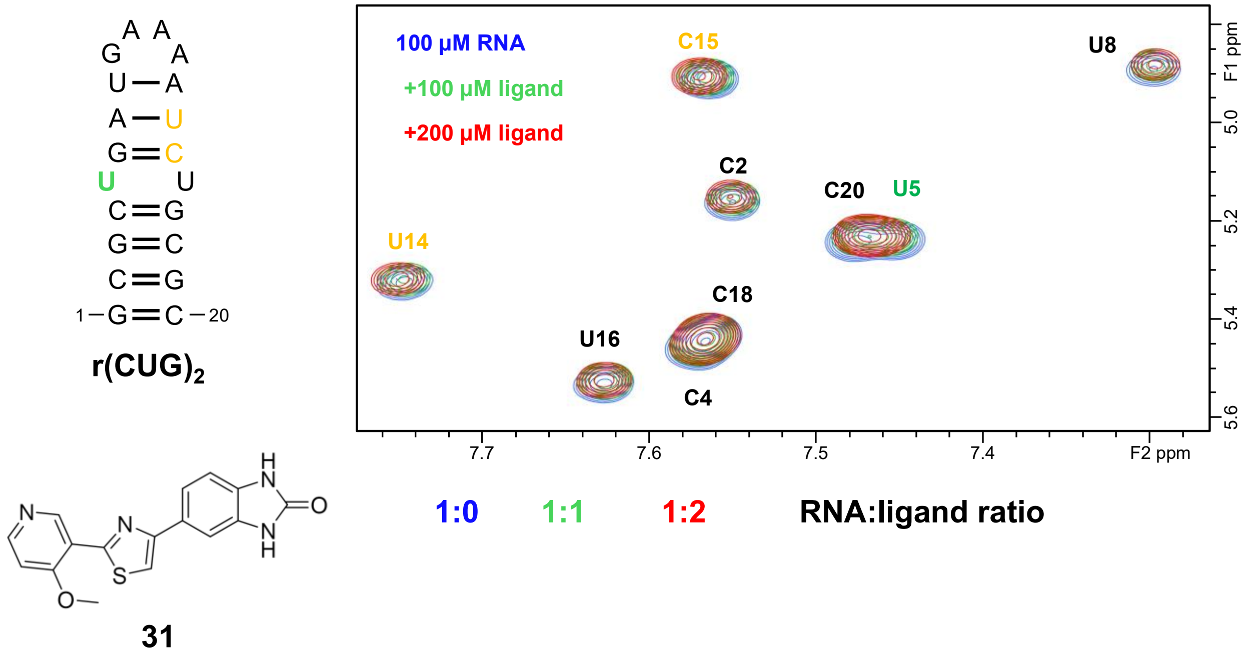


**Figure S21. Interaction of 31 and r(CUG)$_2$ monitored by a 2D TOCSY NMR experiment.** The pyrimidine H5-H6 region of the TOCSY spectrum (60 ms mixing time, 298K) of r(CUG)$_2$ (blue) is superimposed on the spectra of complexes with increasing RNA:ligand molar ratios (color-coded as indicated in the graph). On the right, the nucleotides mainly involved in the interactions are color coded in green when CSP and broadening of the cross peak was observed, and in yellow when only CSP was observed. Buffer is composed of 10 mM $KH_2PO_4$/$K_2HPO_4$, pH 6.0, 0.05 mM EDTA.

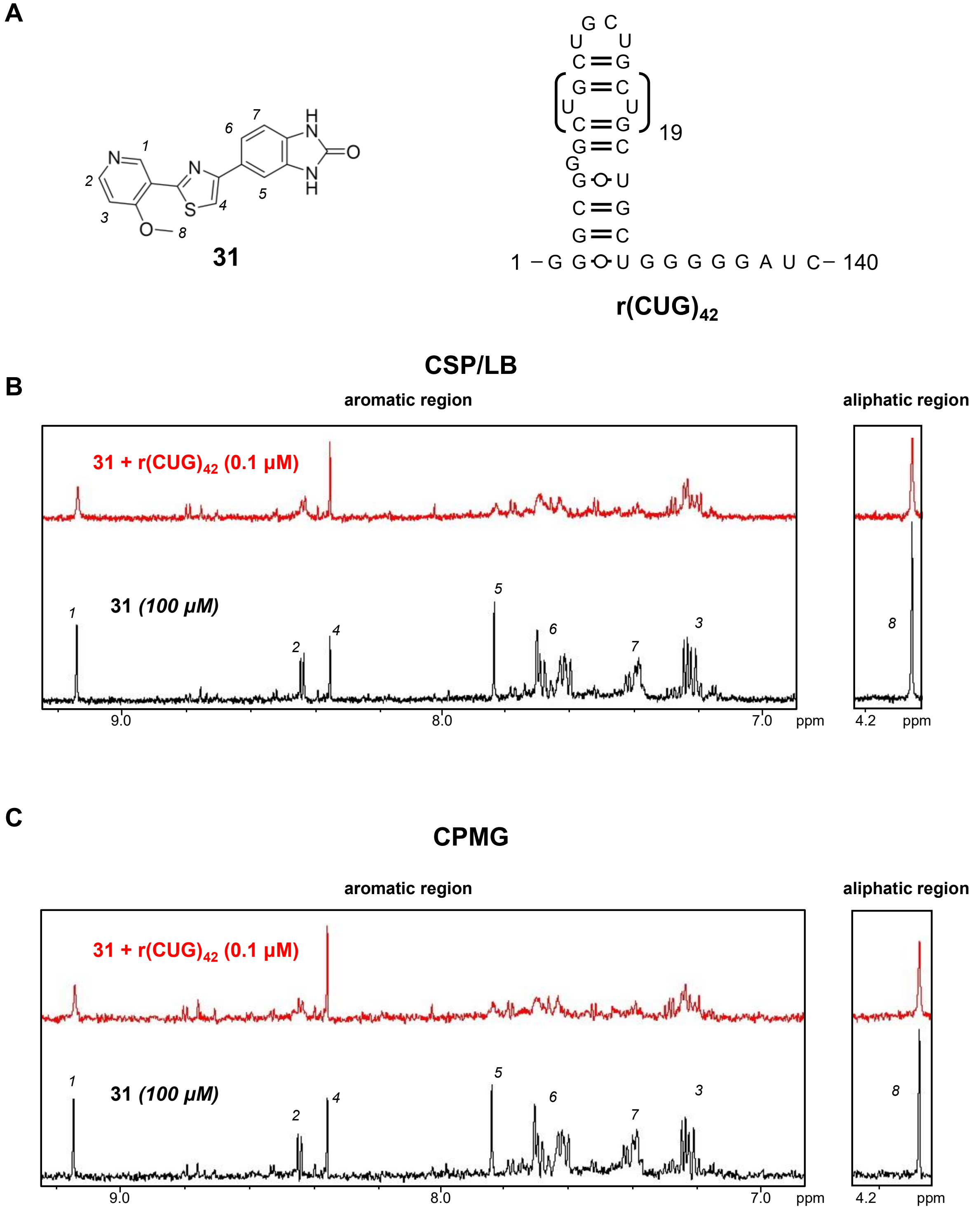


**Figure S22. Compound 31 binds to longer r(CUG) repeats, as assessed by ligand-based NMR experiments.** (A) Chemical structure of **2** and the secondary structure of $r(CUG)_{42}$. (B) Analysis of the aromatic region of **31** showed CSP/LB and peak reduction in 1D $^1H$ spectrum upon addition of $r(CUG)_{42}$ was added (red) as compared to compound alone (black). (C) Likewise, CPMG studies show changes in the relaxation time of various protons upon addition of r(CUG)$_{42}$ (red), indicative of binding. Both ligand-based experiments showed greater effect on protons 5 and 7 from the benzimidazolinone ring, suggesting its importance in binding to r(CUG)$_{42}$. Buffer is composed of 10 mM $KH_2PO_4/K_2HPO_4$, pH 6.0, 0.05 mM EDTA.

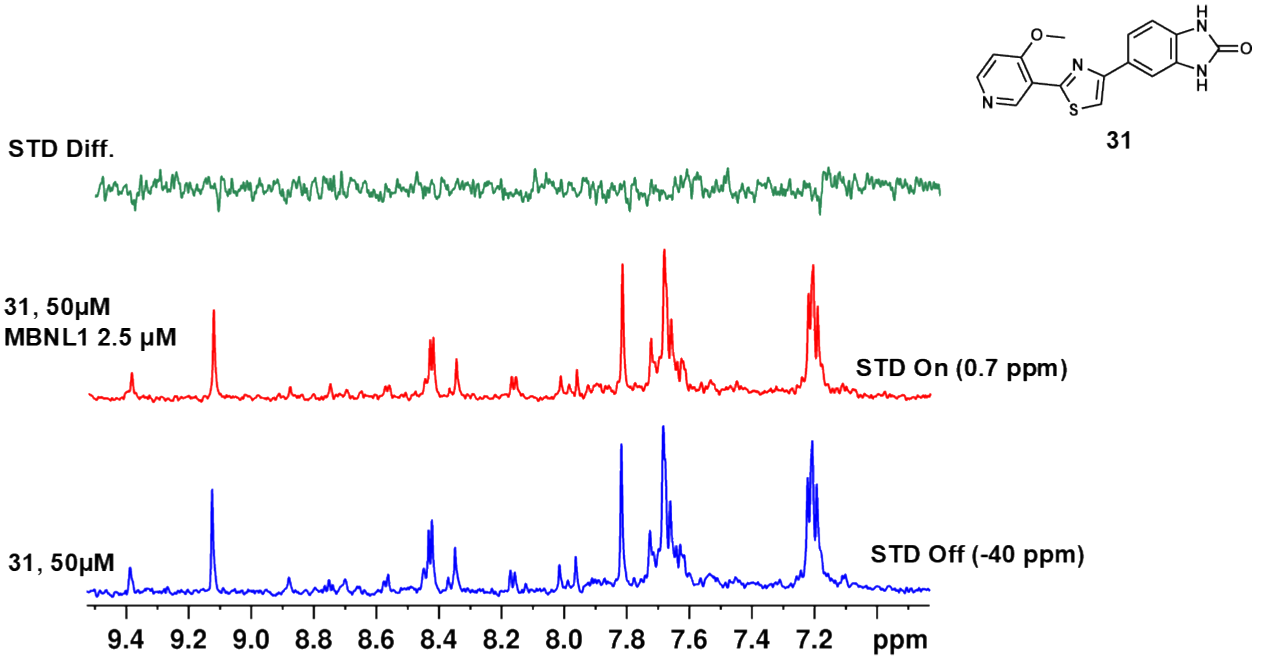


**Figure S23. Compound 31 does not bind to MBNL1, as determined by saturation transfer difference NMR (STD-NMR).** STD difference spectrum of **2** upon addition of MBNL1 (green), which is the result of difference (Δ) between STD NMR spectrum of **31** (50 μM) pulsed at an off- resonance (-40 ppm) (blue) and the STD NMR spectrum of **31** (50 μM) and 2.5 μM MBNL1 (red) pulsed at an on-resonance (0.7 ppm) (red). NMR Buffer 5 mM $KH_2PO_4/K_2HPO_4$, pH 6.0, and 50 mM NaCl.

A

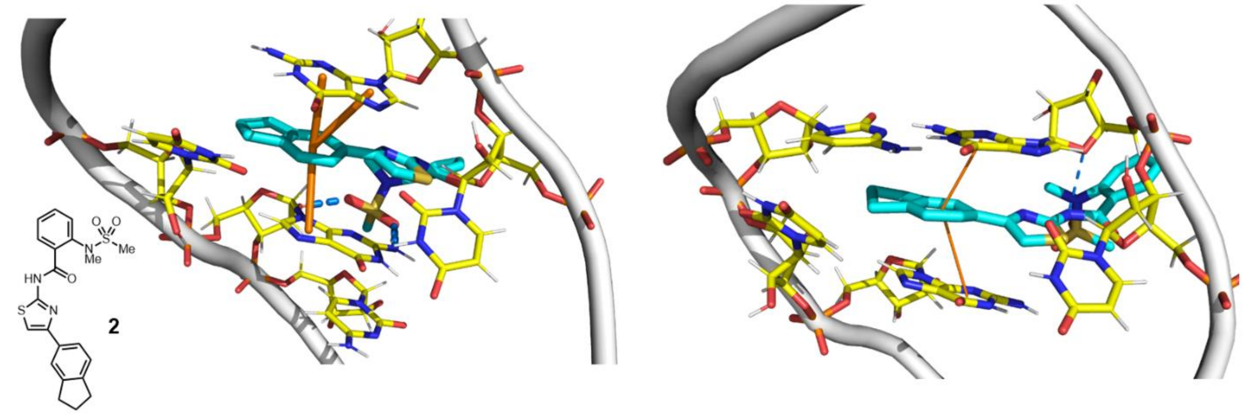


B

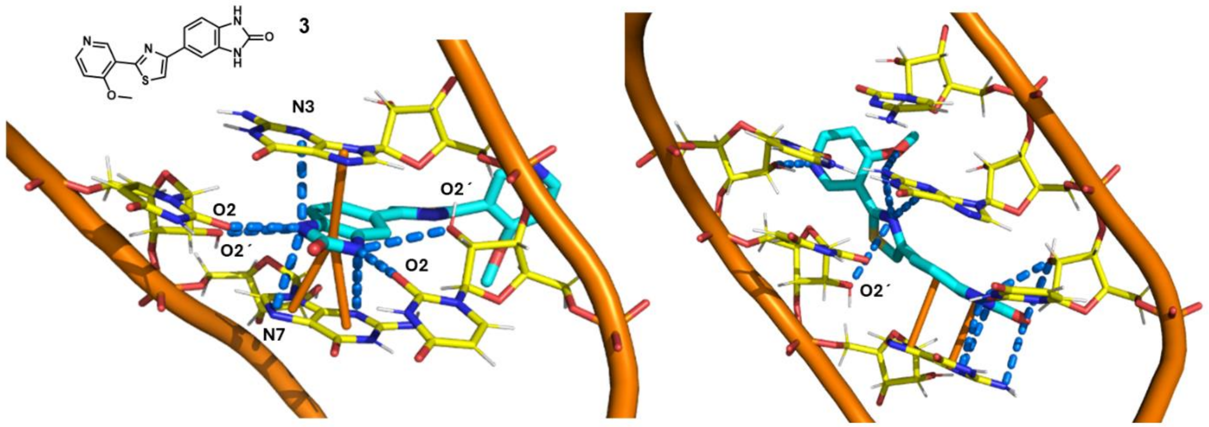


**Figure S24. Distinct binding modes of compounds 2 and 31 in r(CUG) repeat RNA revealed by molecular dynamics (MD) simulations.** (A) Structural model of compound 2 (cyan sticks) bound to an r(CUG) repeat RNA internal loop. The RNA backbone is shown in grey and bases are depicted as yellow sticks. Compound 2 occupies the widened UU internal loop and aligns along the helical axis, forming limited stabilizing contacts with the surrounding nucleotides. The aromatic portion of the ligand engages in π–stacking interactions with adjacent bases, while polar functional groups form occasional hydrogen-bond interactions with nearby nucleobase heteroatoms and backbone phosphate groups (orange lines). However, compared to compound 31, these contacts are less extensive and less geometrically constrained, resulting in fewer direct hydrogen-bond interactions with the uridine O2/O2′/N3 atoms that define the UU loop pocket. Consequently, compound 2 interacts more loosely with the internal loop and closing base pairs. (B) Structural model of compound 31 (cyan sticks) bound to a r(CUG) repeat RNA internal loop. The RNA backbone is shown as orange ribbons, and bases involved in interactions with the ligand are displayed as yellow sticks. Compound 31 engages the loop through a network of hydrogen-bond contacts (blue dashed lines) centered on uridine O2, O2′, and N3 atoms, while simultaneously maintaining π-stacking interactions between the ligand heterocycles and neighboring bases. Two alternate orientations are shown, illustrating how compound 31 adopts a compact pose within the loop and establishes a consistent hydrogen-bonding geometry across both faces of the pocket.

**Table S1**. Murcko scaffolds extracted from the parent library with 400,000 compounds showing uniqueness of ~80%. Representative structures from each group of scaffolds are shown along with the number of times each scaffold is observed in the library.

| **Scaffold (redundancy)** | | | |
|---|---|---|---|
| 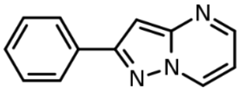 (110) | 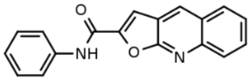 (108) | 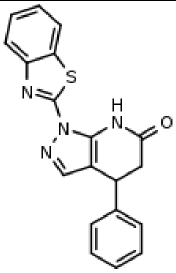 (103) | 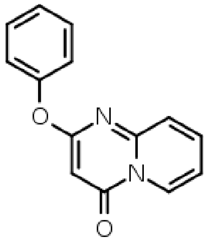 (101) |
| 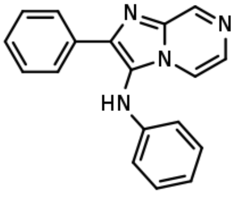 (100) | 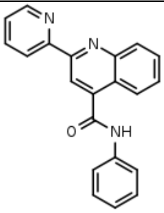 (98) | 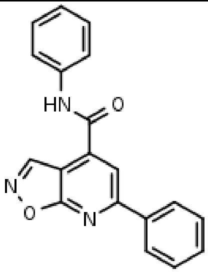 (96) | 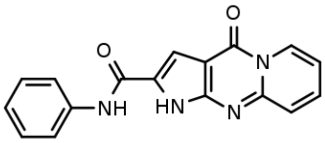 (94) |
| 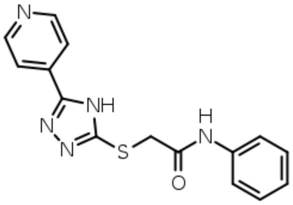 (89) | 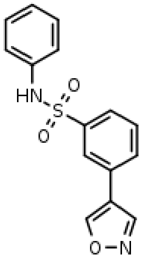 (87) | 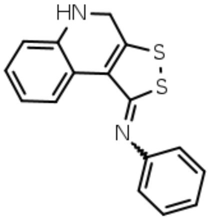 (80) | 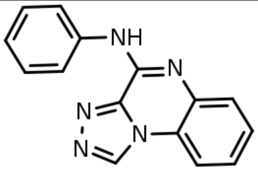 (78) |

| | | | |
|---|---|---|---|
| 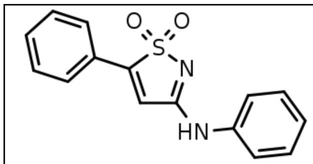 (75) | 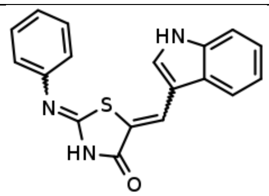 (71) | 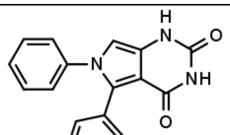 (70) | 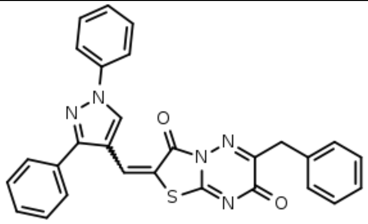 (66) |
| 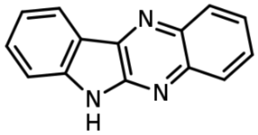 (60) | 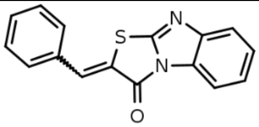 (55) | 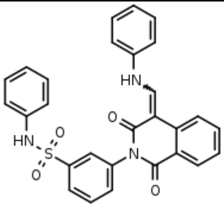 (53) | 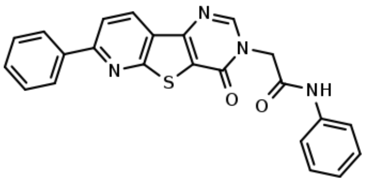 (51) |
| 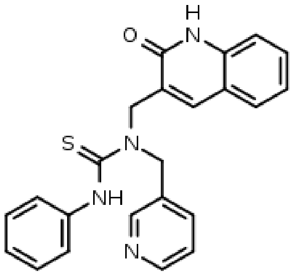 (47) | 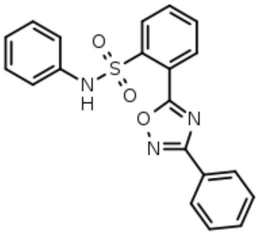 (40) | 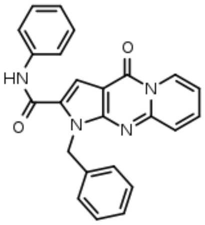 (38) | 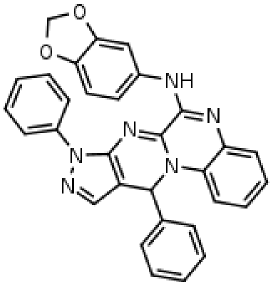 (35) |
| 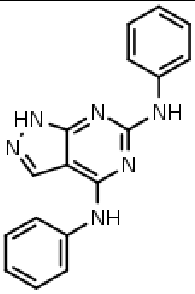 (32) | 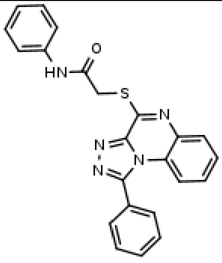 (29) | 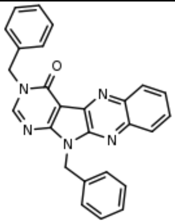 (28) | 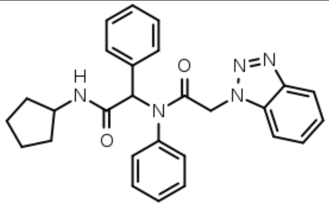 (25) |

| | | | |
|---|---|---|---|
| 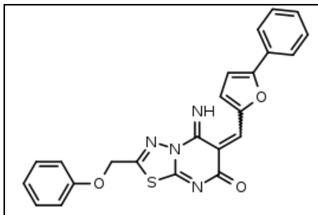 (20) | 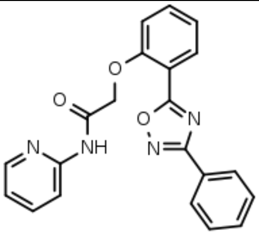 (15) | 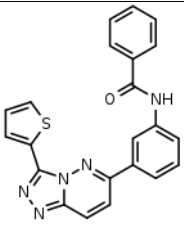 (10) | 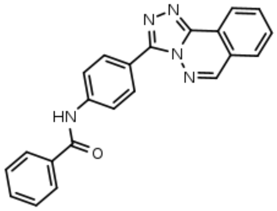 (5) |
| 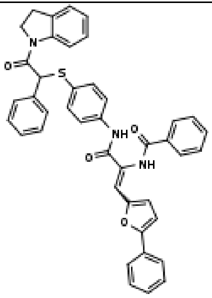 (2) | 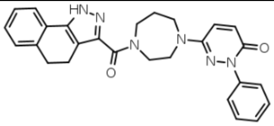 (1) | 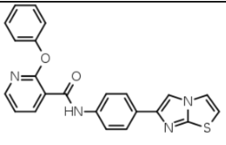 (1) | 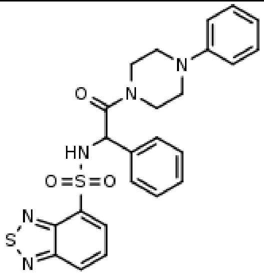 (1) |
| 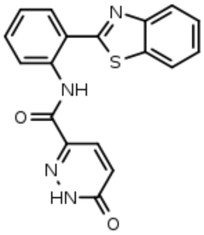 (1) | 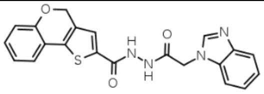 (1) | 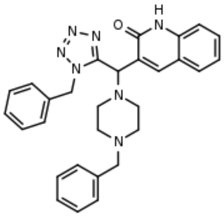 (1) | 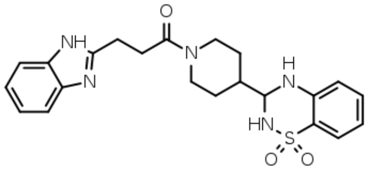 (1) |
| 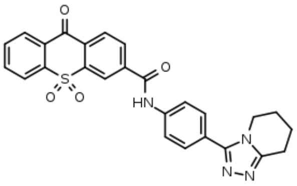 (1) | 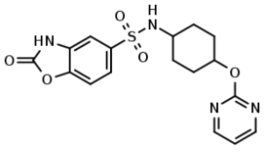 (1) | 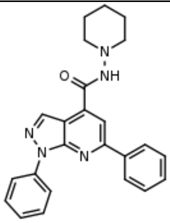 (1) | 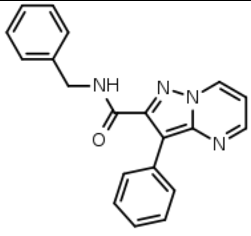 (1) |

| | | | |
|---|---|---|---|
| 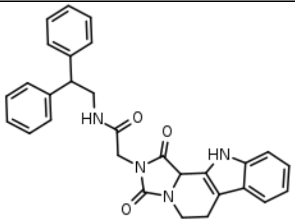 (1) | 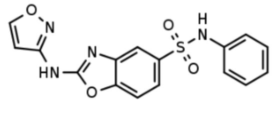 (1) | 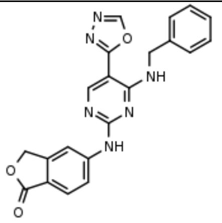 (1) | 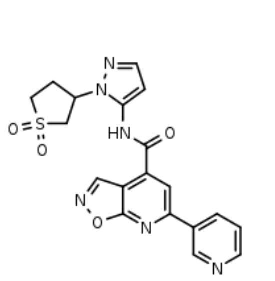 (1) |
| 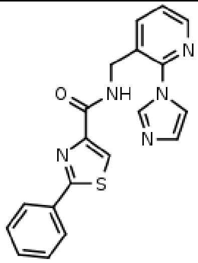 (1) | 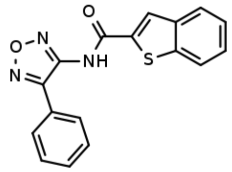 (1) | 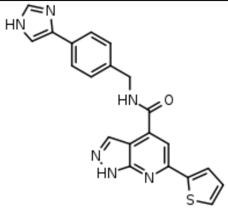 (1) | 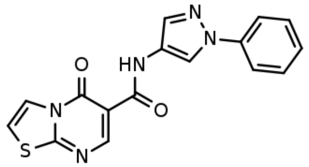 (1) |
| 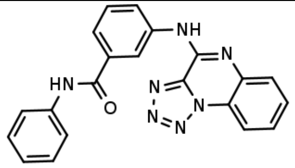 (1) | 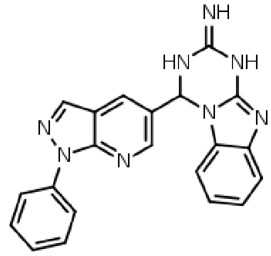 (1) | 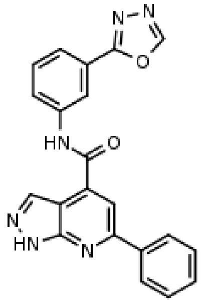 (1) | 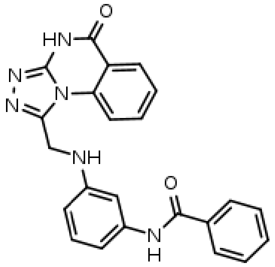 (1) |
| 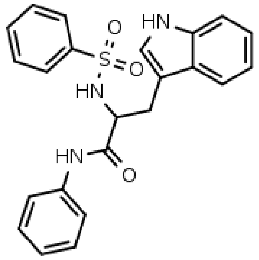 (1) | 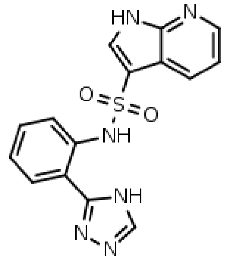 (1) | 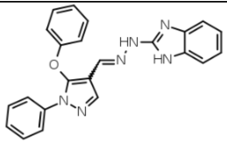 | 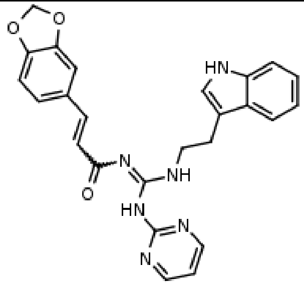 (1) |

| | | | |
|---|---|---|---|
| 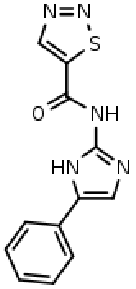<br>(1) | 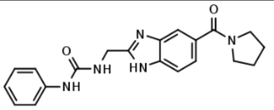<br>(1) | 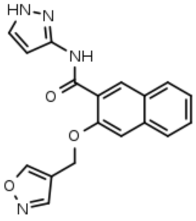<br>(1) | 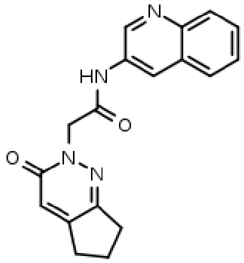<br>(1) |
| 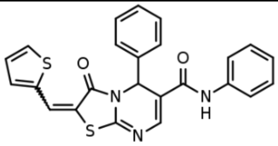<br>(1) | 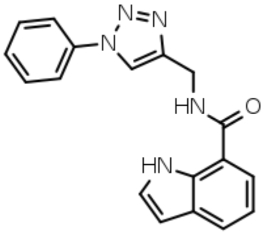<br>(1) | 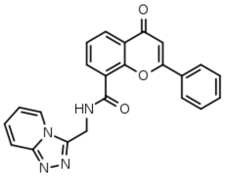<br>(1) | 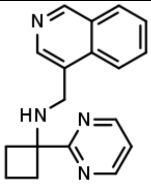<br>(1) |
| 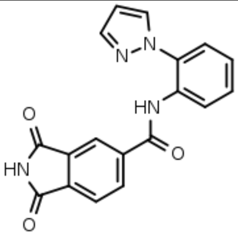<br>(1) | 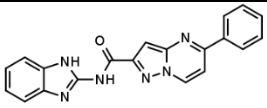<br>(1) | 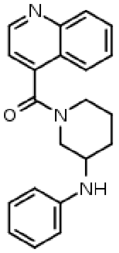<br>(1) | 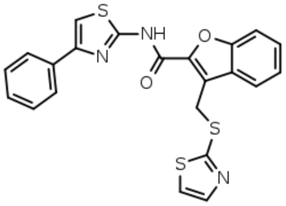<br>(1) |
| 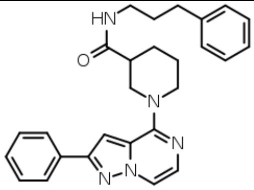<br>(1) | 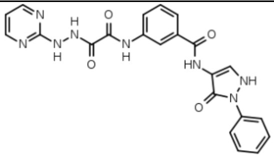<br>(1) | 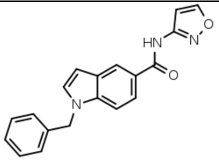<br>(1) | 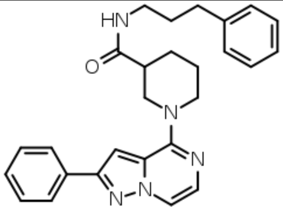<br>(1) |

**Table S2**. Murcko scaffolds extracted from the sub-parent library with 3,000 compounds showing uniqueness of ~80%. Representative structures from each group of scaffolds with different redundancy are shown along with the number of times each scaffold is observed in the library.

| Scaffold<br><br>(Redundancy) | Scaffold<br><br>(Redundancy = 1) | |
|---|---|---|
| 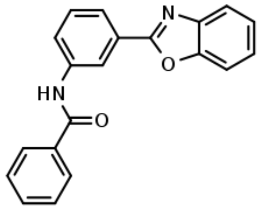 (14) | 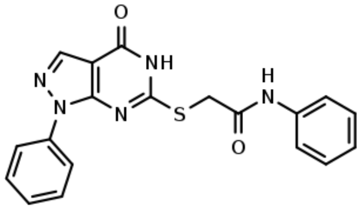 | 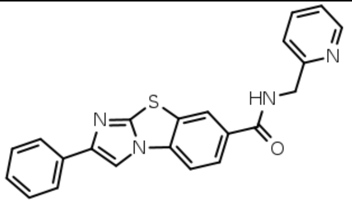 |
| 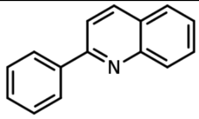 (12) | 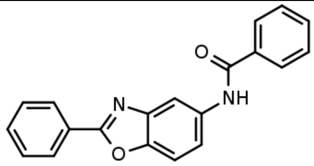 | 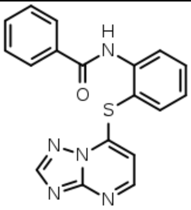 |
| 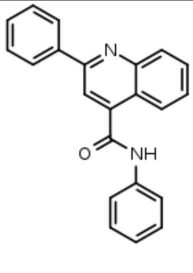 (11) | 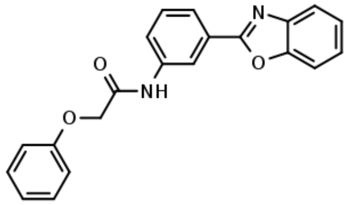 | 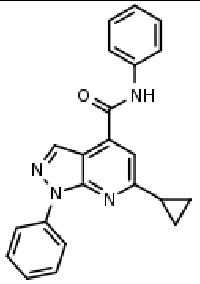 |
| 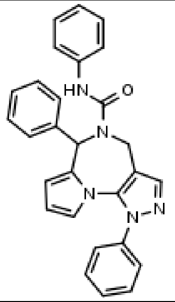 (10) | 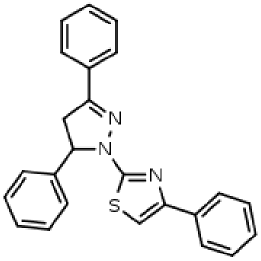 | 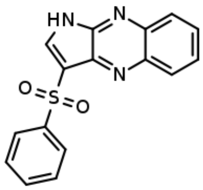 |

| | | |
|---|---|---|
| 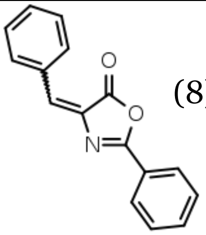 (8) | 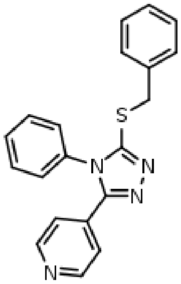 | 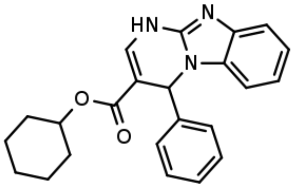 |
| 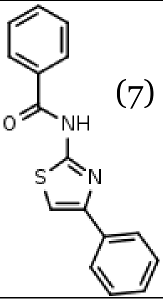 (7) | 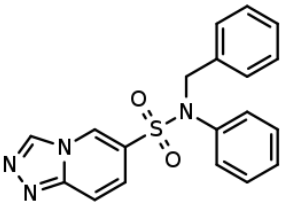 | 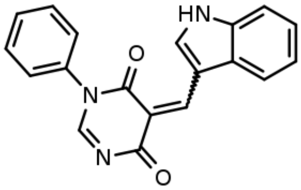 |
| 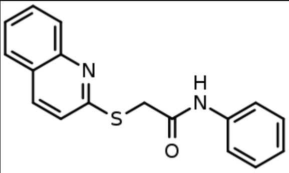 (6) | 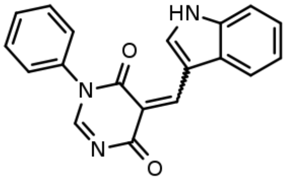 | 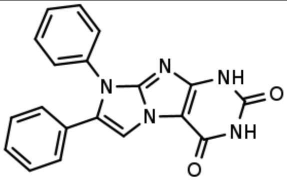 |
| 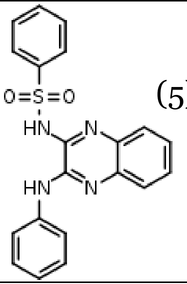 (5) | 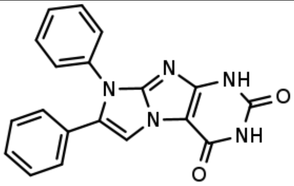 | 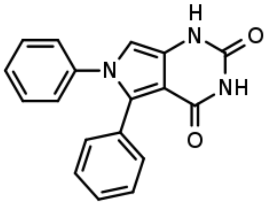 |
| 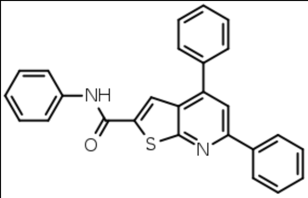 (4) | 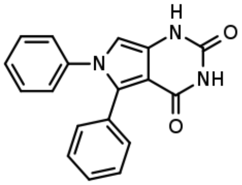 | 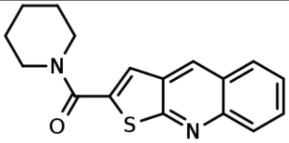 |
| 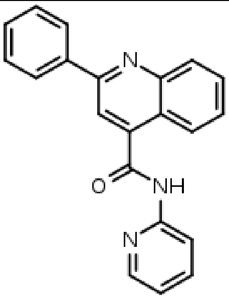 (3) | 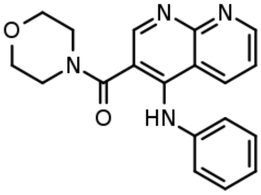 | 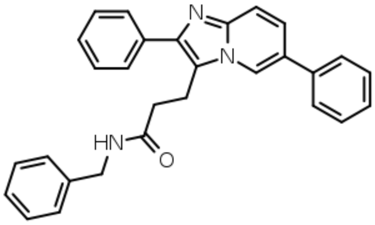 |

| **Table S3**. Physicochemical properties and quantitative estimate of drug-likeness (QED)[4] for compounds identified in the screen. | | | | | | | | |
|---|---|---|---|---|---|---|---|---|
| Compound ID | MW ($g{\cdot}mol^{-1}$) | ALOGP | HBA (count) | HBD (count) | PSA ($Å^2$) | ROTB (count) | AROM (count) | QED |
| 2 | 427.55 | 3.947 | 5 | 1 | 79.37 | 5 | 3 | 0.669 |
| 3 | 307.31 | 3.704 | 4 | 1 | 85.13 | 3 | 3 | 0.590 |
| 4 | 502.57 | 6.200 | 5 | 0 | 68.73 | 5 | 5 | 0.268 |
| 6 | 333.34 | 3.232 | 4 | 1 | 64.63 | 3 | 2 | 0.692 |
| 7 | 350.32 | 5.025 | 3 | 1 | 55.13 | 3 | 4 | 0.564 |
| 8 | 526.57 | 4.531 | 7 | 2 | 117.70 | 7 | 4 | 0.268 |
| 9 | 482.53 | 3.464 | 8 | 3 | 120.94 | 4 | 4 | 0.231 |
| 10 | 518.99 | 6.016 | 7 | 2 | 118.74 | 7 | 5 | 0.154 |
| 11 | 418.52 | 4.392 | 6 | 1 | 76.88 | 4 | 4 | 0.533 |
| 12 | 570.63 | 7.897 | 6 | 0 | 86.81 | 5 | 3 | 0.176 |
| 13 | 452.54 | 4.973 | 6 | 1 | 76.88 | 5 | 5 | 0.301 |
| 14 | 540.55 | 2.858 | 8 | 4 | 168.68 | 9 | 4 | 0.235 |
| 15 | 433.90 | 3.452 | 3 | 2 | 78.51 | 6 | 3 | 0.584 |
| 18 | 332.36 | 3.456 | 4 | 2 | 83.81 | 4 | 4 | 0.599 |
| 19 | 461.54 | 4.442 | 7 | 1 | 82.45 | 7 | 4 | 0.324 |
| 20 | 431.50 | 4.949 | 6 | 0 | 65.60 | 3 | 6 | 0.398 |
| 22 | 520.56 | 5.349 | 4 | 2 | 90.87 | 6 | 4 | 0.336 |
| MW: molecular weight; ALOGP: calculated octanol–water partition coefficient; HBA: hydrogen bond acceptors; HBD: hydrogen bond donors; PSA: polar surface area; ROTB: rotatable bonds; AROM: aromatic rings; QED: quantitative estimate of drug-likeness. | | | | | | | | |

| **Table S4: Sequences of primers used for PCR amplification, both qPCR and end-point PCR.** | | |
|---|---|---|
| **Primers** | **Forward Sequence ($5' \rightarrow 3'$)** | **Reverse Sequence ($5' \rightarrow 3'$)** |
| DMPK (RT-qPCR) | CGTGCAAGCGCCCAG | CTCCACCAACTTACTGTTTCATCCT |
| GAPDH (RT-qPCR) | AAGGTGAAGGTCGGAGTCAA | AATGAAGGGGTCATTGATGG |
| MAP3K4 (RT-qPCR) | CAATAAGCCTTACCTCAGCCTTG | GTTAAGCCAGAAACCAGACGTA |
| MAP4K4_ex22a (RT-PCR) | CCTCATCCAGTGAGGAGTCG | ATCACAGGAAAATCCCACCA |
| MBNL1_ex5 (RT-qPCR) | CTCAGTCGGCTGTCAAATCA | AGAGCAGGCCTCTTTGGTAA |
| MBNL1_ex5 (RT-PCR) | GCTGCCCAATACCAGGTCAAC | TGGTGGGAGAAATGCTGTATGC |
| Total MBNL1 (RT-qPCR) | CCGTTGCTCCAGGGAGAA | CACCAGGCATCATGGCATTG |
| MBNL1 exon 4-exon 6 (RT-qPCR) | CAGCTGCCATGGGAATTCCTCA | AGAGCAGGCCTCTTTGGTAA |
| MBNL1 exon 5 (RT-qPCR) | CTCAGTCGGCTGTCAAATCA | AGAGCAGGCCTCTTTGGTAA |
| MBNL2 exon 5 (RT-PCR) | ACAAGTGACAACACCGTAACCG | TTTGGTAAAGGATGAAGAGCACC |

| **Table S5. Sequences of RNA oligonucleotides used for single molecule studies with 5'-biotinylated DNA origami structures.** | |
|---|---|
| r(CUG)$_{21}$ RNA | 5′-biotin-UCGGCGAUCUACGCAGCGACAUAAUACUGCUGCUGCUGCUGCUGCUGCUGCUGCUGCUGCUGCUGCUGCUGCUGCUGCUGCUGCUGCUGCUGUAACAACCACUUCCUAAUCUGUCAUCUUCUG |
| 5′-DNA splint | GTCGCTGCGTAGATCGCCGAGAATCATAGATGGAGTGTGGTGTGCCTTGAAA |
| 3′-DNA splint | GTGTCTTTTGGTCTTTCTGGTGCTCTTCGAATCAGAAGATGACAGATTAGGAAGTGG |

**Two-Step K-means Clustering of a Large SMILES Library**

To efficiently select a chemically diverse subset from a large SMILES library, we implemented a two-step K-means clustering procedure as described below. All clustering was performed on molecular fingerprints derived from the input SMILES strings.

*Preprocessing and Feature Generation*

All input molecules were provided as SMILES strings in a single library file. SMILES were first standardized (e.g., canonicalization, removal of obvious salts or duplicates where appropriate) using RDKit. Each standardized SMILES was then converted to a fixed-length molecular fingerprint. In our implementation, we used circular (Morgan) fingerprints with a defined radius and bit length, but any RDKit-compatible fingerprint representation could be used.

The result of this step is a large matrix where each row corresponds to one molecule, and each column corresponds to a fingerprint bit (0/1). This matrix is the input for the clustering procedure.

Step 1: K-means Clustering on a Subset of the Library

Because clustering the full library directly is computationally expensive, we first applied K-means to a random subset of the molecules:

1. Subset selection

   A fixed number of molecules (e.g., 5,000 out of a 500,000 compound library) were randomly selected from the full set. Only these molecules and their fingerprints were used in the first clustering step.

2. K-means on the subset

   K-means clustering was run on the fingerprint matrix of this subset using a specified number of clusters ($K_1$). We used a standard K-means implementation (e.g., from scikit-learn) with K-means++ initialization and default convergence criteria.

3. Cluster center representatives

   After convergence, the algorithm returns:

   - A cluster assignment for each molecule in the subset.
   - A fingerprint vector representing the centroid (cluster center) of each cluster.

   For each cluster, we identified a "representative" molecule by finding the subset molecule whose fingerprint is closest to its corresponding cluster center (in Euclidean distance or another standard metric provided by the K-means implementation). These representative molecules are kept as the subset-level cluster representatives and their cluster centers are used in the second step.

Step 2: K-means Clustering on First-Step Cluster Centers

In the second step, we compress the set of first-step clusters into a smaller number of final clusters by running K-means directly on the cluster centers obtained from Step 1:

1. Preparing input for Step 2

   The fingerprint vectors corresponding to the $K_1$ cluster centers from Step 1 were assembled into a new matrix. Each row now represents a cluster center rather than an individual molecule.

2. K-means on cluster centers

   A second K-means run was performed on this matrix, using a smaller number of clusters ($K_2$), corresponding to the desired final number of chemically diverse groups (e.g., $K_2 = 100$). Again, we used K-means++ initialization and standard convergence parameters.

3. Mapping final centers back to molecules

   Once this second K-means run converged, each first-step cluster center was assigned to one of the $K_2$ final clusters. For each final cluster, we then selected a single molecule as the final representative:

   - Within each final cluster, we considered all first-step centers assigned to that cluster.
   - For each of those centers, we already had a representative molecule from Step 1.
   - We chose as the final representative the molecule whose associated first-step center was closest to the second-step cluster center.

This procedure yields one real molecule (a SMILES entry from the original library) for each final cluster. The resulting set of molecules constitutes a chemically diverse panel selected from the large starting library.

- **Fingerprint choice:** We used RDKit Morgan fingerprints with a defined radius and bit length (e.g., radius = 2, nBits = 1024), but other fingerprints could be substituted without changing the logic of the method.

- **Library size handling:** Only the subset is clustered in Step 1, which makes the approach scalable to very large libraries. Step 2 operates only on cluster centers, which are few ($K_1$), so it is computationally inexpensive.
- **Reproducibility:** Random seeds were fixed for both the subset selection and K-means initialization to ensure reproducible clustering and representative selection.
- **Output:** The final output of this two-step procedure is a list of SMILES corresponding to $K_2$ final cluster representatives, which can be used for follow-up calculations such as descriptor analysis, docking, or experimental screening.

**UMAP Analysis to Compare Chemical Space of Two SMILES Libraries**

To visualize and compare the chemical space occupied by two distinct SMILES libraries, we performed a joint Uniform Manifold Approximation and Projection (UMAP) analysis. The goal of this procedure was to embed both libraries into the same low-dimensional "chemical map" and assess the degree of overlap or separation between them.

*Input Libraries and Pre-processing*

Two sets of molecules were provided as separate SMILES lists (e.g., a "full library" and a "hit set," or a "library A" and "library B"). Each library was read from its own input file. We used the following preprocessing steps for both libraries:

1. *Standardization*

   SMILES strings from each library were standardized using RDKit.

   Standardization included:

- Parsing to RDKit Mol objects.
- Canonicalizing SMILES to a consistent representation.
- Optionally removing salts, handling disconnected fragments, and filtering out molecules that fail basic sanitization.

2. *Fingerprint generation*
3. Each standardized molecule was converted to a fixed-length molecular fingerprint using RDKit, typically a circular (Morgan) fingerprint with a defined radius and bit length (e.g., radius 2, 1024 bits). This produced a fingerprint matrix for each library, where each row corresponds to one molecule and each column corresponds to a fingerprint bit.

*Joint Embedding of Both Libraries with UMAP*

To ensure that both SMILES sets are embedded into the same UMAP space, we combined them before fitting the UMAP model:

1. *Concatenation of fingerprints*

   The fingerprint matrices of library A and library B were vertically concatenated into a single combined matrix. We also kept track of which rows belonged to which library (e.g., by storing indices or by assigning a label vector).

2. *UMAP model setup*

   We used the umap-learn Python package to perform dimensionality reduction. The UMAP model was configured with:

- A specified number of neighbors (e.g., n_neighbors=15–50, depending on the desired balance of local vs. global structure).
- A low-dimensional output space (typically 2D for plotting, n_components=2; 3D is optional).
- A suitable distance metric for fingerprints (for binary fingerprints, metrics such as “jaccard” or “cosine” are commonly used).
- A fixed random seed (random_state) to ensure reproducible embeddings.

3. *Fitting and transforming*

   The combined fingerprint matrix was passed to UMAP’s fit_transform method. This generated a low-dimensional embedding where each row corresponded to a single molecule from either library, now represented in 2D coordinates suitable for plotting.

4. *Splitting the embedding back into two sets*

   After embedding, we separated the resulting coordinates back into two groups using the stored indices or labels:

   - Coordinates corresponding to library A.
   - Coordinates corresponding to library B.

This allowed us to plot both libraries in the same UMAP space while preserving the identity of each molecule’s origin.

#### *Visualization and Chemical Space Overlap*

The final UMAP coordinates were visualized using standard plotting libraries (e.g., matplotlib):

1. **Scatter plot of the combined embedding**

   We generated a 2D scatter plot where each point represents one molecule. Points from library A and library B were displayed with different colors or markers. In some cases, transparency (alpha blending) was used to better visualize dense regions and overlaps.

2. **Visual inspection of overlap**

   By plotting both sets in the same UMAP space, we could visually assess:

   - Whether the two libraries occupy similar regions of chemical space.
   - Whether certain regions are dominated by one library or the other.
   - Whether hit compounds (if one set represents hits) cluster in specific zones relative to the full library.

3. **Optional enhancements**

   Depending on the analysis, we optionally:

   - Added contour lines or density estimates for each library to highlight regions of high point density.
   - Drew convex hulls or outlines around groups of points from each library to emphasize the extent of their chemical space.

*Practical Notes and Parameters*

- **Feature consistency:** Both libraries were processed with the same fingerprint settings (same radius, bit size, and preprocessing) so that UMAP operated on a consistent feature space.

- **Joint fitting:** The UMAP model was always fit on the combined dataset, rather than fitting separate models for each library. This ensures that the embedding is directly comparable across the two sets.
- **Reproducibility:** We fixed the UMAP random seed and logged all parameters (number of neighbors, minimum distance, distance metric, number of components) so that the same embedding can be regenerated if needed.
- **Scalability:** For very large libraries, we optionally subsampled the larger set (e.g., randomly selecting a fixed number of molecules) while keeping all molecules from a smaller set of interest, to keep the UMAP computation tractable.

**Principal Moments of Inertia (PMI) Shape Analysis**

Molecular shape analysis was performed using a custom Python workflow built with RDKit.[5] The goal of this analysis was to convert 2D SMILES input into 3D conformers, compute their principal moments of inertia, and position each compound within the established triangular PMI shape space to categorize molecules as rod-like, disc-like, or sphere-like.

*3D Structure Generation.* All input structures were provided as SMILES strings. RDKit was used to generate an initial 3D geometry for each compound. Hydrogen atoms were added, and an initial conformer was produced using the ETKDG algorithm (Experimental-Torsion Knowledge Distance Geometry), which combines distance-geometry embedding with torsion preferences derived from experimental crystallographic databases. Each embedded structure was subsequently subjected to energy minimization using the Universal Force Field (UFF). If the UFF optimization failed to converge, the lowest-energy geometry obtained during the minimization cycle was retained. This process yielded one low-energy 3D conformer per compound.

*Computation of Principal Moments of Inertia.* For each optimized conformer, RDKit's CalcPrincipalMomentsOfInertia function was used to obtain the three principal moments of inertia ($I_1 \le I_2 \le I_3$), which describe the mass distribution of the molecule in 3D space. These values were then converted into normalized PMI descriptors,

$$\mathrm{NPR1} = \frac{I_1}{I_3}, \mathrm{NPR2} = \frac{I_2}{I_3},$$

where $I_3$ is the largest principal moment. The resulting NPR1 and NPR2 values map each molecule to a point within the standard PMI triangle, where the three vertices correspond to idealized shape limits: *disc-like* (0,0), *rod-like* (0,1), and *sphere-like* (1,1).

*Shape Classification.* To assign categorical shape labels, each compound's (NPR1, NPR2) coordinate was compared to the coordinates of the PMI triangle vertices. Euclidean distance was used to determine which of the three idealized shapes—rod-like, disc-like, or sphere-like—was closest to each molecule. This nearest-vertex approach provides a reproducible, geometry-based metric for molecular shape classification that has been widely used in chemoinformatics and medicinal chemistry.

*Data Handling and Output.* For every molecule, the workflow recorded the SMILES string, 3D-generated conformer, NPR1 and NPR2 values, and the final shape classification. These results were exported to a CSV file for downstream analysis and comparison across screening libraries.

*PMI Triangle Visualization*. A PMI shape map was generated using matplotlib. The three edges of the PMI triangle were drawn to delineate the physically accessible shape space. Molecules were plotted at their (NPR1, NPR2) coordinates and colored according to their assigned shape class. This visualization highlights the distribution of compound shapes across the chemical library and provides an intuitive comparison of rod-like, disc-like, and sphere-like chemotypes. The resulting PMI triangle plot is shown in Figure X (panel E).

**Synthetic Methods**

**Abbreviations:** COMU, 1-Cyano-2-ethoxy-2-oxoethylidenaminooxy)dimethylamino-morpholino-carbenium hexafluorophosphate; DIPEA, N,N-diisopropylethylamine; DCM, dichloromethane; DMF, N,N-dimethylformamide; DMSO, dimethyl sulfoxide; TEA, triethylamine; EtOAc, ethyl acetate; $H_2O$, water; ESI-MS, electrospray ionization-mass spectrometry; $NaHCO_3$, sodium bicarbonate; $Na_2SO_4$, sodium sulfate; NMR, nuclear magnetic resonance; $SiO_2$, silica; TLC, thin layer chromatography.

**General.** All reagents and solvents used for chemical synthesis were purchased from standard suppliers and were used without further purification unless mentioned otherwise. Reactions were monitored with an Agilent 1260 Infinity LC system coupled to an Agilent 6230 TOF (HR-ESI) equipped with a Poroshell 120 EC-C18 column (Agilent, 50 mm x 4.6 mm, 2.7 µm) or by TLC. Products were purified by Isolera One flash chromatography system (Biotage) using pre-packed silica irregular 40-60 µm 60A column (Claricep Flash, Agela Technologies). NMR spectra for compound characterization were measured on measured by a 400 UltraShieldTM (Bruker) (400 MHz for $^1H$ and 100 MHz for $^{13}C$) or AscendTM 600 (Bruker) (600 MHz for 1H and 150 MHz for $^{13}C$). Chemical shifts are expressed in ppm relative to trimethylsilane (TMS) for $^1H$ and residual solvent for $^{13}C$ as internal standards. Coupling constant (J values) are reported in Hz.

**Synthesis of N-(4-(2,3-dihydro-1H-inden-5-yl)thiazol-2-yl)-2 (methylamino)benzamide**

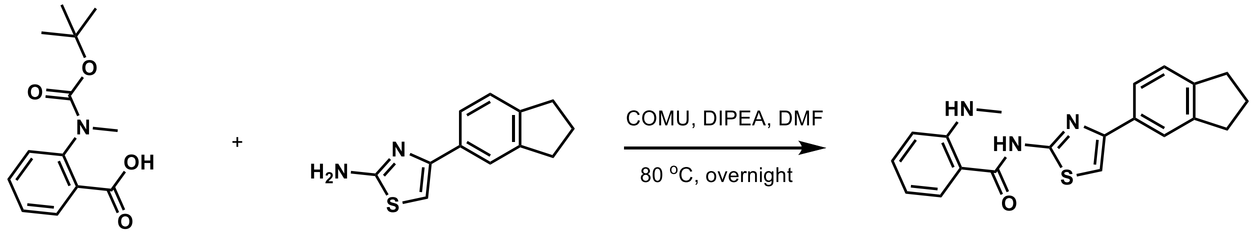


N-Boc-N-methylanthranilic acid (0.69 mmol, 174 mg) (Santa Cruz, sc-263820A) and 4-(2,3-dihydro-1H-inden-5-yl)-1,3-thiazol-2-amine (0.46 mmol, 100 mg) (Enamine, EN300-10998) were dissolved in anhydrous DMF. To the solution was added COMU (0.69 mmol, 297 mg), oxyma (0.69 mmol, 98.5 mg) and DIPEA (0.69 mmol, 120 µL). The reaction was heated at 80 °C overnight and monitored by TLC with (5% ethyl acetate in hexane). The crude product was washed with saturated $NaHCO_3$ three times, extracted with ethyl acetate and dried over $Na_2SO_4$. Solvent was evaporated in vacuo and the mixture was further purified by silica gel at 5% ethyl acetate in hexane and yield as pale-white solid (60%). ESI-MS $[M+H]^+$ =350.4; Found: 350.3611. $^1$H NMR (600 MHz, DMSO-$d_6$) δ (ppm) 12.34 (s,1H), 7.94-7.92 (d, J = 6 Hz, 1H), 7.8 (s, 1H), 7.72-7.71 (d, J = 6 Hz, 1H), 7.54 (s, 1H), 7.41-7.38 (m, J = 6 Hz, 1H), 7.28-7.27 (d, J = 6 Hz, 1H), 6.73-6.72 (d, J = 6 Hz, 1H), 6.63-6.61 (m, J = 6 Hz, 1H), 2.92-2.86 (m, J = 12 Hz, 4H), 2.84 (s, 3H), 2.06-2.01 (m, J = 6 Hz, 2H); $^{13}$C NMR (125 MHz, DMSO-$d_6$) δ (ppm) 167.49, 158.17, 150.99, 149.59, 144.31, 143.51, 134.20, 132.62, 129.56, 124.47, 124.02, 121.75, 114.23, 111.97, 111.12, 107.29, 32.34, 32.19, 29.46, 25.12.

**Synthesis of N-(4-(2,3-dihydro-1H-inden-5-yl)thiazol-2-yl)-2-(N-methylmethylsulfonamido) benzamide (2).**

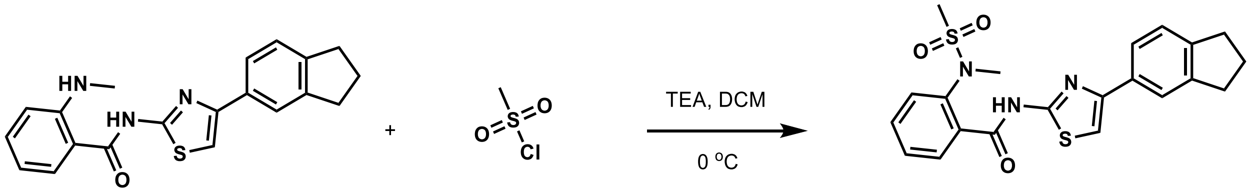


N-(4-(2,3-dihydro-1H-inden-5-yl)thiazol-2-yl)-2-(methylamino)benzamide (0.02 mmol, 10 mg) was dissolved into dry DCM and stirred in ice bath. To the solution was added 1 equivalent of Methanesulfonyl chloride (0.02 mmol, 2 µL) and triethyl amine (0.04 mmol, 25 µL). The mixture was stirred at 0 °C for one hour then, slowly warmed to room temperature and was allow to continue to react overnight. The reaction mixture was evaporated under reduced pressure, and the product was extracted using DCM (25 mL×2). The organic phase was washed with sat. $NaHCO_3$ (20 mL×2) and dried over $Na_2SO_4$. The solvent was then evaporated under reduced pressure to give a pale-yellow solid. ESI-MS $[M+H]^+$ =428.54; Found: 428.3583. $^1$H NMR (600 MHz, DMSO-$d_6$) δ (ppm) 12.46 (s,1H), 7.78 (s, 1H), 7.70-7.67 (m, d = 6 Hz, 2H), 7.63-7.62 (m, d=6 Hz, 2H), 7.58 (s, 1H), 7.51-7.48 (m, J = 6 Hz, 1H), 7.28-7.26 (d, d = 12 Hz, 1), 3.29 (3, 3H), 2.99 (s, 3H), 2.92-2.86 (m, d = 12 Hz, 4H), 2.07-2.02 (m, d = 12 Hz, 2H); $^{13}$C NMR (125 MHz, DMSO-$d_6$) δ (ppm) 165.56, 157.75, 149.53, 144.23, 143.48, 139.27, 134.76, 132.49, 131.83, 129.44, 128.05, 127.91, 124.43, 123.97, 121.67, 107.24, 45.77, 38.63, 36.34, 32.28, 32.12, 25.05, 8.66.

**[1]H-NMR characterization of compound N-(4-(2,3-dihydro-1H-inden-5-yl)thiazol-2-yl)-2-(methylamino)benzamide.**

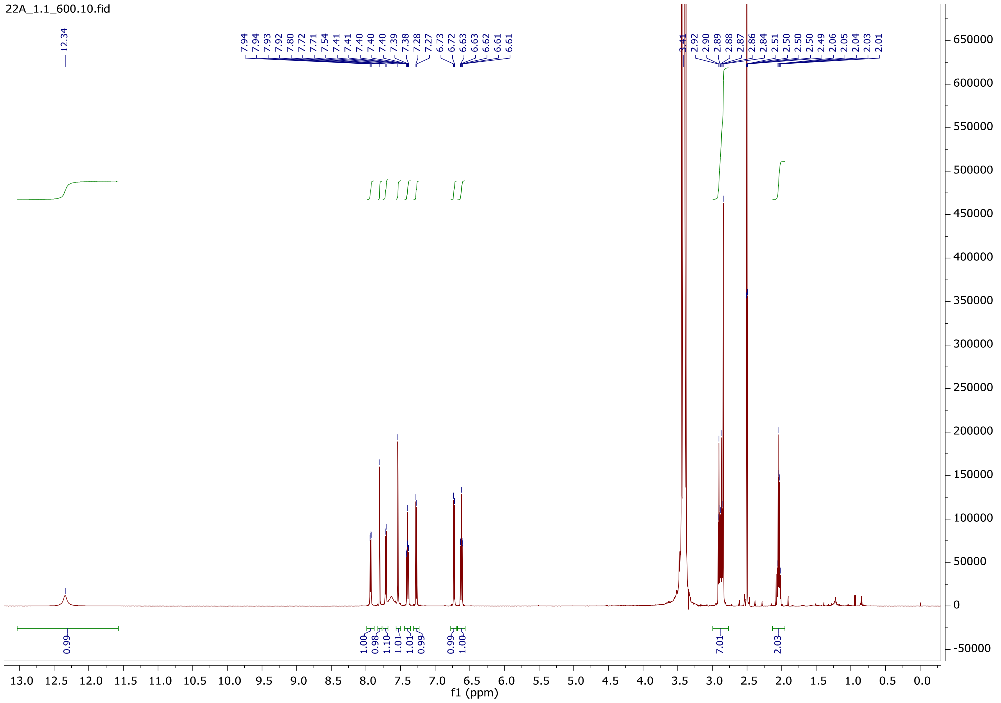


**[13]C-NMR characterization of compound N-(4-(2,3-dihydro-1H-inden-5-yl)thiazol-2-yl)-2-(methylamino)benzamide.**

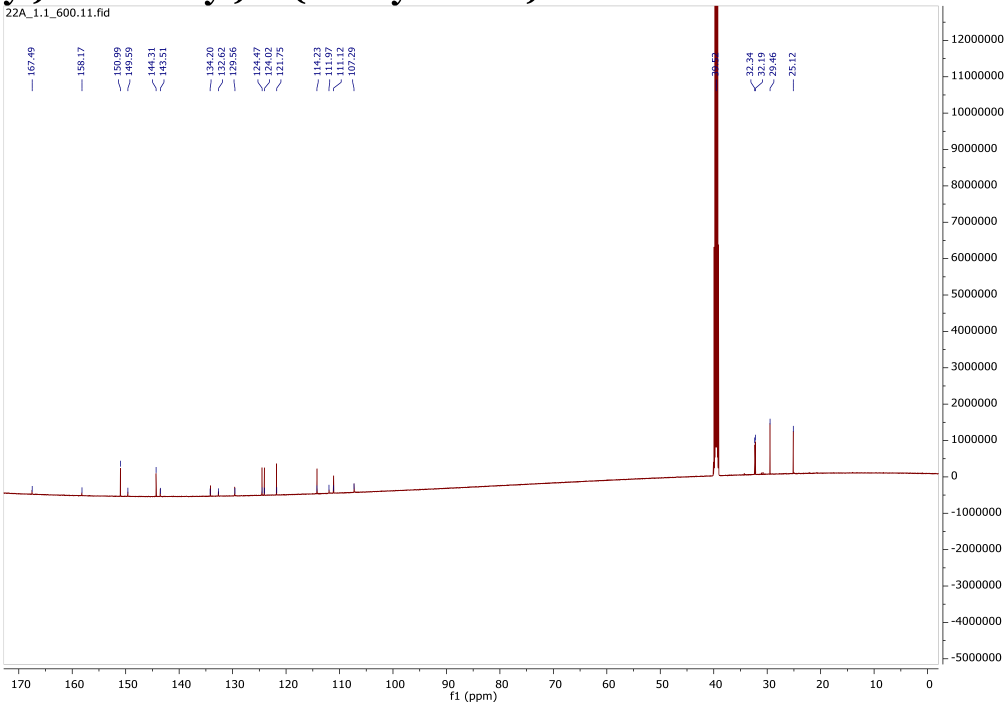

**¹H-NMR characterization of compound N-(4-(2,3-dihydro-1H-inden-5-yl)thiazol-2-yl)-2-(N-methylmethylsulfonamido)benzamide. (Compound 2).**

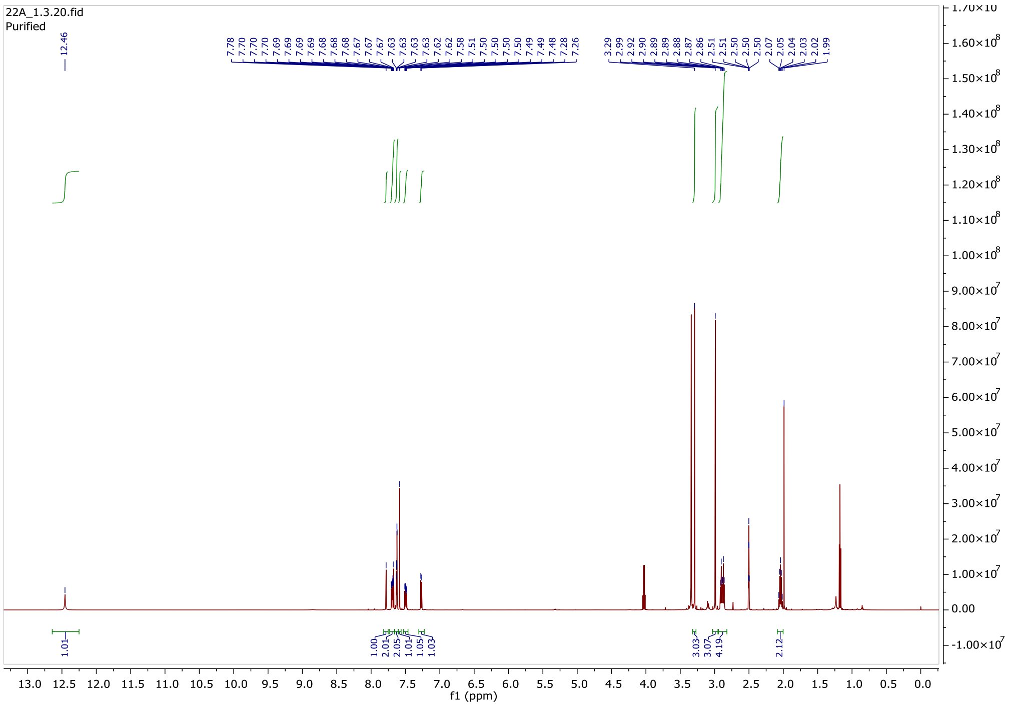

**¹³C-NMR characterization of compound N-(4-(2,3-dihydro-1H-inden-5-yl)thiazol-2-yl)-2-(N-methylmethylsulfonamido)benzamide. (Compound 2).**

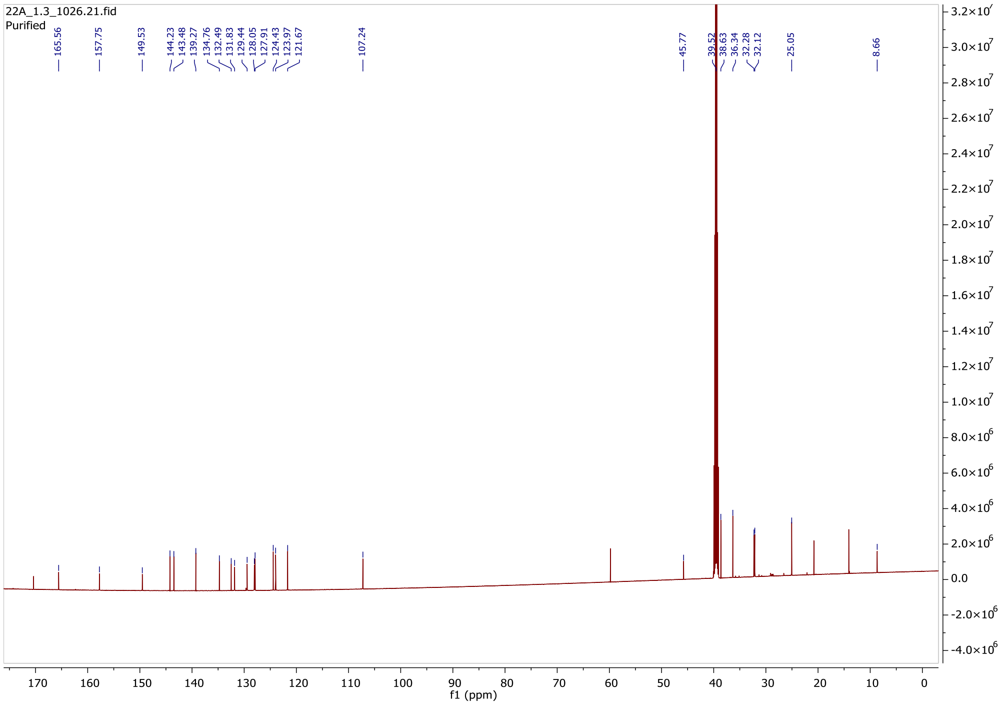
